# Programmable domestication of thermophilic bacteria through removal of non-canonical defense systems

**DOI:** 10.64898/2026.03.21.713436

**Authors:** Jae-Yoon Sung, Mun Hoe Lee, Jungsoo Park, Hyungbin Kim, Dariimaa Ganbat, Donggyu Kim, Hyun-Woo Cho, Min Kuk Suh, Jung-Sook Lee, Sang Jae Lee, Seong Bo Kim, Dong-Woo Lee

## Abstract

Thermophilic bacteria offer major advantages for industrial biotechnology, yet most remain genetically intractable because cellular defense systems block efficient DNA acquisition. Here, we present a programmable domestication strategy that converts wild *Geobacillus* strains into genetically tractable thermophilic hosts. We developed the Domestication of Non-Model Bacteria (DNMB) Suite, a multi- omics-guided computational framework that systematically identifies genetic barriers to transformation. DNMB analysis revealed that non-canonical nuclease-based defense systems, including Wadjet II, constitute dominant barriers to DNA uptake in previously intractable *Geobacillus* strains. Targeted deletion of these loci increased transformation efficiency by up to six orders of magnitude. We further established a hierarchical thermophilic engineering toolkit that integrates plasmid artificial modification, conjugation-assisted DNA delivery, and genome editing using an endogenous CRISPR-Cas9 system. The resulting domesticated strains support stable heterologous expression and tunable genetic control at elevated temperatures. Together, these results establish a generalizable framework for transforming genetically intractable thermophiles into programmable industrial chassis.

## Introduction

Microbial platform organisms play an important role in modern biotechnology, enabling the sustainable production of fuels, chemicals, pharmaceuticals, enzymes, and food ingredients ^1, 2^. Traditionally, microbial cell factories have relied on mesophilic hosts such as *Escherichia coli*, *Bacillus subtilis*, and *Saccharomyces cerevisiae*, because of their genetic tractability, extensive molecular toolkits, and well-characterized physiology ^3–5^. However, these organisms often encounter limitations under industrial conditions, including contamination risks, restricted substrate solubility, and reduced enzyme stability at elevated temperatures ^6, 7^. Thermophilic microorganisms offer a compelling alternative, as elevated process temperatures can accelerate reaction kinetics, increase substrate and product solubility, and reduce contamination during large-scale bioprocessing ^8–10^. Consequently, there has been growing interest in developing thermophiles as next-generation industrial hosts ^11^.

Despite these advantages, most thermophilic bacteria remain genetically intractable. Effective genetic engineering requires efficient DNA delivery, stable plasmid maintenance, and predictable gene expression, yet these capabilities are poorly developed in many thermophiles ^12^. Among thermophilic genera, *Geobacillus* species are particularly promising due to their rapid growth, environmental resilience, and natural ability to secrete industrially relevant hydrolases such as proteases, lipases, and cellulases ^13–16^. Notably, *G. stearothermophilus* and *G. thermodenitrificans* are designated as Qualified Presumption of Safety (QPS) and Generally Recognized as Safe (GRAS) organisms, making them attractive candidates for food-grade thermophilic bioproduction ^17^. Nevertheless, domestication of wild-type *Geobacillus* strains has been hindered by the scarcity of versatile genetic tools and by substantial strain-to-strain variability in both genotype and phenotype ^18–20^. As a result, engineering efforts have largely been confined to a few tractable species, including *G. kaustophilus, G. thermodenitrificans,* and *Parageobacillus thermoglucosidasius* ^21^.

Restriction–modification (R–M) systems have long been recognized as major barriers to DNA transformation in bacteria ^22^. Recent advances in single-molecule real-time (SMRT) sequencing and methylome analysis have revealed extensive diversity in DNA modification patterns and highly strain- specific defense architectures ^23^. However, these approaches often fail to detect less-characterized non- canonical defense systems that can impose equally strong, or even dominant, constraints on genetic manipulation. Such cryptic defense modules are frequently embedded within defense islands associated with mobile genetic elements but remain poorly annotated in many thermophilic genomes, thereby hindering systematic engineering of non-model microorganisms ^24^.

Here, we show that cryptic non-canonical defense systems, including a Wadjet II-like complex, Gabija, and pAgo modules, constitute major barriers to horizontal DNA acquisition in wild-type *G. stearothermophilus* strains EF60045 and SJEF4-2 ^25^. These loci have largely escaped conventional genome annotation and are refractory to traditional plasmid artificial modification strategies that mimic host DNA methylation patterns ^26^. To systematically uncover and dismantle these barriers, we developed the Domestication of Non-Model Bacteria (DNMB) Suite, a modular multi-omics framework integrating comparative genomics, transcriptomics, and motif-based defense mining. DNMB enabled precise mapping of defense islands, comprehensive functional annotation, and rational host redesign, including codon optimization, ribosome binding site engineering, and construction of *Geobacillus*-specific *E. coli*- shuttle vectors with native replication origins and tunable promoters. We further engineered an endogenous CRISPR-Cas9 system (GeoCas9EF) and incorporated it into a plasmid artificial modification-assisted conjugative engineering (PACE) workflow to enable efficient genome editing ^22, 27^. Systematic locus-by-locus removal of defense modules, rather than canonical R–M loci, resulted in transformation efficiency increases of up to six orders of magnitude and markedly improved genetic stability. To demonstrate the utility of this domesticated thermophilic chassis, we reprogrammed *G. stearothermophilus* for D-tagatose metabolism and established a growth-coupled screening platform for rare-sugar isomerases. Collectively, these results identify cryptic defense systems as dominant barriers to thermophile domestication and establish *G. stearothermophilus* as a genetically stable, programmable chassis for high-temperature industrial biotechnology.

## Results

### The DNMB Suite: A modular computational framework for deconstructing domestication barriers in non-model bacteria

Domestication of wild thermophilic bacteria is frequently impeded by cryptic genetic barriers, including incomplete genome annotations, poorly characterized defense systems, and the lack of predictive tools for assessing engineering tractability. To systematically identify these constraints, we developed the DNMB Suite, a modular R-based computational framework that integrates comparative genomics and functional annotation to uncover genetic determinants limiting DNA uptake and heterologous expression (**Fig. 1a**).

**Figure 1.**
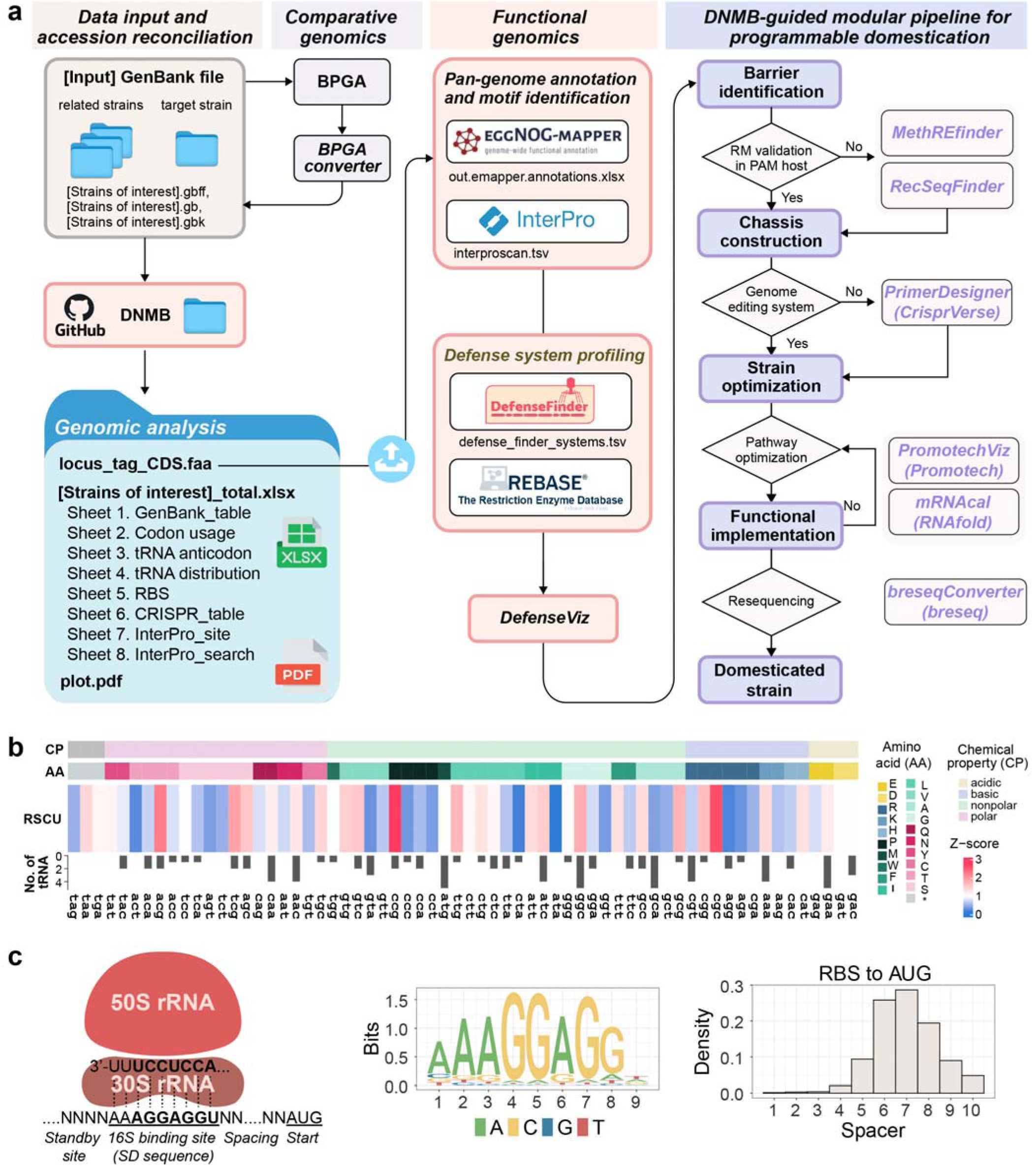
DNMB pipeline for programmable domestication of non-model bacteria. The DNMB (Domestication of Non-Model Bacteria) Suite integrates comparative genomics, functional annotation, and translational analysis to identify strain-specific barriers to genetic accessibility and guide rational chassis engineering. **a,** Overview of the DNMB workflow. Genomic inputs (GenBank format) from target and related strains are processed through comparative genome analysis (BPGA) and functional annotation (eggNOG-mapper, InterProScan). Defense systems are detected using DefenseFinder and REBASE and visualized with DefenseViz. Integrated outputs identify candidate domestication barriers, including restriction-modification (R–M) systems, anti-conjugation loci, CRISPR–Cas elements, and other defense modules, and inform downstream engineering steps such as barrier removal, genome editing system establishment, chassis optimization, and pathway implementation. **b,** Comparative analysis of codon usage and translation capacity in *G. stearothermophilus*. Heat maps show relative synonymous codon usage (RSCU), predicted tRNA gene copy numbers, and amino-acid classifications grouped by physicochemical properties. **c,** Genomically inferred ribosome binding site (RBS) architecture. Left, schematic of 30S ribosome interaction with the Shine-Dalgarno sequence. Middle, sequence logo of the consensus RBS motif identified across *Geobacillus* genomes. Right, distribution of spacer lengths between the RBS and start codon (AUG), indicating a preferred spacing of approximately 5–8 nucleotides.

The DNMB pipeline converts GenBank files into structured annotation tables and enriches gene- and protein-level features using eggNOG-mapper and InterProScan. A key component of the framework is a gene/protein identifier harmonization module that reconciles inconsistencies in gene symbols, protein names, and accession identifiers across databases, enabling reliable cross-genome comparisons. This normalized step proved essential for accurately tracking homologous loci and defense modules across diverse strains.

To resolve strain-specific defense architectures, DNMB integrates multiple layers of genomic layers, including gene neighborhood organization, operon structure, motif distribution, and methylation-related features. Candidate defense islands were identified using DefenseFinder and REBASE and visualized with DefenseViz (**Fig. 1a; DataSets 1–2**). This analysis revealed a diverse combination of restriction– modification (R–M) systems, anti-conjugation loci, CRISPR–Cas elements, and additional defense modules across *Geobacillus stearothermophilus* isolates, indicating substantial heterogeneity in genetic accessibility barriers.

Beyond defense profiling, DNMB evaluates translational constraints affecting heterologous gene expression. Codon usage was quantified using complementary metrics including effective codon usage, frequency, and relative synonymous codon usage (RSCU), and compared with predicted tRNA gene copy numbers and amino-acid composition to identify potential translational bottlenecks (**Fig. 1b**). An ribosomal binding site (RBS)-scanning module was further implemented to detect Shine-Dalgarno sequences by aligning the reverse complements of the 3’ terminus of 16S rRNA to upstream coding regions. Application of this module identified a conserved consensus motif (AAGGGAGGN) and a preferred ∼7-nt spacing between the RBS and start codon in *Geobacillus*, consistent with efficient native translation initiation (**Fig. 1c**).

Together, these analyses systematically dissect genetic barriers to engineering by integrating annotation normalization, defense system detection, regulatory motif discovery, and translational optimization within a unified framework. The DNMB Suite enabled prioritization of strain-specific domestication targets and directly guided subsequent engineering of genetically tractable *G. stearothermophilus* hosts described below.

### Comparative genomic and methylomic analyses reveal strain-specific defense architectures in *Geobacillus*

To systematically investigate barriers to genetic transformation in thermophilic *Geobacillus*, we applied the DNMB Suite to a representative dataset of 51 complete genomes spanning diverse *Geobacillus* and *Parageobacillus* species, including the hot spring isolates EF60045 and SJEF4-2, the type strain *G. stearothermophilus* ATCC 12980, and additional environmental and industrial strains (**DataSet 3**).

Comparative genomic analysis revealed extensive inter-strain variability in both canonical and non- canonical phage defense systems, including R–M systems, CRISPR-Cas modules, and diverse antiphage immunity loci (**Fig. S1a**). Pan-genome reconstruction across 157,210 predicted genes identified only 1,217 core genes shared among all strains (≥ 40% identity), indicating substantial genome plasticity and expansion of the accessory genome driven by ecological diversification (**DataSet 3**). Notably, many defense loci were embedded within strain-specific genomic islands enriched for mobile genetic elements and prophage remnants, suggesting that phage-mediated horizontal gene transfer has played a major role in shaping defense system diversity.

Strains known to be genetically recalcitrant, including EF60045 and SJEF4-2 ^25^, contained particularly large defense islands densely populated with R–M systems, orphan methyltransferases, CRISPR arrays, and multiple non-canonical defense modules, including Wadjet, cyclic oligonucleotide-based antiphage signaling systems (CBASS), and prokaryotic Argonaute (pAgo) systems (**Fig. 1**).

**Figure 2.**
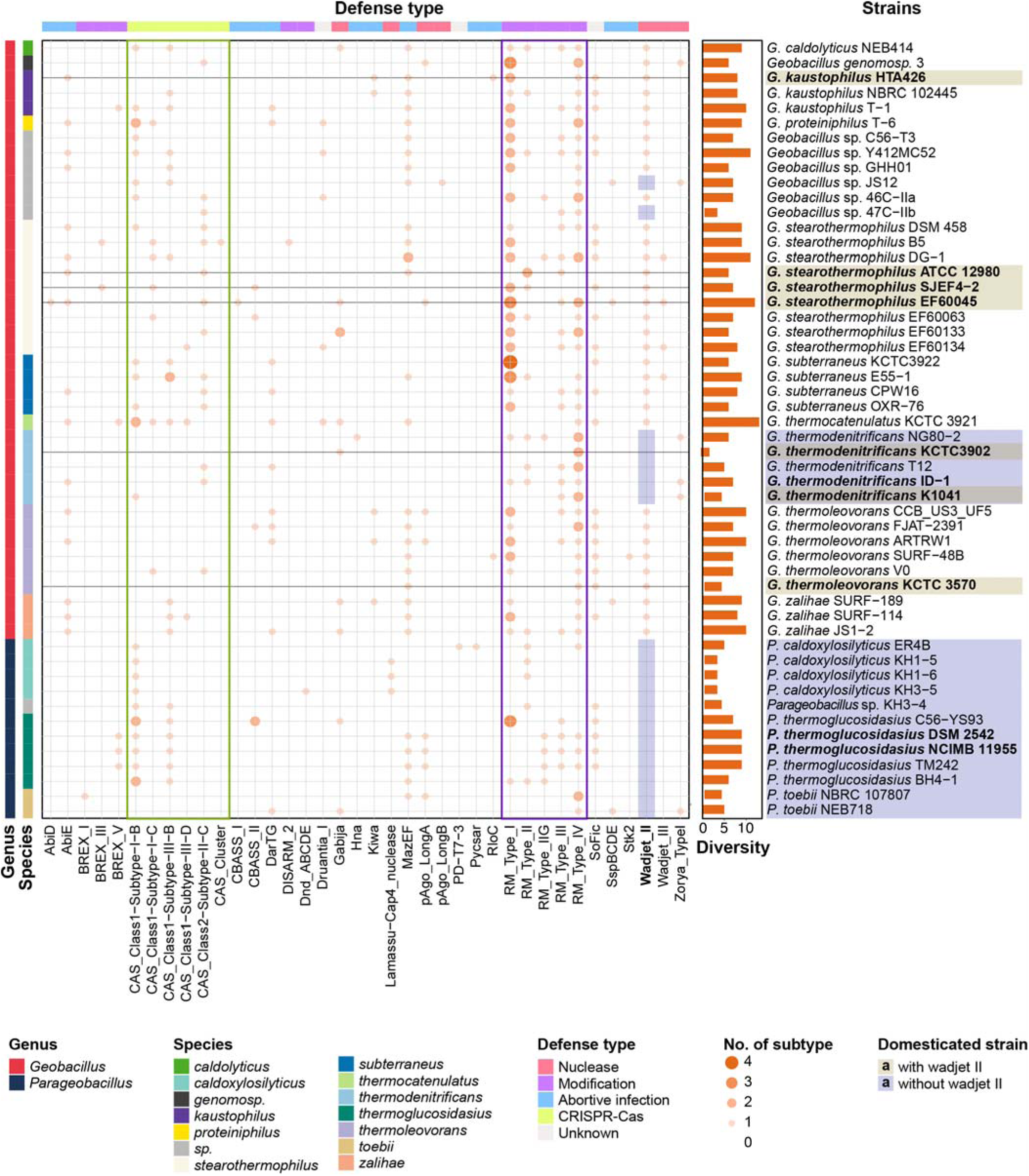
Comparative genomic distribution of defense systems across (*Para*)*Geobacillus* genomes. Bubble heatmap showing the distribution of predicted bacterial defense systems across 51 complete genomes of *Geobacillus* and *Parageobacillus*. Columns represent defense system subtypes grouped by functional category, including R–M, CRISPR–Cas, abortive infection systems, and additional defense modules such as BREX, DISARM, Wadjet, and Dnd. Rows correspond to individual strains. Circle size indicates the copy number of each subtype, whereas color denotes functional category (nuclease, modification, abortive infection, CRISPR–Cas, or unknown). Colored side bars indicate genus and species classification. The bar plot on the right summarizes overall defense diversity per strain, defined as the number of distinct defense subtypes detected in each genome. Strains analyzed or domesticated in this study are highlighted, and the presence or absence of the Wadjet II system is indicated.

Phylogenomic analysis further showed that these strains occupy distinct evolutionary clades compared with transformation-permissive species such as *G. thermodenitrificans* and *P. thermoglucosidasius*, suggesting that the accumulation of extensive defense repertoires reflects lineage-specific adaptation to viral predation and long-term retention of complex immunity modules.

Using DefenseFinder and REBASE, we annotated substantial diversity in R–M system architectures across the dataset (**DataSets 3-4**). For example, EF60045 encoded seven complete R–M systems (three Type I, one Type II, one Type III, and two Type IV), whereas SJEF4-2 carried a distinct combination of two Type I systems with intact HsdS specificity subunits, one Type III system, and one Type IV system (**Fig. S1b**). PacBio SMRT sequencing confirmed activity of these systems, identifying four and three active methylation motifs in EF60045 and SJEF4-2, respectively (**Tables S1 and S2**). In addition, a conserved thermostable Type III methyltransferase present across the *Geobacillus spp*. (GSJ10_06490, recognizing the GCC^6m^AT motif) was purified and characterized by differential scanning calorimetry (DSC), revealing melting transitions at 69.2°C and 77.5°C, consistent with structural adaptation to high-temperature environments (**Fig. S1c**). This enzyme was subsequently used as a universal methyltransferase to introduce GCC^6m^AT methylation during preparation of plasmid artificial modification donor hosts.

Beyond canonical restriction systems, DNMB-based defense profiling identified a broad diversity of non-canonical antiphage defense modules, including Wadjet-II, bacteriophage exclusion (BREX), Gabija, SoFic, SspBCDE, CBASS, and pAgo systems (**Fig. 1**). Individual genomes encoded between 2 and 13 distinct defense subtypes, indicating that thermophilic *Geobacillus* species possess highly modular and polygenic immune architectures. Strikingly, strains lacking Wadjet-II, including *P. thermoglucosidasius* and *G. thermodenitrificans*, generally exhibited reduced defense burdens and correspondingly higher genetic tractability, whereas undomesticated strains harboring complete Wadjet- II operons, such as EF60045 and SJEF4-2, showed dense clusters of immunity loci.

Collectively, these results demonstrate that defense system architecture in (*Para*)*Geobacillus* is highly variable, modular, and strongly strain-specific, and suggest that the cumulative burden of both canonical and non-canonical defense systems constitutes a major barrier to genetic accessibility. By integrating comparative genomics, methylome profiling, and functional validation, the DNMB Suite provides a predictive framework for identifying candidate defense loci that limit transformation and for rationally prioritizing targets for genome engineering in thermophilic chassis strains.

### Host-matched methylation improves but is insufficient for the transformation of wild *Geobacillus*

Developing genetically tractable thermophilic *Geobacillus* strains requires efficient DNA delivery, which remains a major bottleneck for domestication of wild-type isolates. Guided by DNMB-based comparative genomics and methylome analyses, we first sought to overcome R–M barriers by establishing a plasmid artificial modification system that reproduces host-specific methylation patterns during plasmid preparation in *Escherichia coli* (**Fig. 2a**).

**Figure 3.**
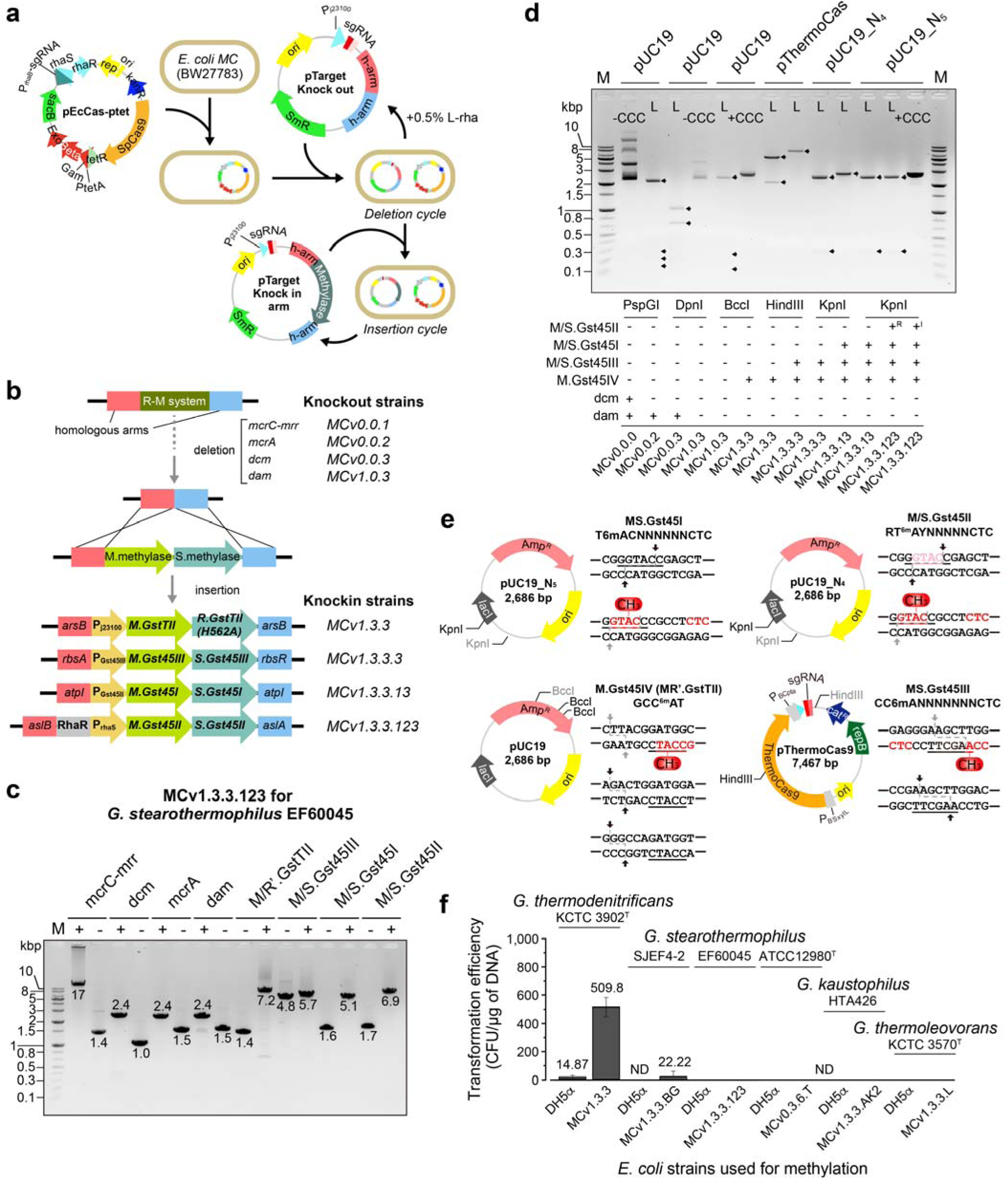
Strain-specific methylome mimicry and plasmid artificial modification for DNA delivery in *Geobacillus*. a, Strategy for constructing plasmid artificial modification hosts in *Escherichia coli* (MCv series). A minicircle-derived background was engineered by iterative CRISPR-assisted genome editing to remove endogenous restriction systems and install heterologous methyltransferase modules that reproduce *Geobacillus*-specific methylation patterns during plasmid propagation. **b**, Modular engineering scheme of plasmid artificial modification hosts. Native *E. coli* restriction and methylation genes (*mcr*, *mrr*, *dam*, and *dcm*) were sequentially deleted, followed by chromosomal integration of *Geobacillus*-derived methyltransferases at neutral loci (e.g., *arsB*, *rbsA*, *atpI*, and *asl*) under inducible promoters to reconstruct strain-specific methylomes. Representative knockout (MCv0.0.x–MCv1.0.3) and knock-in (MCv1.3.3 series) lineages are shown. **c,** PCR validation of deletion and methyltransferase knock-in events in a representative plasmid artificial modification host (MCv1.3.3.123) matched to *G. stearothermophilus* EF60045. Expected fragment sizes corresponding to wild-type, deletion (-), and insertion (+) alleles are indicated. **d,** Methylation-sensitive restriction digestion assays confirming plasmid methylation by engineered hosts. Plasmids isolated from different MC variants exhibit distinct digestion patterns depending on installed methyltransferase modules. DNA topology states are indicated: covalently closed circular (CCC) and linear (L). +^R^, repression; +^I^, induction. **e,** Representative plasmid constructs and cognate methylation motifs used to validate *Geobacillus*-derived methyltransferases, including Type I and Type III systems and their protected recognition sequences. **f,** Transformation efficiencies of methylome-matched plasmid DNA in multiple *Geobacillus* strains. Reporter plasmids prepared in plasmid artificial modification hosts exhibit increased transformation efficiency relative to DNA prepared from standard *E. coli* strains. Efficiency is expressed as CFU µg ¹ DNA. ND, not detected.

To construct methylation-compatible donor strains, endogenous *E. coli* restriction (*mcrA*, *mcrBC*, *mrr,* and *hsdR*) and methylation (*dam*, *dcm,* and *hsdMS*) systems were sequentially removed. Methyltransferase modules derived from target *Geobacillus* strains were then chromosomally integrated into neutral loci under inducible promoters (*arsB*, *rbsAR*, *aslAB*, and *atpI*) (**Fig. 2b and Table S3**). Correct genomic integration and module stability were verified by locus-specific PCR and reporter (sfGFP) assays (data not shown). Methylation activity of each engineered donor host was further confirmed using methylation-sensitive restriction digestion assays guided by DNMB-derived motif predictions and REBASE annotations (**Fig. 2c-e**).

Using this modular plasmid artificial modification framework, we constructed strain-matched methylation hosts for multiple *Geobacillus* isolates, including EF60045 (MCv1.3.3.123), SJEF4-2 (MCv1.3.3.BG), ATCC12980^T^ (MCv0.3.6.T), HTA426 (MCv1.3.3.AK2), KCTC 3570^T^ (MCv1.3.3.L), and KCTC 3902^T^ (MCv1.3.3) (**Figs. 3 and S2-S3**). We then evaluated whether methylation-matched plasmid DNA could overcome host restriction barriers by electroporating an sfGFP reporter plasmid (pG1Kt-P_igG_-kan-P_pdhA_-sfGFP) prepared either from standard *E. coli* DH5α (*dam*^+^/*dcm*^+^) or from plasmid artificial modification strains harboring the corresponding methylome.

Methylation-matched plasmids substantially improved transformation efficiencies in a subset of strains, including SJEF4-2 and *G. thermodenitrificans* KCTC 3902^T^ (**Fig. 2f**), confirming that R–M compatibility is an important determinant of transformation success. However, no transformants were recovered from several wild-type strains, including *G. stearothermophilus* EF60045, *G. stearothermophilus* ATCC12980^T^, *G. kaustophilus* HTA426, and *G. thermoleovorans* KCTC 3570^T^, despite methylome matching and optimized electroporation conditions. These results indicate that overcoming R–M barriers alone is insufficient to enable DNA uptake and maintenance in many wild *Geobacillus* backgrounds.

Together, these findings demonstrate that plasmid artificial modification can partially bypass classical R–M restriction but does not fully resolve transformation resistance. The persistent refractoriness observed in several strains is consistent with DNMB predictions and DefenseFinder annotations identifying multiple non-canonical defense systems, including Wadjet, Gabija, and pAgo- like modules, which likely function as dominant post-entry barriers to genetic domestication.

### CRISPR-based genome editing reveals non-canonical defense systems as dominant barriers to thermophile domestication

To further enhance DNA delivery, we implemented plasmid artificial modification-assisted conjugative engineering (PACE), in which methylation-matched plasmids generated in engineered donor strains are transferred to *Geobacillus* recipients via pRK24-mediated conjugation (**Figs. 4a and S4**). This approach enabled efficient restriction evasion during DNA transfer. Conjugative delivery of an sfGFP reporter plasmid into SJEF4-2 occurred at high frequency (3.3 ×10^-3^ recipients^-1^), whereas EF60045 yielded only extremely low but reproducible transfer events (∼9 ×10^-9^ recipients^-1^), representing a difference of more than six orders of magnitude under identical donor conditions. Transconjugant colonies frequently displayed heterogeneous fluorescence, including mixtures of fluorescent and non-fluorescent cells, indicating post-entry plasmid silencing or degradation. These results demonstrate that intracellular defense mechanisms continue to restrict plasmid establishment even after successful DNA transfer.

**Figure 4.**
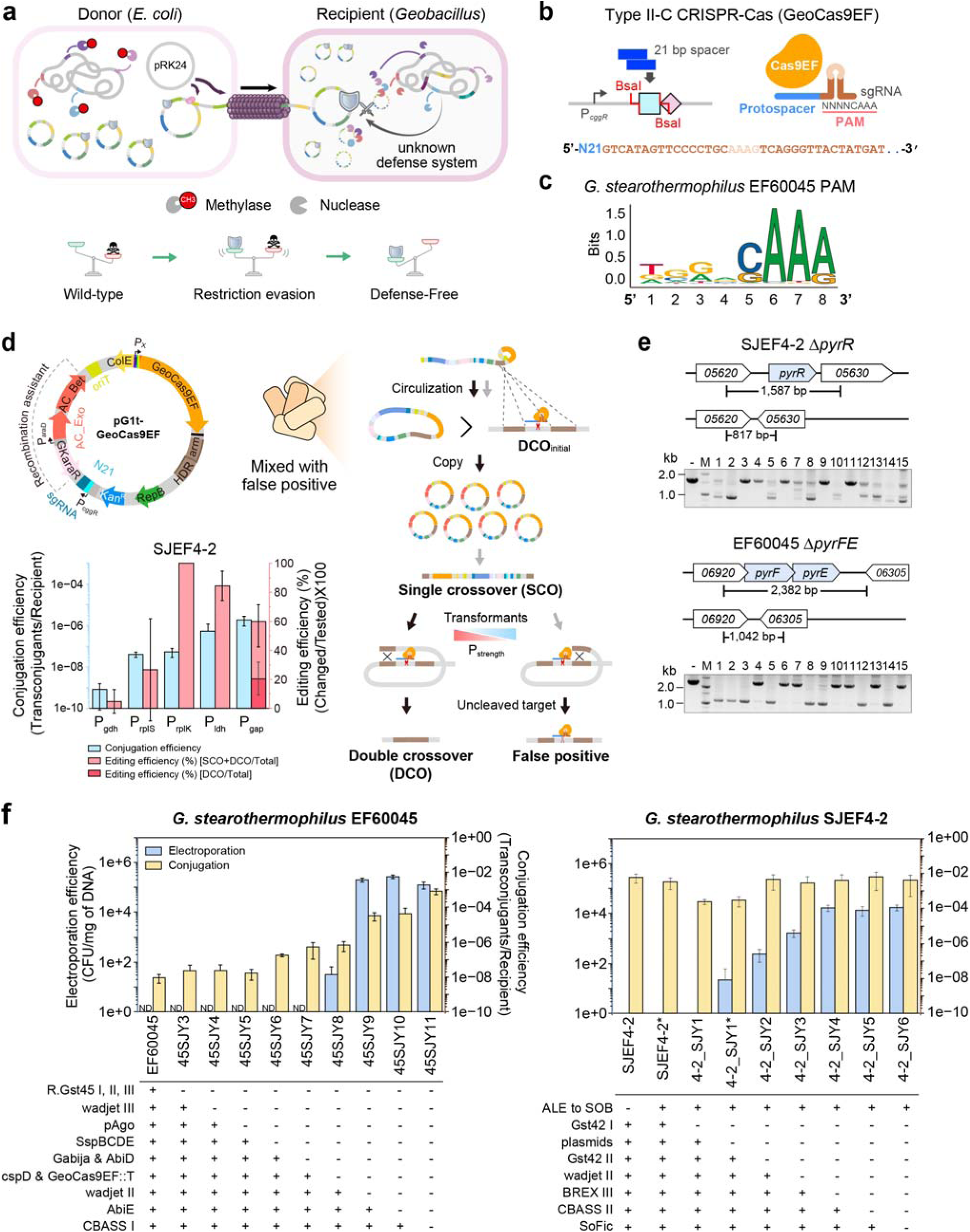
CRISPR-assisted genome editing and removal of defense barriers in *Geobacillus*. a, Plasmid artificial modification–assisted conjugative engineering (PACE). Methylation-matched plasmids produced in engineered *E. coli* donor strains are transferred to *Geobacillus* recipients via pRK24-mediated conjugation. While host-specific methylation enables restriction evasion, intracellular defense systems limit plasmid establishment in wild strains. **b,** Architecture of the native Type II-C CRISPR–Cas system (GeoCas9EF) adapted for genome editing. A 21-bp spacer sgRNA cassette targets genomic loci adjacent to a protospacer-adjacent motif (PAM). **c,** Sequence logo of the preferred PAM recognized by GeoCas9EF in *G. stearothermophilus* EF60045, indicating a consensus 5’-NNNNCAAA- 3’ motif. **d,** CRISPR-mediated genome editing strategy. A conjugative plasmid carrying GeoCas9EF, sgRNA, and homologous repair arms enables genome modification via Cas9-induced cleavage and homologous recombination. Single-crossover (SCO) intermediates are resolved into double-crossover (DCO) mutants or false positives. Promotor optimization for Cas9 expression in strain SJEF4-2 is shown. **e,** PCR validation of representative gene deletions generated by CRISPR editing in SJEF4-2 and EF60045, targeting pyrimidine biosynthesis loci (e.g., *pyrR* and *pyrFE*). Expected fragment sizes are indicated. **f,** Transformation efficiencies following sequential removal of endogenous defense systems. In EF60045 and SJEF4-2, deletion of native plasmids and non-canonical defense modules, including Wadjet II, BREX, CBASS, Gabija, Abi, and SspBCDE, dramatically increased electroporation efficiency (CFU µg ¹ DNA) and conjugation efficiency (transconjugants per recipient), revealing these systems as dominant barriers to genetic domestication.

Because efficient DNA delivery alone did not confer genetic tractability, we next established a native CRISPR-based genome editing platform to interrogate endogenous defense barriers. Genome analysis revealed that *G. stearothermophilus* EF60045 encodes a Type II-C CRISPR-Cas system (GeoCas9EF) sharing 94.8% amino-acid identity with GeoCas9 from ATCC 7953 and 87.4% identity with GtCas9 (**Fig. S5; DataSets 1-3**). In contrast, SJEF4-2 harbors a Type I-C CRISPR system, highlighting subtype diversity among closely related strains. Spacer sequences predominantly matched phages and mobile genetic elements, consistent with active viral defense histories.

Comparative analysis of GeoCas9 orthologs revealed a conserved tracrRNA architecture compatible with a common sgRNA scaffold despite minor sequence variation (**Fig. S5b**). Protospacer- adjacent motif (PAM) profiling indicated that GeoCas9EF (GeoCas9EF2) preferentially recognizes a relaxed 5’-NNNNCAAA-3’ (5’-NNNNSAAR-3’) motif (**Fig. 3b–c**), consistent with PAM preferences (5’-NNNNCWAA-3’) reported for thermophilic Cas9 enzymes ^28^.

To enable programmable editing, we constructed a modular vector expressing Cas9 under transcriptomics-informed promoters and the sgRNA under P_cggR_ control with a customizable N21 protospacer (**Fig. 3d**). Among tested promoters, P_gap_ provided the optimal balance between editing efficiency and cellular tolerance. Targeting non-essential loci such as *pyrR* resulted in cleavage in approximately 60% of transformants, with about 20% already exhibiting double-crossover recombination during early selection (**Fig. 3e**). Efficient plasmid curing (>50% within 12 h of non- selective growth) enabled a complete edit-cure cycle in approximately two days (**Fig. S4**). PCR analysis confirmed precise genome modifications in both SJEF4-2 and EF60045, including deletions of pyrimidine biosynthesis genes such as *pyrR* and *pyrFE* (**Fig. 3e**). These results demonstrate that GeoCas9EF supports efficient genome editing in thermophilic hosts once DNA delivery barriers are overcome.

Having established an efficient editing platform, we used this system to dissect the defense architecture predicted by DNMB analysis. Comparative genomics across 51 (*Para*)*Geobacillus* genomes revealed highly strain-specific repertoires of defense systems, including Wadjet II, BREX, SoFic, MazEF, Abi-like modules, SspBCDE, and CBASS (**Fig. 1**). We hypothesized that these non-canonical defense systems, rather than classical R–M systems alone, constitute the primary barriers to transformation.

Sequential deletion of endogenous plasmids and predicted defense loci in EF60045 confirmed this hypothesis (**Fig. 3f**). Despite methylation-matched donor DNA, the presence of 14 defense modules severely restricted conjugation. Progressive removal of SspBCDE, Gabija, AbiD, Wadjet II, and CBASS I resulted in cumulative increases in DNA transfer efficiency exceeding six orders of magnitude.

Notably, deletion of the Wadjet II alone increased conjugation efficiency from below detection (<10^-8^) to ∼10^-5^, and additional removal of pAgo_Long, Gabija, and CBASS I elevated efficiencies to ∼10^-3^ (**Fig. 3f**). In contrast, deletion of classical R–M systems alone produced minimal improvements, identifying Wadjet II as a dominant barrier to DNA acquisition.

A similar pattern was observed in SJEF4-2 (**Fig. 3f**). Although optimized donor methylation enabled near-saturating conjugation (10^-3^ to 10^-2^), electroporation remained impossible until native plasmids were eliminated. Subsequent deletion of Wadjet_II and BREX_III further increased electroporation efficiency. In EF60045, a spontaneous nonsense mutation in *cspD* altered membrane properties and enabled electroporation, while additional deletion of Wadjet II produced an additional ∼10^3^-fold increase in transformation efficiency (**Fig. S6**).

To evaluate generality, we applied this strategy to additional thermophiles historically resistant to genetic manipulation, including *G. stearothermophilus* ATCC 12980^T^, *G. kaustophilus* HTA426, and *G. thermoleovorans*. Removal of native plasmids together with Wadjet II homologs substantially increased both conjugation and electroporation efficiencies, whereas reintroduction of the *jetD* effector restored transformation resistance across species (**Fig. S7**).

Collectively, these results demonstrate that non-canonical defense systems, particularly Wadjet II, represent the principal barriers to thermophile domestication. Rational removal of these modules enables broad transformation competence, high plasmid productivity, and programmable genome engineering in previously genetically intractable *Geobacillus* strains.

### A hierarchical thermophilic genetic toolkit enables programmable engineering of *G. stearothermophilus*

With major defense barriers removed, we next asked whether these thermophilic hosts could be transformed into programmable engineering platforms. To this end, we established a hierarchical molecular toolkit for *G. stearothermophilus* that integrates tunable replication modules, quantitative promoter libraries, inducible regulatory elements, and scalable genome editing components, enabling precise control of gene dosage and expression in thermophilic chassis.

Genome analysis revealed that *G. stearothermophilus* and related (*Para*)*Geobacillus* strains typically harbor large conjugative plasmids (> 30 kb) but rarely encode small cryptic plasmids (<10 kb) ^25^. Quantitative PCR analysis of wild-type strains (ATCC 12980, EF60063, and SJEF4-2) showed that large native plasmids (e.g., pBSO1, pEF60063-1, and pSJEF4-2-1) are maintained at low copy numbers (∼2–5 copies per chromosome), whereas smaller plasmids (pSJEF4-2-2 and pSJEF4-2-3) reach substantially higher levels (89–106 copies), suggesting diverse replication control mechanisms (**Table S4**).

To identify minimal and high-performance replication elements, we cloned candidate replication proteins into standardized reporter vectors containing a swappable replication module (**Fig. S8a**). Flow cytometry profiling in strain SJEF4-2 revealed distinct sfGFP fluorescence distributions among constructs carrying different replication proteins, consistent with differences in plasmid copy number (**Fig. S8b**). Direct qPCR quantification confirmed that Rep1, Rep5, and Rep7 supported high plasmid copy numbers (>100 copies per chromosome), whereas other replication modules maintained lower or moderate levels (**Fig. S8c**). Importantly, similar relative copy-number patterns were observed across multiple host backgrounds, indicating that these replication modules function as portable and tunable genetic elements.

Additional validation in the defense-reduced strain 45SJY11 demonstrated that the hierarchy of replication strength is largely preserved among domesticated *Geobacillus* hosts (**Fig. S8d-e**). Notably, defense-reduced backgrounds exhibited dramatic improvements in plasmid performance, with plasmid copy numbers increasing from single-digit levels in wild-type strains to hundreds of copies per chromosome, accompanied by stable and homogeneous GFP expression. Removal of defense modules did not impair cellular growth or expression capacity under laboratory conditions (**data not shown**). Together, these results establish a scalable replication panel spanning low- to high-copy regimes for rational gene dosage optimization in thermophilic chassis.

To enable quantitative transcriptional control, we constructed a promoter library guided by transcriptomic profiling of EF60045 grown under different nutrient conditions. RNA-seq analysis identified genes with consistently high expression levels, including loci encoding glycolytic enzymes and housekeeping functions (**Fig. S8f; DataSet 5**). Eleven candidate promoters derived from these genes were fused to an sfGFP reporter and evaluated by flow cytometry (**Fig. S8g and Table S5**). The promoters exhibited a broad dynamic range of activity, with P_gdh_ and P_igG_ among the strongest, thereby providing a graded expression spectrum suitable for pathway balancing and burden-sensitive circuit design. Coverage analysis of the highly expressed P_cggR_ locus further confirmed its robust transcriptional activity and suitability for guide RNA expression (**Fig. S8h**).

To further expand regulatory flexibility, we evaluated inducible systems derived from thermophilic bacteria. The most effective module was an L-arabinose–responsive system from *G. kaustophilus* HTA426 (GK*araR* and P_araD_), which achieved ∼10.6-fold induction with moderate basal leakage (**Fig. S8i**). This inducible system enabled expression of genes that are unstable or toxic in *E. coli*, thereby expanding the accessible expression space for thermophilic biotechnology applications.

Collectively, these components define a modular thermophilic engineering toolkit that supports tunable gene dosage, quantitative expression control, inducible regulation, and cross-strain portability. When combined with the DNMB framework and rational removal of defense barriers, this platform converts previously genetically intractable thermophilic strains into programmable chassis suitable for high-temperature industrial biotechnology.

### Growth-coupled rare sugar screening in the domesticated *Geobacillus* chassis

To demonstrate the utility of the domesticated *Geobacillus* chassis for metabolic engineering and enzyme discovery, we established a growth-coupled screening platform for rare-sugar metabolism using the engineered SJEF4-2 strain. In this design, cellular growth directly reports enzymatic activity: a host incapable of utilizing a given sugar as a sole carbon source will growth only if an introduced enzyme converts that substrate into metabolically usable intermediates. Genome analysis revealed that SJEF4-2 encodes multiple carbohydrate utilization pathways, including the *araBAD* operon for L-arabinose (L- Ara) metabolism, the *galKETR* operon of the Leloir pathway, and a putative *dgoDKAx* cluster corresponding to the De Ley Dourdoroff pathway for D-Gal utilization (**Fig. 4a**) ^16, 29^. Growth profiling confirmed robust utilization of L-Ara and D-Gal but no growth on D-Tag, which instead inhibited growth relative to control conditions (casamino acid-only), indicating the absence of a native D-Tag catabolic route (**Fig. S9a–b**).

**Figure 5.**
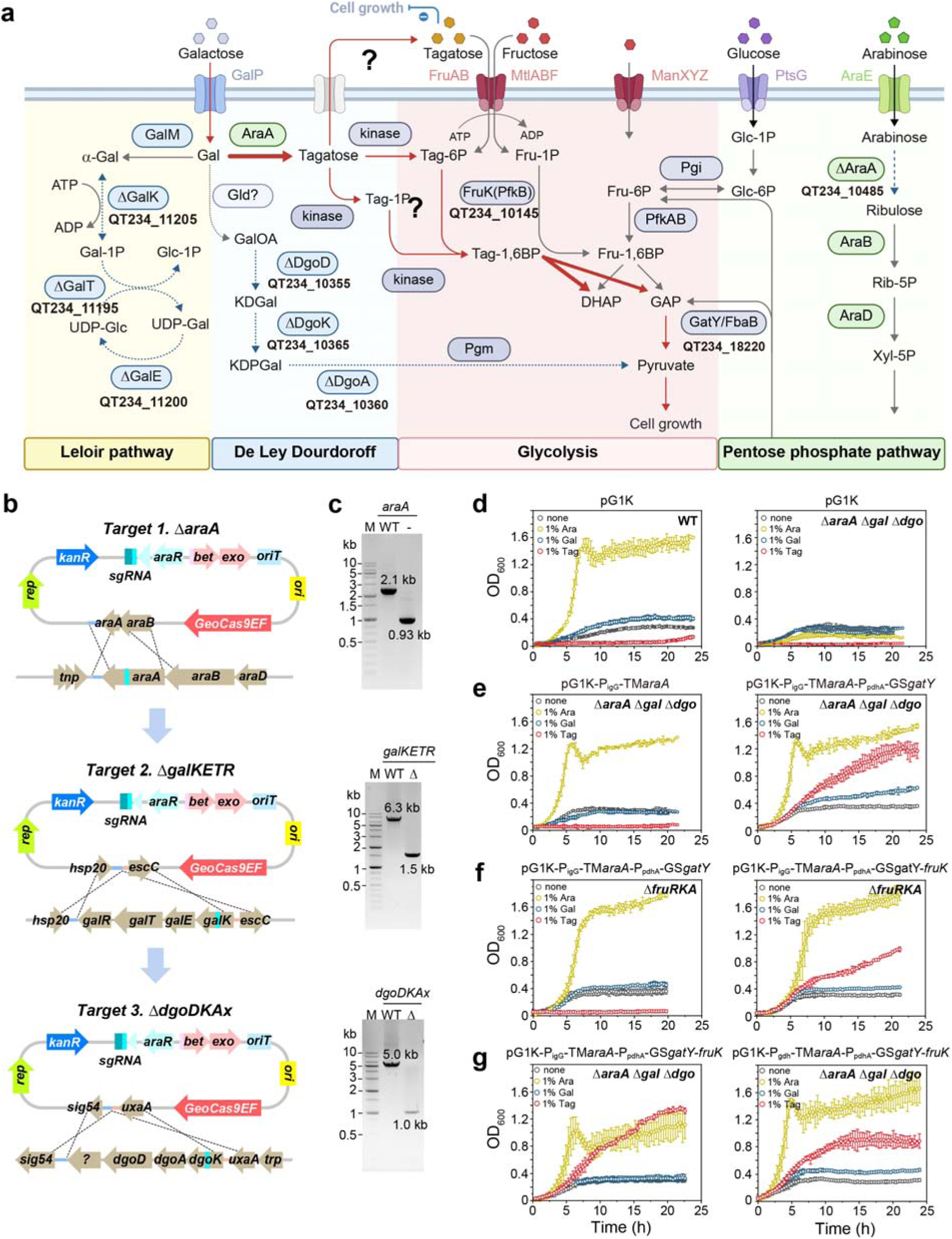
Metabolic rewiring enables tagatose utilization in domesticated *Geobacillus*. a, Reconstructed carbohydrate metabolic network in *G. stearothermophilus*. Native galactose utilization pathways (Leloir and De Ley–Doudoroff pathways) were disrupted by deletion of *galKETR* and *dgoDKA*, while endogenous *araA* was removed to eliminate background isomerase activity. Heterologous expression of *T. maritima* L-arabinose isomerase (*TM_araA*) enabled conversion of D-Gal to D-Tag, which was further metabolized through glycolytic intermediates via tagatose-1,6-bisphosphate cleavage catalyzed by tagatose 1,6-bisphosphate aldolase gene from *G. stearothermophilus* A1 (*GS_gatY*). **b,** CRISPR–Cas9EF-mediated genome editing strategy for sequential deletion of *araA*, *galKETR*, and *dgoDKA* loci in *G. stearothermophilus* SJEF4-2. PCR validation of targeted genome deletions. **c,** Agarose gel electrophoresis confirms successful editing events for *araA*, *galKETR*, and *dgoDKA* loci. M, DNA size marker. **d,** Growth profiles of wild-type (*G. stearothermophilus* 42DF) and engineered strains harboring the control plasmid (pGK1-NULL) in minimal medium supplemented with different sugars, demonstrating altered sugar utilization following gene deletions. **e,** Complementation of the engineered chassis with heterologous *TM_araA* and *GS_gatY* expression constructs in *G. stearothermophilus* Δ*araA* Δ*galKETR* Δ*dgoDKA* mutant strains. Growth recovery on tagatose indicates functional establishment of the engineered tagatose utilization pathway. **f,** Growth profiles of Δ*fruRKA* mutant strains expressing either *TM_araA–GS_gatY* or *TM_araA*–*GS_gatY*–*fruK*. Introduction of *fruK* enhanced growth on tagatose, indicating that fructose kinase activity facilitates downstream metabolism of tagatose-derived intermediates. **g,** Effect of promoter strength on tagatose-dependent growth. Engineered Δ*araA* Δ*galKETR* Δ*dgoDKA* mutant strains harboring *TM_araA–GS_gatY–fruK* under either the *PigG* or *Pgdh* promoter show differential growth kinetics, demonstrating that increased *TM_araA* expression improves flux through the engineered D-Tag pathway. Growth curves represent cultures grown in minimal *Geobacillus* salts medium supplemented with individual sugars. Data represent mean ± s.d. from three biological replicates.

To create a selective screening background, we sequentially deleted *galKETR, dgoDKAx*, and the endogenous *araA* gene, eliminating both L-Ara metabolism and native D-Gal catabolism (**Fig. 4b–c**). The resulting mutants showed loss of growth on their respective substrates, and the triple mutant (Δ*araA* Δ*galKETR* Δ*dgoDKAx*) was unable to utilize either L-Ara or D-Gal while maintaining normal growth on other nutrients (**Fig. 4d**). To enable D-Tag metabolism, we introduced *GSgatY*, encoding tagatose-1,6- bisphosphate aldolase from *G. stearothermophilus* A1. Expression of *GSgatY* restored growth on D-Tag (**Fig. 4e**). However, growth remained limited, suggesting a bottleneck in upstream phosphorylation or conversion steps.

To reconstruct a complete D-Gal–to–D-Tag utilization pathway, we expressed a thermostable L- arabinose isomerase (L-AI) from *Thermotoga maritima* MSB8 (TM_AraA), which converts D-Gal to D- Tag. In this background, deletion of *fruRKA* abolished tagatose utilization, whereas complementation with *fruK* restored growth (**Fig. 4f**), indicating that FruK functions as a D-Tag kinase that channels the sugar into central metabolism. These results also imply the presence of an endogenous tagatose uptake system independent of canonical D-Fru transporters (**Fig. 4a**).

Reintroduction of *araA* restored L-Ara metabolism and enabled isomerization of D-Gal to D-Tag, completing the engineered pathway through FruK-dependent phosphorylation, phosphofructokinase (*pfkB*), and *GSgatY*-mediated cleavage to glycolytic intermediates (**Fig. 4g**). Although growth on D-Gal was recovered, it remained slower than growth on D-Tag, indicating that the isomerization step catalyzed by TM_AraA is rate-limiting. Consistent with this interpretation, increasing TM_AraA expression using the strong constitutive P_gdh_ promoter markedly improved growth rate and biomass yield (**Figs. 5g and S9c**).

Together, these results establish an activity-dependent, growth-coupled screening system in a thermophilic host, enabling direct selection of functional enzyme variants and synthetic metabolic pathways at high temperature. This engineered *Geobacillus* chassis therefore provides a powerful platform for thermostable enzyme evolution, pathway prototyping, and rare-sugar biotransformation under industrially relevant conditions.----------------------------

## Discussion

Developing genetically tractable microbial hosts remains a major obstacle to exploiting non-model organisms, particularly extremophiles, for functional genomics and biotechnology. Despite their unique metabolic capabilities and industrial potential, most thermophilic bacteria remain genetically inaccessible due to multilayered defense systems and the absence of systematic domestication strategies. Here, we present a systematic framework for the programmable domestication of thermophilic bacteria that integrates computational genome analysis, targeted removal of genetic barriers, and modular engineering tools to convert previously intractable strains into versatile genetic platforms (**Fig. 5a**). Central to this effort is the DNMB pipeline, which consolidates task-oriented scripts and curated reference databases into an integrated analytical suite capable of rapidly identifying defense systems and genetic bottlenecks (**Figs. 1–2**). By enabling rational prioritization of engineering targets, this framework transforms domestication from a trial-and-error process into a predictable workflow for engineering thermophilic hosts (**Fig. 5**). Importantly, this framework is not limited to *Geobacillus* but provides a generalizable strategy for identifying and overcoming genetic barriers across diverse thermophilic taxa.

**Figure 6.**
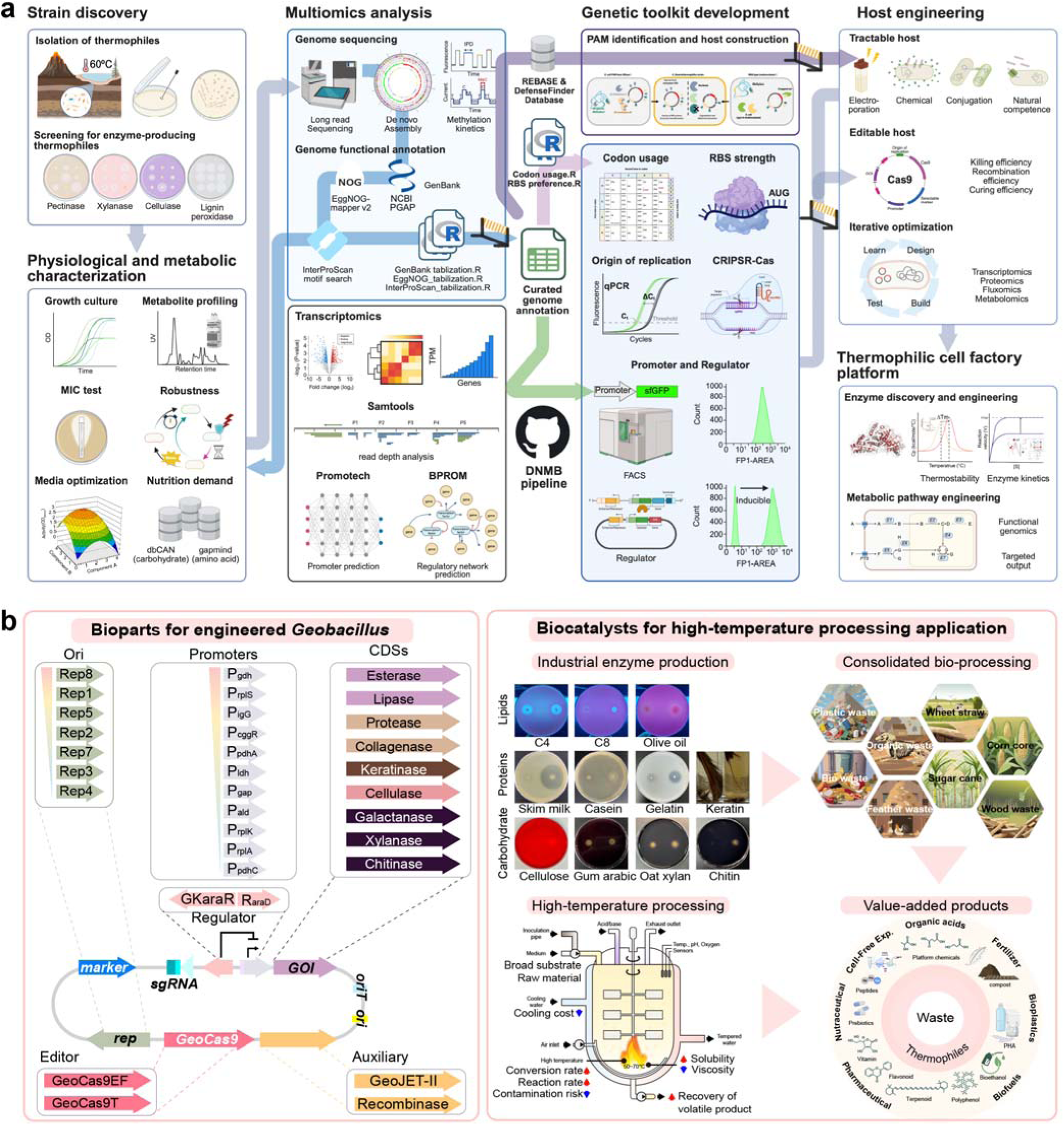
Programmable thermophilic cell factory platform based on engineered *Geobacillus*. **a**, Conceptual workflow for developing genetically tractable thermophilic hosts. The pipeline integrates strain discovery, multi-omics characterization, and DNMB-guided genome analysis to identify genetic barriers and construct programmable engineering toolkits. These include plasmid artificial modification systems, codon usage optimization, promoter and regulator libraries, replication modules, and CRISPR– Cas genome editing tools. Iterative host engineering enables the development of tractable and editable thermophilic strains suitable for metabolic and enzyme engineering. **b,** Modular bioparts and industrial applications of the engineered *Geobacillus* platform. Orthogonal replication origins, promoters, regulators, coding sequences, and CRISPR-based editors enable programmable pathway construction. The resulting thermophilic chassis supports high-temperature biocatalysis and consolidated bioprocessing, converting diverse substrates—including lipids, proteins, carbohydrates, and lignocellulosic residues—into value-added products such as organic acids, biofuels, bioplastics, nutraceuticals, and pharmaceuticals.

R–M systems are widely regarded as the principal barriers to horizontal DNA transfer in bacteria ^30^. Consistent with this view, *in vivo* methylation strategies were initially required to enable efficient DNA delivery into *Geobacillus* strains. Heterologous expression of methylases provided a practical entry point for genetic manipulation, and notably, the *Geobacillus*-common Type III methylase (GCC^6m^AT) alone was sufficient to methylate target sequences, challenging the assumption that both modification and restriction subunits are required for enzymatic activity (**Figs. 3 and S1c**) ^31^. However, methylation alone proved insufficient to fully overcome genetic barriers. Several adenine methylases (MS.SJEF4-2I, M.AlwI, M.BstEII, MS.60045II, MS.GK426II) exhibited toxicity in *E. coli*, likely reflecting widespread perturbation of DNA-dependent processes such as replication, transcriptional regulation, and SOS responses ^32^. These effects necessitated tightly regulated expression systems (e.g., rhamnose-inducible promoters) to mitigate methylation-induced toxicity (**Figs. 3 and S2-3**) ^33^. Collectively, these findings indicate that R–M evasion alone cannot ensure genetic accessibility and that genetic intractability arises from multilayered host defenses rather than a single dominant mechanism.

Beyond classical R–M systems, our results reveal that non-canonical defense systems constitute dominant barriers to genetic domestication in thermophilic *Geobacillus* (**Fig. 1**). Systematic deletion of recently described defense modules ^34–37^, including Wadjet II, Gabija, and BREX III, dramatically increased transformation efficiencies (**Fig. 3f**). In particular, removal of native plasmids together with Wadjet II produced the largest improvement in conjugation efficiency, identifying this system as a critical barrier to DNA acquisition (**Figs. 4f and S7**). These observations reinforce emerging evidence that bacterial defense architectures are multilayered and highly strain-specific, often involving mechanisms distinct from classical restriction systems.

The Wadjet defense system belongs to a family of SMC-like complexes that recognize circular DNA substrates and cleave them via a toprim-like nuclease subunit (JetD) ^36, 38–40^. Its widespread distribution across (*Para*)*Geobacillus* species, except for *Parageobacillus* and *G. thermodenitrificans*, suggests a major role in restricting foreign DNA in thermophilic environments (**Fig. 1**). The coexistence of Wadjet systems with large endogenous plasmids (>30 kb) in many strains raises the possibility of size- or topology-dependent discrimination that allows certain DNA molecules to evade cleavage. Structural variation within SMC-like domains may influence substrate recognition ^41^, and the susceptibility of related complexes to DNA-mimic proteins suggests potential avenues for transient inhibition of Wadjet activity ^42^. Alternatively, controlled activation of the JetD nuclease could be exploited as a programmable DNA elimination system.

Efficient electroporation requires additional optimization beyond defense system removal, indicating that intracellular defenses are not the sole determinants of transformability. Even after eliminating R–M systems, successful electro-transformation depended on membrane properties and host physiology (**Fig. 3f**). Previous studies have identified host factors such as *manB* and *mutS* that may influence competence, potentially through effects on membrane integrity ^43, 44^. A spontaneous *cspD* nonsense mutation that altered membrane fluidity markedly improved electroporation efficiency, indicating that cellular envelope characteristics represent an additional layer governing genetic accessibility (**Figs. 4f and S6**). Together, these findings suggest that successful transformation requires coordinated optimization of both intracellular defense systems and cell envelope properties.

In parallel with removing genetic barriers, we established an integrated thermophilic engineering toolkit enabling iterative genome modification and tunable gene expression (**Figs. S5 and S8**). Conjugation proved particularly effective due to the transfer of single-stranded DNA that is readily recombined by native repair pathways ^45, 46, 47^. Paired with this delivery method, the native Type II GeoCas9EF enabled rapid genome editing cycles with high efficiency, complementing previously reported CRISPR approaches in related thermophiles ^44^ (**Figs. 4a-e and S4**). Optimizing Cas9 expression under the P*_gap_* promoter minimized toxicity while reducing false-positive colonies during counter selection (**Fig. 3d**).

At the expression level, several regulatory systems previously reported in other *Parageobacillus* species were not functional in this host, likely reflecting species-specific regulatory hierarchies (**Fig. S8**). In contrast, modulation of plasmid copy number via alternative replication origins proved highly effective for tuning expression levels ^48^. The high-copy pGxt vector, which lacks copy-control genes present in native plasmids (e.g., ParA, CopG), yielded substantially higher expression levels (**Fig. S8c-d and Table S4**). The coexistence of multiple native plasmids further suggests that compatible multi-plasmid expression systems could be developed in *Geobacillus*, analogous to those reported in related thermophilic species ^21^. Collectively, these components provide a versatile genetic toolkit for thermophilic synthetic biology (**Fig. S8**).

To demonstrate practical utility, we engineered a growth-coupled rare sugar screening platform in the domesticated *Geobacillus chassis.* By rewiring native carbohydrate metabolism and introducing thermostable enzymes, cellular growth was made dependent on the activity of L-AIs converting D-Gal to D-Tag (**Fig. 4**) ^49^. Thermophilic hosts offer intrinsic advantages for such applications, including enhanced substrate solubility, increased enzymatic rates, and reduced side reactions at elevated temperatures. This platform enables rapid screening of thermostable enzyme variants and synthetic pathways under process-relevant conditions.

Beyond this proof-of-concept application, the domesticated *Geobacillus* chassis supports large-scale genome engineering. The high efficiency of homologous recombination using relatively short homology arms (∼500 bp) suggests that genome-wide editing strategies may be feasible. Coupled with automated design tools, this system could enable the construction of comprehensive knockout libraries and systematic pathway rewiring for functional genomics and strain optimization.

Finally, the engineered strain retains broad degradative capabilities toward diverse substrates, including proteins, lipids, and complex polysaccharides, while operating at temperatures incompatible with mesophilic hosts (**Figs. 6b and 9d**). These features, combined with improved genetic accessibility, position *Geobacillus as* an attractive chassis for high-temperature biomanufacturing and extremophile biotechnology (**Fig. 5b**). Such properties are particularly advantageous for consolidated bioprocessing, thermophilic biocatalysis, and sustainable manufacturing processes where contamination resistance and elevated reaction rates are advantageous.

Collectively, this study demonstrates that the genetic recalcitrance of thermophilic bacteria is not an intrinsic property but arises from identifiable and removable defense architectures. By providing a generalizable roadmap for domestication, our work expands the scope of synthetic biology beyond traditional model organisms and opens new opportunities to harness the metabolic diversity of thermophilic microbes. By enabling the systematic domestication of previously inaccessible microbes, this approach may unlock a vast reservoir of metabolic diversity across extreme environments for synthetic biology and industrial biotechnology.

## Materials and methods

### Bacterial strains and growth conditions

All bacterial strains used in this study are listed in **Table S3**. *Escherichia coli* strains were cultured in Terrific Broth (TB) or Luria-Bertani (LB) medium at 37°C. For *Geobacillus* spp., a modified LB (mLB) supplemented with 0.5% (w/v) of the designated sugar was used to assess sugar-dependent growth profiles.

For recovery following electroporation, several media were evaluated, including Super Optimal Broth (SOB), modified SOC (mSOC), 2SPYNG, 2SPY, and their sugar concentration variants. Growth kinetics were monitored using an RTS-8 bioreactor (Biosan; Riga, Latvia). For bioreactor inoculation, 10 mL of preheated medium was dispensed into 50 mL centrifuge tubes and inoculated with 100 µL of bacterial suspension. Cultures were incubated at 55°C for *Geobacillus* or 37°C for *E. coli* with orbital shaking at 2,000 rpm for up to 72 h, depending on strain and growth phase.

For directed evolution experiments, a defined minimal medium was prepared per liter with the addition of 10 mL vitamin solution (2 mg of biotin, 2 mg of folic acid, 10 mg of pyridoxine-HCl, 5 mg of thiamine-HCl, 5 mg of riboflavin, 5 mg of nicotinic acid, 5 mg of calcium pantothenate, 0.1 mg of vitamin B_12_, 5 mg of p-aminobenzoic acid, 5 mg of lipoic acid) and 10 mL trace element stock solution containing 2 g of nitrilotriacetic acid, 0.18 g of ZnSO_4_·7H_2_O, 3 g of MgSO_4_·7H_2_O, 0.5 g of MnSO_4_·2H_2_O, 1 g of NaCl, 0.1 g of FeSO_4_·7H_2_O, 0.01 g of H_3_BO_3_, 0.18 g of CoSO_4_·7H_2_O, 0.01 g of CuSO_4_·5H_2_O, 0.1 g of CaCl_2_·2H_2_O, 0.1 g of KAI(SO_4_)_2_·12H_2_O, 0.001 g of Na_2_SeO_3_·5H_2_O, 0.025 g of NiCl_2_·6H_2_O, 0.01 g of Na_2_MoO_4_·2H_2_O.

### Detection of DNA methylation and motif analysis

DNA modifications and associated sequence motifs were characterized using the Base Modification and Motif Analysis pipeline implemented in SMRTLink v10.2.1 (Pacific Biosciences). PacBio circular consensus sequencing (CCS) reads were aligned to the complete genome reference to detect methylated bases. Motif identification results, particularly the “motifs.csv” file generated by SMRTLink, were processed to identify methylation patterns. Detected motifs and modification types were subsequently submitted to the REBASE DB (https://rebase.neb.com) for comparative analysis and public access.

### Motif analysis pipeline for restriction-modification (R–M) and defense systems

Comprehensive genome annotation was performed using coding sequences (CDSs) initially predicted by GeneMarkS2+ and deposited in the National Center for Biotechnology Information (NCBI). Protein motif and functional annotation were conducted using InterProScan v5.60-92.0, integrating multiple signature DBs including TIGRFAM (v15.0), SFLD (v4), SUPERFAMILY (v1.75), PANTHER (v17.0), Gene3D (v4.3.0), Hamap (v2021_04), ProSiteProfiles (v2022_01), Coils (v2.2.1), SMART (v7.1), CDD (v3.20), PRINTS (v42.0), PIRSR (v2021_05), ProSitePatterns (v2022_01), AntiFam (v7.0), Pfam (v35.0), MobiDBLite (v2.0), PIRSF (v3.10), Phobius (v1.01), SignalP (v4.1), and TMHMM (v2.0c) ^50^.

CDSs were additionally mapped to the eggNOG v5.0 DB using eggNOG-mapper v2.1.10 under default parameters ^51^. Functional annotations, including Kyoto Encyclopedia of Genes and Genomes (KEGG) modules, pathways, reactions, rClass, TCDB, BRITE categories, CAZy families, BiGG reactions, and PFAMs generated using a DIAMOND blastp-based automated pipeline (**Fig. 1a**). Annotation results for each ORF were integrated into the GenBank feature table using an in-house script (**DataSet 1** and **DataSet 2**).

R–M systems were identified by BLAST searches against the REBASE Gold Standard DB with an E-value cutoff of ≤ 0.01 ^52^. Consensus motifs were assigned based on PROSITE patterns and literature- defined signatures, including SAM binding motifs (FxGx[AG]), catalytic motifs ([DS]PP[FY] for methylase; PDx16[DE]xK for restriction enzyme), and ATP-binding P-loop motifs (GxGK[ST]).

Non-canonical bacterial defense systems were predicted using DefenseFinder v2.0.0 ^34^. System counts were derived from the “Systems” output table, satisfying the MacSyFinder rules defining mandatory and accessory components for each defense system.

### Comparative genomics

Pan-genome analysis was performed using the Bacterial Pan Genome Analysis (BPGA) pipeline version

1.3 ^53^. Fifty-one complete (*Para*)*Geobacillus* genomes were obtained from NCBI GenBank (**Sheet 1 of DataSet 3**). Protein sequences from each genome were clustered using USEARCH with a 40% identity threshold, allowing classification of gene families into core, accessory, and strain-specific components (**DataSets 3 and 4)**. The pan-genome and core-genome size were calculated iteratively with each added genome (**Sheet 2 of DataSet 3**). Clustering outputs were formatted for downstream analysis using <u>BPGAconverter</u>. Defense system distributions were visualized using <u>DefenseViz</u>, which aggregates DefenseFinder outputs and summarizes system counts per genome (**Sheet 3 of DataSet 3**). Polysaccharide-utilization loci (PULs) and CAZyme gene clusters (CGCs) were identified using dbCAN3, which integrates HMMER, DIAMOND, and Hotpep (**Sheet 4 of DataSet 3**). To further analyze R–M system diversity, a subgroup analysis was conducted using 16 *Geobacillus* genomes with PacBio long-read methylome data available in REBASE (**DataSet 4**).

### Plasmid construction

All primers used in this study are listed in **Tables S7-15**. Plasmids were constructed using NEBuilder HiFi DNA Assembly Master Mix (New England Biolabs) following the manufacturer’s instructions. PCR-amplified fragments and linearized vectors were assembled in a single reaction and transformed into chemically competent *E. coli*. All plasmid constructions were sequence-verified prior to use.

### Construction of *E. coli* plasmid artificial modification hosts

The parental plasmid artificial modification strain *E. coli* ZYCY10P3S2T (#MN900A-1), derived from BW27783, was obtained from System Biosciences (SBI) ^54^. Whole-genome sequencing data for this strain were kindly provided by Christopher D. Johnston ^55^. Following the strategy described for systematic evasion of R–M barriers ^55^, *Geobacillus*-derived methyltransferases were integrated into the *E. coli* chromosome using the KIKO system ^56^. Expression levels were validated using an sfGFP reporter, and fluorescence was quantified using a S3e^TM^ Cell Sorter System (Bio-Rad).

### Preparation of electrocompetent *Geobacillus* cells

Electrocompetent *G. stearothermophilus* cells were prepared by growing cultures in SOB medium at 55°C until OD_600_ = 1.0–1.4. Cells were cooled on ice for 20 min, harvested by centrifugation (4,000×g, 10 min, 4°C), and washed four times with ice-cold electroporation buffer containing 10% glycerol. Cells were resuspended to 1% of the original culture volume.

For selection on media lacking divalent cations (e.g., mLB), kanamycin concentration was reduced to 12 µg/mL; whereas in divalent-cation-rich media (e.g., SOB), the standard 50 µg/mL concentration was used.

### Electroporation of *Geobacillus*

Electroporation was performed by mixing 60 μL of electrocompetent cells with 2.4 µL of plasmid DNA. Fifty microliters of the mixture were transferred to prechilled 1 mm GenePulser^TM^ cuvettes (Bio-Rad). Electroporation conditions were 2.0 kV, 10 μF, and 600 Ω. Post-pulse. Cells were immediately recovered in SOB medium and incubated at 55°C with shaking for ∼3 h before plating on selective agar. After recovery, cells were pelleted (4,500×g for 5 min at 25°C), resuspended in 200 μL SOB, and plated on selective SOB agar plates containing 50.0 μg/mL kanamycin or 7.0 µg/mL thiamphenicol for *G. stearothermophilus* EF60045 and SJEF4-2, *G. thermodenitrificans* KCTC3902^T^, and *G. thermoleovorans* KCTC3570^T^. SOB containing 12.5 µg/ml kanamycin or 7.0 µg/mL thiamphenicol for *G. stearothermophilus* ATCC 12980^T^ and *G. kaustophilus* HTA426.

### Assessment of electroporation efficiency

Electroporation efficiency was evaluated using the reporter plasmid pG1Kt-P_igG_-kan-P_pdhA_-sfGFP, extracted from various plasmid artificial modification hosts (*E. coli* DH5a, MCv1.3.3.BG, MCv1.3.3.123, MCv0.3.6.T, MCv1.3.3.AK2, MCv1.3.3.L, and MCv1.3.3). Detailed plasmid genotypes and methylation signatures are listed in **Tables S1-2**. DNA was diluted to 200 ng/µL, and 2 µL (400 ng) was used for each electroporation. Fifty microliters of *Geobacillus* electrocompetent cells were used per transformation reaction.

### Transconjugation of *Geobacillus spp*

Transconjugation was performed with modifications based on a previously described protocol ^45^. The broad-host-range conjugative plasmid pRK24 (Addgene plasmid # 51950, courtesy of Farren Isaacs) ^57^ was first introduced into *E. coli* MC variants and selected on tetracycline (10 µg/mL). In a second step, plasmids carrying the *oriT* sequence of pRK24 were transformed into the same *E. coli* host. Recipient cells (*Geobacillus* spp.) were streaked on non-selective plates and incubated at 55°C one day before conjugation. Because donor growth varied depending on the vector, donor cells were streaked on TB agar supplemented with tetracycline (10 µg/mL) and the corresponding vector-specific antibiotic marker two days before conjugation and incubated overnight at 32°C. These donor strains were subcultured on the same medium again on the evening before mating. On the day of conjugation, recipient *Geobacillus* cultures were started in round-bottom tubes, whereas donor cultures were inoculated at 2% into 25 mL pre-warmed TB and incubated at 32°C with vigorous shaking until they reached the exponential phase (OD_600_ of ∼1.0 after 2–3 h). When donor cultures reached OD_600_ of ∼0.3, rhamnose was added to a final concentration of 1.0% to induce methylase expression when required, and cultures were incubated for an additional 2 h. Final OD_600_ values were approximately 0.5 for donors and 1.0 for recipients. Donor cells were harvested by centrifugation (4,500×g, 5 min), washed twice with antibiotic-free LB broth (5 mL), and kept on hold while recipient cells remained at room temperature. Donor and recipient cells were then mixed at ratios ranging from 1:1 to 1:4, centrifuged (4,500×g, 10 min), and gently resuspended in approximately 200 µL of residual LB. Aliquots (10 µL) were spotted onto antibiotic-free LB agar plates and incubated overnight at 32°C. The following day, mating spots were harvested by repeated rinsing with 2 mL SOB and transferred to 14 mL round-bottom tubes. Suspensions were incubated at 55°C with shaking at 250 rpm for ∼3 h with the tubes positioned at an angle of ∼30° to improve aeration. Cultures were then plated on selective media for transconjugant recovery.

### Conjugation efficiency

Conjugation efficiency was quantified using the shuttle plasmid pG1Kt-P_igG_-kan-P_pdhA_-sfGFP and donor *E. coli* PAM strains harboring pRK24. Efficiency was calculated as the number of transconjugant colonies per recipient cell. Detailed genotypes and methylation patterns are provided in **Table 1**. All assays were performed in biological triplicate.

### Construction of *Geobacillus* mutants

To enable conjugative DNA transfer, the *oriT* sequence from pRK24 was inserted adjacent to the pBR322 origin in a modified *Geobacillus*-*E. coli* shuttle vector. Mutants were generated by Gibson assembly ^58^ using homologous arms of approximately 0.5 kb flanking the target deletion or insertion site (**Fig. 3a**). Several Cas9 expression plasmids were constructed, including the *oriT*-integrated pNW33N- derived pThermoCas9 (pGtCas9t), the pG1K-based pG1Kt-GeoCas9EF (Cas9 from *G. stearothermophilus* EF60045), and pG1Kt-GeoCas9T (Cas9 from *G. stearothermophilus* ATCC 12980^T^). These plasmids conferred kanamycin or thiamphenicol resistance and functioned in multiple *Geobacillus* strains.

Initial transformants carrying single crossover (SCO) integrations were selected, followed by recovery of double crossover (DCO) mutants through counter-selection. DCO strains were validated by PCR amplification and sequencing of flanking regions. After 2–3 passages in antibiotic-free SOB broth, diluted cultures (∼10^3^ CFU/mL) were spread on SOB agar and incubated overnight at 55°C. Resulting colonies were screened by PCR using AccuPower® ProFi Taq PreMix (Bioneer, Daejeon, Korea) and tested for antibiotic sensitivity to confirm allelic replacement and loss of the plasmid backbone.

### Transcriptomic analysis

Total RNA was extracted from *Geobacillus* cells using the RNeasy Mini Kit (Qiagen GmbH, Hilden, Germany), followed by rRNA depletion with the NEBNext rRNA Depletion Kit (NEB). RNA-seq libraries were prepared and sequenced on an Illumina NovaSeq 6000 (Illumina, Inc., San Diego, CA, USA), yielding an average of 28.5 million reads per sample after quality trimming. Reads were trimmed using Trimmomatic v0.39 with the following parameters: ‘ILLUMINACLIP: TruSeq3-SE.fa:2:30:10 LEADING:3 TRAILING:3 SLIDINGWINDOW:4:15 MINLEN:30 -threads 12 -phred33’ ^59^. Cleaned reads were aligned to the reference genome (*G. stearothermophilus* EF60045, CP128453.1) using Bowtie2 with default settings ^60^. Gene-level counts were generated using HTSeq-count, and differential expression analysis was performed using the DESeq2 package in R ^61^. Transcript abundance was normalized as transcripts per million (TPM). Genes showing a ≥ 2-fold change in normalized TPM and *P* < 0.05 were considered differentially expressed. Complete results are provided in **DataSet 5**.

### Illumina library preparation and single-nucleotide variation (SNV) analysis

To improve genome accuracy, circulated assemblies were corrected using Illumina short-read resequencing. Genomic libraries were prepared using the TruSeq Nano DNA Library Prep Kit (Illumina) and as 2 × 150 bp paired-end reads on a NovaSeq 6000 platform. Raw reads were processed using Trimmomatic v0.39 for adapter removal and quality trimming, followed by quality assessment with FastQC v0.11.9 ^59^. Variants were identified using breseq with default parameters ^62^, and results were visualized using <u>breseqConverter</u>.

### Fluorescence measurement

Green fluorescence intensity of superfolder GFP (sfGFP) was measured using the FL1 channel (excitation 488 nm, emission 510-540 nm) of an S3e^TM^ Cell Sorter (Bio-Rad). Fluorescence data (FL1- AREA) from 50,000 cells per sample were collected from three biological replicates and summarized statistically. Raw FCS files were normalized to 10,000 events using the R package flowCore ^63^, and visualized using ggcyto ^64^.

### Screening of replication origins and plasmid copy number determination

To identify functional replication origins from native *G. stearothermophilus* plasmids, the RepBST1 (Rep1) region of pG1K-sfGFP was replaced with replication-associated genes, including replication proteins, initiator proteins, or helix-turn-helix domain-containing proteins derived from native plasmids pSJEF4-2-1 (Rep8), pSJEF4-2-2 (Rep7), pSJEF4-2-3 (Rep5), pEF60063-2 (Rep3), and pEF60063-3 (Rep4). RepB of pNW33N (Rep2) and RepSTK1 (Rep6) were used as additional positive controls alongside RepBST1.

Plasmid copy number (CN) was determined by qPCR using 5 pg of genomic DNA and 0.25 pmol of each primer per reaction. Amplification was performed using the CFX 2StepAmp+Melt protocol with 2× BioRad iQ^TM^ SYBR Green^®^ SuperMix. Cq values were normalized to the single-copy chromosomal genes *rpoD*, *infB,* and *dnaA*. Copy number was calculated using the -2^ΔΔCq^ method. Plasmid DNA was extracted from late-exponential to stationary-phase cultures using the Promega Wizard^®^ DNA Extraction kit. Primer sequences are listed in **Table S9**.

### Promoter and replication origin screening

Promoter candidates were selected from highly expressed genes identified by transcriptomic analysis. Each promoter (P*_igG_*, P*_cggR_*, P*_pdhA_*, P*_pdhC_*, P*_rplK_*, P*_gap_*, P*_gdh_*, P*_ldh_*, P*_ald_*, and P*_rplKA_*) was cloned upstream of sfGFP and introduced into *G. stearothermophilus* SJEF4-2. Fluorescence was measured after 18 h of cultivation in mLB medium containing kanamycin. Fluorescence intensity was recorded from 50,000 cells per culture using a S3e^TM^ Cell Sorter System (Bio-Rad).

## Comparative analysis of Cas9 proteins

Forty-seven non-redundant Cas9/Csn1 sequences from *Geobacillus* species were obtained from NCBI (**Fig. S5a**). Multiple sequence alignment was performed using MUSCLE ^65^, and structural predictions were generated using AlphaFold2 (**Fig. S5b**)^66^. Domain boundaries were assigned based on previously reported Cas9 structures (**Fig. S5c**)^67^.

## Acknowledgements

We are grateful to Prof. Katherine P. Lemon (Forsyth Institute) for generously providing the genome sequence of ZYCY10P3S2T (System Biosciences), a BW27783-derived strain that serves as the mother strain of the MC series. We thank Dr. Richard J. Roberts and Dr. Tamas Vincze of New England Biolabs for their helpful discussions and guidance on methylation pattern analysis.

## Author’s Contributions

J.Y. Sung, S.J. Lee, S.B. Kim, and D.W. Lee formulated the research plan. J.Y. Sung, M.H. Lee, J.S. Park, H.B. Kim, G. Darima, D.G. Kim, and H.W. Cho performed the experiments. J.Y. Sung conducted computational analysis and script coding. J.Y. Sung, S.J. Lee, S.B. Kim, and D.W. Lee analyzed the data.

J.Y. Sung and D.W. Lee wrote the manuscript. S.B. Kim and D.W. Lee conceived, planned, supervised, and managed the study.

