## Supplementary information for "Programmable domestication of thermophilic bacteria through removal of non-canonical defense systems"

**Running title:** Programmable domestication of thermophilic bacteria

**Corresponding authors:**

**Seong Bo Kim**

Bio-Living Engineering Major, Global Leaders College, Yonsei University, Seoul 03722, South Korea.

**Dong-Woo Lee**

Department of Biotechnology, Yonsei University, Seoul 03722, Republic of Korea.

**Table S1. Restriction-modification (R–M) repertoire of *G. stearothermophilus* EF60045**

| **REBASE** | **Locus_ tag (QT235_)** | **Type** | **Length (AA)** | **Best hit**  **Isoschizo-mer**  **(identity %)** | **Detected Motif (No. of sites)** | **Catalytic motif** | **SAM binding**  **motif** | **P-loop** |
| --- | --- | --- | --- | --- | --- | --- | --- | --- |
|  | 07815 | IV | 326 | BclIMrrP (100%) |  |  |  |  |
|  | 10955 | IV | 335 | McrB |  |  |  | _91_PDx11ExK_104_ |
|  | 07625 | III (M) | 425 | Partial (transposase) | - | _117_DPPY_120_ |  |  |
|  | 07635 | III (M) | 243 | EcoPI (incomplete) |  |  | _27_FxGxA_31_ |  |
|  | 07640 | III (R) | 984 | R.GthID1IIIP (94%) |  |  |  |  |
|  | 07975 | III (R) | 865 | M.Gth3570II  M.Gst10I | GCC^6m^AT  (9512/12812) |  |  | _133_PDx16DxK_152_  _812_GTGKT_816_ |
| M.Gst45IV | 07980 | III (M)* | 398 |  |  | _113_DPPY_116_ | _356_FxGxA_360_ |  |
|  | 07985 | III (M)* | 79 |  |  |  |  |  |
| R.Gst45I | 04845 | I (R) | 1080 |  | T^6m^ACNNNNNNCTC (451/712)  G^6m^AGNNNNNNGTA (442/712) |  |  | _504_DEAH_508_  _400_GTGKT_404_ |
| M.Gst45I | 04850 | I (M) | 493 | M.Gst10II (100%) |  | _255_NPPF_258_ |  |  |
| S.Gst45I | 04855 | I (S) | 485 |  |  |  | _276_FxGxA_280_ |  |
| M.Gst45II | 01945 | I (M) | 484 |  | RT^6m^AYNNNNNCTC (615/892)  G^6m^AGNNNNNRTAY (581/892) | _272_NPPF_275_ |  |  |
| S.Gst45II | 01950 | I (S) | 369 |  |  |  |  |  |
| R.Gst45IIP | 01955 | I (R) | 1113 |  |  |  |  | _373_GSGKT_377_ |
| R.Gst45III | 00960 | I (R) | 779 |  | CC^6m^ANNNNNNNCTC (1096/1547)  G^6m^AGNNNNNNNTGG (949/1547) |  |  | _192_GTGKT_196_ |
| M.Gst45III | 00970 | I (M) | 479 | M.BstXI (26.2%) |  | _276_NPPY_279_ |  |  |
| S.Gst45III | 00975 | I (S) | 496 | S. BstX1 (46.0%) |  |  | _444_FxGxA_448_ |  |

Table S2. Restriction-modification (R–M) repertoire of *G. stearothermophilus* SJEF4-2

| **REBASE** | **Locus_tag  (QT234_)** | **Type** | **Length**  **(AA)** | **Best hit**  **Isoschizomer**  **(identity %)** | **Detected**  **Motif (No. of sites)** | **Catalytic motif** | **SAM binding**  **motif** | **P-loop** |
| --- | --- | --- | --- | --- | --- | --- | --- | --- |
| M.Gst42IIIP | 17960 (plasmid) | II (M) | 414 | M.BamHI | GGAT^4m^CC |  |  |  |
| R.Gst42IIIP | 17950 (plasmid) | II (R) | 211 | BamHI |  |  |  |  |
| M. Gst42I | 07090 | III (M) | 477 | M.Gth3570II (92.4) | GCC^6m^AT | _114_DPPY_117_ | _356_FxGxA_360_ |  |
| R. Gst42I | 07085 | III (R) | 1253 | Mba11I (36.2) |  |  |  | _132_PDx16DxK_152_  _812_GTGKT_816_ |
| M.Gst42II | 01885 | I (M) | 484 | M.EF60063I (100.0) | AA^6m^AYNNNNNRTCNC GNG^6m^AYNNNNNRTT |  |  |  |
| S.Gst42II | 01890 | I (S) | 389 | S.EF60063I (100.0) |  |  |  |  |
| R.Gst42II | 01895 | I (R) | 1113 | M.EF60063I (100.0) |  |  |  |  |
|  | 08745 | I (M) | 643 | M1.Gsp12II (98.0%) |  |  |  |  |
|  | 08755 | I (M) | 515 | M.Gsp300I (100.0) | - |  |  |  |
|  | 08760-08770 | I (S) | 134  310 | S.Gsp300I  08760 (28%)  08770 (72%) |  |  |  |  |
|  | 08775 | I (R) | 1020 | R.Gsp300I (99.0) |  |  |  |  |

Table S3. Bacterial strains used in this study

| **Strain** | **Genotype** | **Relevant characteristics** | **Reference** |
| --- | --- | --- | --- |
| ***G. stearothermophilus*** | | | |
| ATCC 12980^T^ | *G. stearothermophilus* ATCC 12980^T^ | Type strain | ^1^ |
| ATCC 12980^T^-1 | ATCC 12980^T^ ∆pBSO1 |  | This study |
| ATCC 12980^T^-1w | ATCC 12980^T^-1 ∆QSJ10_01575-90 (∆Wadjet_II) |  | This study |
| EF60045 | *G. stearothermophilus* EF60045 | Type strain | ^2^ |
| 45SJY1 | EF60045 ∆QT235_00960 (∆R.Gst45III) |  | This study |
| 45SJY2 | 45SJY1 ∆QT235_04845 (∆R.Gst45I) |  | This study |
| 45SJY3 | 45SJY2 ∆QT235_01955 (∆R.Gst45II) |  | This study |
| 45SJY4 | 45SJY3 ∆QT235_07660-75 (∆Wadjet III) |  | This study |
| 45SJY5 | 45SJY4 ∆QT235_07775 (∆pAgo_Long) |  | This study |
| 45SJY6 | 45SJY5 ∆QT235_11435-60 (∆SspBCDE) |  | This study |
| 45SJY7 | 45SJY6 ∆QT235_01955-70 (∆Gabija) |  | This study |
| 45SJY8 | 45SJY7 ∆QT235_03025 (∆AbiD) | Prophage-associated large deletion | This study |
| 45SJY9 | 45SJY8 ∆QT235_01700-20 (∆Wadjet_II) |  | This study |
| 45SJY10 | 45SJY9 ∆QT235_17105-10 (∆AbiE) |  | This study |
| 45SJY11 (45DF) | 45SJY10 ∆QT235_00950-60 (∆CBASS_I) |  | This study |
| EF60063 | *G. stearothermophilus* EF60063 | Wild-type | This study |
| SJEF4-2 | *G. stearothermophilus* SJEF4-2 | Wild-type | ^2^ |
| 42SJY1 | SJEF4-2 ∆QT234_01895 (∆R.Gst42I) |  | This study |
| 42SJY1_p∆23 | 42SJY1 ∆pSJEF4-2-2 ∆pSJEF4-2-3 |  | This study |
| 42SJY1_p∆12 | 42SJY1 ∆pSJEF4-2-1 ∆pSJEF4-2-2 |  | This study |
| 42SJY1_p∆13 | 42SJY1 ∆pSJEF4-2-1 ∆pSJEF4-2-3 |  | This study |
| 42SJY1_p∆123 (42SJY1*) | 42SJY1 ∆pSJEF4-2-1 ∆pSJEF4-2-2 ∆pSJEF4-2-3 |  | This study |
| 42SJY2 | 42SJY1* ∆QT234_08775 (∆R.GstII, inactive form) |  | This study |
| 42SJY3 | 42SJY2 ∆QT234_01685-700 (∆Wadjet_II) |  | This study |
| 42SJY4 | 42SJY3 ∆QT234_06800-15 (∆BREX_III) |  | This study |
| 42SJY5 | 42SJY4 ∆QT234_08520-80(∆CBASS_II) |  | This study |
| 42SJY6 (42DF) | 42SJY5 ∆QT234_03560 (∆SoFic) |  | This study |
| 42DF ∆*araA* | 42DF ∆*araA* |  | This study |
| 42DF ∆*dgoDKA* | 42DF ∆*dgoDKA* |  | This study |
| 42DF ∆*galKETR* | 42DF ∆*galKETR* |  | This study |
| 42DF ∆*araA* ∆*galKETR* | 42DF ∆*araA* ∆*galKETR* |  | This study |
| 42DF ∆*araA* ∆*galKETR,* ∆*dgoDKA* | 42DF ∆*araA* ∆*galKETR* ∆*dgoDKA* |  | This study |
| ***G. kaustophilus*** |  |  |  |
| HTA 426 | *G. kaustophilus* HTA426 | Type strain | ^3^ |
| HTA 426-1 | HTA 426 ∆pHTA426 |  | This study |
| HTA 426-1w | HTA 426-1 ∆GK_0317-20 (∆Wadjet_II) |  | This study |
| ***G. thermoleovorans*** |  |  |  |
| KCTC 3570^T^ | *G. thermoleovorans* KCTC 3570^T^ wildtype |  | ^4^ |
| KCTC 3570-1 | KCTC 3570 ∆pLDW-1 |  | This study |
| KCTC 3570-1w | KCTC 3570-1 ∆QT3570_01570-85 (∆Wadjet_II) |  | This study |
| ***G. thermodenitrificans*** | | | |
| KCTC 3902^T^ | *G. thermodenitrificans* KCTC 3902^T^ | Type strain | ^3, 5^ |
| ***E. coli*** |  |  |  |
| DH5a | *fhuA2 lac(del)U169 phoA glnV44 Φ80' lacZ(del)M15 gyrA96 recA1 relA1 endA1 thi-1 hsdR17* |  |  |
| ZYCY10P3S2T (MCv0) | BW27783 *Cp8.araE ΔendA bla.lacY A177C 3BAD.I-SceI 2BAD.ΦC31 4BAD.Φ31 4BAD.Φ31* |  | ^6^ |
| MCv0.0.1 | MCv0 ∆(*mrr-hsdRMS-symE-mcrBC*) | To express heterologous methylases | This study |
| MCv0.0.2 | MCv0.0.1 ∆*dcm* | To remove C^5m^CWGG methylase | This study |
| MCv0.0.3 | MCv0.0.2 ∆*mcrA* | To express heterologous methylases | This study |
| MCv0.0.4 | MCv0.0.3 ∆*recA* | To increase copy number and plasmid stability | This study |
| MCv0.0.5 | MCv0.0.4 ∆*nupG* | To increase copy number | This study |
| MCv0.0.6 | MCv0.0.5 ∆*deoR* | To increase copy number | This study |
| MCv0.3.6 | MCv0.0.6 ∆*arsB*::(P_J23100_- QSJ10_06485-90) | To express GCC^6m^AT methylase | This study |
| MCv0.3.6.T | MCv0.3.6 ∆*rbs(A-R)*::*(rhaR-*P_rhaB_*-*QSJ10_15455-15460) | To express RGTN^6m^ ACC methylase (M.BstPI) | This study |
| MCv1.0.3. | MCv0.0.3 ∆*dam* | To remove G^6m^ATC methylase | This study |
| MCv1.3.3 | MCv1.0.3 ∆*arsB*::(P_J23100_- QSJ10_06485-90) | To express GCC^6m^AT methylase | This study |
| MCv1.3.3.3 | MCv1.3.3 ∆*rbs*(*A-R*)::(P_QT235_00965_- QT235_00965-75) | To express CC^6m^ ANNNNNNNCTC/  G^6m^AGNNNNNNNTGG methylase | This study |
| MCv1.3.3.13 | MCv1.3.3.3 ∆*atpI*::(P_QT235_04850_-QT235_04850-55) | To express T^6m^ACNNNNNNCTC/ G^6m^AGNNNNNNGTA methylase | This study |
| MCv1.3.3.123 | MCv1.3.3.13 ∆*aslAB*::(*rhaR-*P_rhaB_*-*QT235_01945-50) | To express RT^6m^AYNNNNNCTC/  G^6m^AGNNNNNRTAY methylase | This study |
| MCv1.3.3.B | MCv1.3.3 ∆*atpI*::(QT234_17955-60) | To express GGAT^4m^CC methylase (Isoschizomer of the M.BamHI) | This study |
| MCv1.3.3.BG | MCv1.3.3.B ∆*aslAB*::(*rhaR*-P_rhaB_-QT234_01885-90) | To express AA^6m^AYNNNNNRTCNC/GNG^6m^AYNNNNNRTT methylase | This study |
| MCv1.3.3.A | MCv1.3.3 ∆*atpI*::(*rhaR*-P_rhaB_- GKP08) | To express GG^6m^ATC methylase(Isoschi- zomer of the M.AlwI) | This study |
| MCv1.3.3.AK | MCv1.3.3.A ∆*aslAB*::(*rhaR*-P_rhaB_-GK1380-1) | To express TT^6m^ ACNNNNGTR/Y^6m^ACNNNNGTAA methylase | This study |
| MCv1.3.3.AK2 | MCv1.3.3.AK | To express MS.*Gka*426II but not functional | This study |
| MCv1.3.3.L | MCv1.3.3 ∆*atpI*::(GT3570_17465) | To express C^4m^CWGG | This study |

Table S4. Copy numbers of native plasmids in *G. stearothermophilus* SJEF4-2

| **Strain** | **Plasmid** | **Copy number per chromosome (mean ± s.d.)*** |
| --- | --- | --- |
| ATCC12980 | pBSO1 | 4.82 ± 0.86 |
| EF60063 | pEF60063-1 | 2.77 ± 0.86 |
| SJEF4-2 | pSJEF4-2-1 | 4.14 ± 0.97 |
| SJEF4-2 | pSJEF4-2-2 | 100.8 ± 62.8 |
| SJEF4-2 | pSJEF4-2-3 | 91.2 ± 46.9 |
| SJEF4-2 ∆pSJEF4-2-2 ∆pSJEF4-2-3 | pSJEF4-2-1 | 4.9 ± 2.2 |
| SJEF4-2 ∆pSJEF4-2-1 ∆pSJEF4-2-3 | pSJEF4-2-2 | 89.1 ± 40.6 |
| SJEF4-2 ∆pSJEF4-2-1 ∆pSJEF4-2-2 | pSJEF4-2-3 | 106.4 ± 42.2 |

*Copy numbers were calculated by qPCR using the 2^-∆Ct^ method relative to the chromosomal *rpoB* gene. ΔCt = Ct(*repX*) − Ct(*rpoB*).

**Table S5. Promoter activity screening in *E. coli* and *G. stearothermophilus* using sfGFP reporter**

|  | ***E. coli* MCv1.3.3.123** | | | | ***G. stearothermophilus* SJEF4-2** | | | |
| --- | --- | --- | --- | --- | --- | --- | --- | --- |
| **Promoter** | **Mean** | **CV** | **s.d.** | **Median** | **Mean** | **CV** | **s.d.** | **Median** |
| P_PTrplS_ (Control) | 2747.53 | 88.18 | 2422.75 | 2358.00 | 9256.9 | 16.9 | 1562.2 | 10000.0 |
| P_pdhA_ | 73.44 | 168.05 | 123.42 | 51.00 | 4919.6 | 53.49 | 2631.4 | 4795.0 |
| P_pdhC_ | 66.76 | 261.63 | 174.67 | 46.00 | 125.3 | 142.6 | 178.6 | 68.7 |
| P_ldh_ | 39.99 | 146.51 | 58.59 | 34.00 | 3392.0 | 77.1 | 2614.7 | 2662.4 |
| P_gdh_ | 1185.6 | 240.25 | 2848.46 | 48.00 | 9530.6 | 17.0 | 1624.5 | 10000.0 |
| P_rplK2_ | 1520.7 | 106.0 | 1612.5 | 899.2 | 815.0 | 126.1 | 1027.4 | 483.1 |
| P_rplKA_ | 185.9 | 196.7 | 365.6 | 59.1 | 756.4 | 129.6 | 980.6 | 500.4 |
| P_igG_ | 2249.2 | 177.58 | 3994.09 | 46.00 | 7693.1 | 38.8 | 2981.3 | 9091.0 |
| P_gap_ | 66.58 | 293.48 | 195.35 | 46.00 | 1086.8 | 122.3 | 1329.1 | 656.6 |
| P_ald_ | 123.77 | 223.38 | 276.47 | 48.00 | 866.4 | 128.1 | 1109.5 | 420.3 |
| P_cggR_ | 2517.44 | 163.75 | 4122.31 | 51.00 | 7471.4 | 44.1 | 3296.8 | 9044.2 |

s.d.: standard deviation.

**Table S6. Replication initiator proteins and origins of replication used in this study**

| **Replication**  **protein** | **Replication mechanism** | **Plasmid origin** | **Original**  **host** | **Homologs**  **(derivatives)** | | **Core size**  **(bp)** | **Compatible strains** | **Reference** |
| --- | --- | --- | --- | --- | --- | --- | --- | --- |
| RepBSO1 | Theta replication | pBSO1 | ATCC 12980 | | (Mob^+^) | 4,004 |  | ^7^ |
| RepBST1 (rep1) | Theta replication | pG1AK-sfGFP  pUCG18  pBST22 | ATCC 7953 | | pEF60063-1  (Mob^+^) | 873 | *Parageobacillus thermoglucosidasius,*  *Parageobacillus sp. NUB3621* | ^8^  ^9, 10^ |
| RepB (rep2) | Rolling circle | pNW33N | *B. coagulans* (pBC1)  *S. aureus* (pC194) | |  | 1,005 | *Clostridium thermocellum,*  *Geobacillus thermodenitrificans* K1041 | ^11^ |
| RepSTK1 (rep6) | Rolling circle | pSTE33 | STK1 (cryptic) | |  | 1,883 | *Geobacillus kautophilus* HTA426,  *Geobacillus thermodenitrificans* K1041 | [PDB: 4CIJ]  ^12^ |
| RepA-GST (rep8) | Theta replication | pSJEF4-2-1 | SJEF4-2 | | pSJEF4-2-1 (Mob^+^),  pNCI002,  pGeo12a,  pLDW-2, pDSM14590 |  |  | In this study |
| RepGST1 (rep3) | Rolling circle | pEF60063-2 | EF60063 | |  | 7,074  (rep gene: 3,527) |  | In this study |
| RepGST2 (rep4) | Rolling circle | pEF60063-3 | EF60063 (cryptic) | | pGTG4S,  pGTG5 | 2,214 | *Parageobacillus thermoglucosidasius* | In this study |
| RepGST4 (rep5) | Rolling circle | pSJEF4-2-3 | SJEF4-2 (cryptic) | |  | 1,996 bp |  | In this study |
| RepGST3 (rep7) | Rolling circle? | pSJEF4-2-2 | SJEF4-2 (cryptic) | |  | 6,676 bp |  | In this study |

**Table S7. Primers used for the construction of a plasmid artificial modification host in *Geobacillus***

| **Category** | **Primer** |  | **Sequence (5'-3')** | **T_m_(℃)** |
| --- | --- | --- | --- | --- |
| Common template | pTarget_inv. LA | F | AGCTTCTGCAGGTCGACTCTAGAG | 61 |
|  | pTarget_inv. RA | R | TCGAGTTCATGTGCAGCTCCATAAGC | 63 |
| **∆(*mcrC-mrr*)** | | | | |
| sgRNA | *hsdR*-sgRNA inv. | F | AGCTGGTTTTATTCACCTCCGTTTTAGAGCTAGAAATAGCAAGTT | 74 |
|  |  | R | AGGTGAATAAAACCAGCTACTAGTATTATACCTAGGACTGAGCTAG | 72 |
| Left arm | *mcrC* arm | F | CGACCTGCAGAAGCTCCGGCGTTCACTATACCGATCGC | 79 |
|  |  | R | CCTGATATTGCGGTAAGTTTTATAGAAAATACCGCTCCCG | 72 |
| Right arm | *mrr* arm | F | ACTTACCGCAATATCAGGCCGGATGCGGCTG | 77 |
|  |  | R | GCTGCACATGAACTCGATGACCAAAACCGACGTCGCAG | 78 |
| Check primer | ∆(*mcrC-mrr*) check | F | GGCTAAAGCCACCAACGCTGTC | 69 |
|  |  | R | ATCCCGGCCCGATTATTCAGACC | 69 |
| **∆*dcm*** | | | | |
| sgRNA | *dcm*-sgRNA inv. | F | GTCAAAGTCTTTCTCACCCGGTTTTAGAGCTAGAAATAGCAAGTT | 66 |
|  |  | R | GGGTGAGAAAGACTTTGACACTAGTATTATACCTAGGACTGAGCTAG | 65 |
| Left arm | *vsr* arm | F | CGACCTGCAGAAGCTTCGGTAAGCGCTTCATCCGTCAGC | 73 |
|  |  | R | GAAATCTATGCATGGCCGACGTTCACGATA | 64 |
| Right arm | *yedJ* arm | F | GTCGGCCATGCATAGATTTCACCGGCCATC | 68 |
|  |  | R | GCTGCACATGAACTCGATGTCCAGGATGCGGATCGGCTG | 72 |
| Check primer | ∆*dcm* check | F | GCGCTCGGCGTAATTGTCTTCG | 63 |
|  |  | R | CGGAGAAAATCGAGGCCGTTTGTC | 62 |
| **∆*mcrA*** | | | | |
| sgRNA | *mcrA*-sgRNA inv. | F | GATATACGCTTGAAGATGTGGTTTTAGAGCTAGAAATAGCAAGT | 65 |
|  |  | R | CACATCTTCAAGCGTATATCACTAGTATTATACCTAGGACTG | 62 |
| Left arm | *pinE* arm | F | TCGACCTGCAGAAGCTGCAGGTGACACTCTGGTTGTCTGGAA |  |
|  |  | R | AATGTGACCACGACAATACTACTTTTATTGATAAAATTGCAACAAGTTGC | 65 |
| Right arm | *idh* arm | F | TATTGTCGTGGTCACATTTAAGACGTAATACCCTACAGGGT | 66 |
|  |  | R | TGCACATGAACTCGAACTCAAAAGCAGTCAAGAGTGTCCTCCA |  |
| Check primer | ∆*mcrA* check | F | CAGAGCCTTCTCCCAAACCAACG | 62 |
|  |  | R | GACACGGTAATTCAACCTACATGTGG | 61 |
| **∆*recA*** | | | | |
| sgRNA | *recA*-sgRNA inv. | F | TCGTCGTTGACTCCGTGGGTTTTAGAGCTAGAAATAGCAAGTT | 65 |
|  |  | R | ACGGAGTCAACGACGATAACTAGTATTATACCTAGGACTGAGCTAG | 66 |
| Left arm | *pncC* arm | F | CGACCTGCAGAAGCTAGCGAACAGGTTGGGCAGGCG | 74 |
|  |  | R | AAACAAGACGATTTTACTCCTGTCATGCCGGGTAATACCGG | 68 |
| Right arm | *recX* arm | F | GGAGTAAAATCGTCTTGTTTGATACATAAGGGTCGCATCTGC |  |
|  |  | R | CTGCACATGAACTCGACAAATCTCCTGGATATCTTCCATCAGATAGCCAC |  |
| Check primer | ∆*recA* check | F | GGCTGGTAACTGAAAAGTGGGAAT | 59 |
|  |  | R | TTGCAACGCCAACACCATCT | 61 |
| **∆*nupG*** | | | | |
| sgRNA | *nupG*-sgRNA inv. | F | CCACATGCAGCTGTATATGTTTTAGAGCTAGAAATAGCAAGT | 65 |
|  |  | R | ATATACAGCTGCATGTGGCTACTAGTATTATACCTAGGACTG | 64 |
| Left arm | *mtlC* arm | F | TCGACCTGCAGAAGCTTGCGCACAAATATCTCGGCATGGTCC | 73 |
|  |  | R | TTCTTTGCGTAAGTTAATTTCCTCACATCGTGATGCG | 65 |
| Right arm | *speC* arm | F | ATGTGAGGAAATTAACTTACGCAAAGAAAAACGGGTCGCCA | 67 |
|  |  | R | TGCACATGAACTCGATATCTGCGTGAGAACGGCATTGTGC | 69 |
| Check primer | ∆*nupG* check | F | CTGAAAAACCGTCTGAAGAGCCGC | 62 |
|  |  | R | GGCGAATATAGCGACTTTGGCGT | 62 |
| **∆*deoR*** | | | | |
| sgRNA | *deoR*-sgRNA inv. | F | AAGAAACGCGCGGCAATGGTTTTAGAGCTAGAAATAGCAAGTT | 69 |
|  |  | R | ATTGCCGCGCGTTTCTTTGACTAGTATTATACCTAGGACTGAGCTAG | 68 |
| Left arm | *dacC* arm | F | CGACCTGCAGAAGCTTATGGCATTGCTGGGTAAAGCATTG | 70 |
|  |  | R | AGAGGGATTTGACGTATAACCGGATGACGTTTCG | 66 |
| Right arm | *ybjG* arm | F | TTATACGTCAAATCCCTCTGAATAGTTATTGAAGCGAGC | 64 |
|  |  | R | TGCACATGAACTCGAATGATCTCGTTGGCGATTTTTATTGCTAAAG | 67 |
| Check primer | ∆*deoR* check | F | GCCAAAAAACTGGGTCTGACCAAC | 62 |
|  |  | R | CAACGACTTGCCTGTATTGGCTCCC | 64 |
| **∆*dam*** | | | | |
| sgRNA | *dam*-sgRNA inv. | F | CAACTCTGCTTCCGGGAAATGTTTTAGAGCTAGAAATAGCAAGTT | 67 |
|  |  | R | ATTTCCCGGAAGCAGAGTTGACTAGTATTATACCTAGGACTGAGC | 67 |
| Left arm | *rpe* arm | F | CGACCTGCAGAAGCTAGACTGACCGCCGAAACCAGG | 72 |
|  |  | R | AGAAAAATGTTTCACCCGCGAAAAAATAATTCTCAAGG | 63 |
| Right arm | *damX* arm | F | CGCGGGTGAAACATTTTTCTTCATGCTGACTAAC | 65 |
|  |  | R | GCTGCACATGAACTCGACCCGCTGCTGGGGCGAAG | 74 |
| Check primer | ∆*dam* check | F |  |  |
|  |  | R |  |  |
| **∆*arsB*::(*QSJ10_06485-06490_*K152A) MR’.Gst45IV (*Geobacillus* common)** | | | | |
| sgRNA | *arsB*-sgRNA inv. | F | AATCGCGGCGATATCCACGTTTTAGAGCTAGAAATAGCAAGTT | 68 |
|  |  | R | GGATATCGCCGCGATTGTACTAGTATTATACCTAGGACTGAGCTAG | 67 |
| Left arm | *arsB* up arm (P_J23100_ hang) | F | CGACCTGCAGAAGCTGTCCTGACCATCGTATTGGTTATCTGG | 70 |
|  |  | R | CTGTACCTAGGACTGAGCTAGCCGTCAATCCACCGGCACCATCACCG | 74 |
| promoter | P_J23100_ (arsB_up hang) | F | CCGGTGGATTGACGGCTAGCTCAGTCC | 67 |
|  | P_J23100_ | R | GGTATATCTCCTTCTTAAAGTTAAACAAAATTATTTCTAGAGGGCTAGC | 63 |
| Insert | QJS10_06485 (P_J23100_ hang) | F | CTTTAAGAAGGAGATATACCATGCTTTACGTTTCTCCTGCCAAAATCGC | 67 |
|  | QSJ10_06490 (T7 term hang) | R | GCGGCCGCCTAGCGAGCGGGCTGCTTCG | 77 |
| terminator | T7 terminator (QSJ10_06490_hang) | F | CGCTCGCTAGGCGGCCGCACT | 71 |
|  | T7 terminator (arsB_down hang) | R | CGCGGCGATACAAAAAACCCCTCAAGACCCG | 70 |
| Right arm | *arsB* down arm (T7ter hang) | F | AGGGGTTTTTTGTATCGCCGCGATTGTTGCCACG | 71 |
|  | arsB_down (pTargetF hang) | R | TGCACATGAACTCGAGTCCCAAATCGCAGCCAATC | 69 |
| Check primer | *arsB* check | F | CGCAACCTGGCTCGACAAAACTG | 63 |
|  |  | R | CGCAGGCCGGGTTGTGATAAATGG | 64 |
| **∆*rbs*(*A-R*)::(QT235_00965-75) MS.Gst45III** | | | | |
| sgRNA | *rbsC*-sgRNA inv. | F | TGAGAACGCCGATCTGTTGTTTTAGAGCTAGAAATAGCAAGTT | 66 |
|  |  | R | CAGATCGGCGTTCTCAGTACTAGTATTATACCTAGGACTGAGCTAG | 66 |
| Left arm | *rbsA* arm  (QT235_00965 hang) | F | CGACCTGCAGAAGCTCGGCATTGTGTATATCTCCGAAGACCG | 72 |
|  |  | R | TTCAGGGGTGGGATTGTCCCCCTCCCTCTTTTGTTTG | 70 |
| Insert 1 | QT235_00955 (rbsA hang) | F | CTCTGCTGGTGCTGATCGGAAGCGGATGAAACTTGGTTTTATGATGACAC | 71 |
|  | QT235_00955 (linker) | R | TTCAGGGGTGGGATTGTCCCCCTCCCTCTTTTGTTTG | 70 |
| Insert 2 | QT235_00975 (linker) | F | GACAATCCCACCCCTGAAAGAAGGGGTGTTTTG | 67 |
|  | QT235_00965-75 (rbsAR hang) | R | TCGGTGCACAGTAACAGCCTCAACTTGTCGACTTTGCCGACAATC | 72 |
| Right arm | rbsR_arm | F | GCTGTTACTGTGCACCGAAACG | 61 |
|  | *rbsR* arm (pTarget hang) | R | TGCACATGAACTCGACCCCAGTTCATCTTTCGGTTGGTGGAT | 71 |
| Check primer | *rbs(A-R)* check | F | GGACCTGGCGTTAACGATGTCTCT | 62 |
|  |  | R | GGCACAACGACTGTCGAACAATTCC | 62 |
| **∆*rbs*(*A-R*)::(P_rplsWT_-QT234_08745-08760, 08770) M.Gst42II (inactive form)** | | | | |
| promoter | P_rplsWT_ (pTarget-rbsAR hang) | F | CTCTGCTGGTGCTGATCGAACAATCGTTAAAGCGGACGTTTTTGCGCCGC | 75 |
|  | pTarget_P_rplsWT__hang_inverse | R | TCGAATCACTCCTTATCTAGACAATGC | 58 |
| Insert 1 | QT234_08745 (P_rplsWT_ hang) | F | CTAGATAAGGAGTGATTCGAATGAGCCGTGACAAAGCGTTGC | 69 |
|  | QT234_08760 (08770 hang) | R | ATAAACATCTATTTCATCCAGCTAACATGTTTCTGGGATTG | 63 |
| Insert 2 | QT234_08770 (08760 hang) | F | GCTGGATGAAATAGATGTTTATTTCACCGCTAACATTATTGCCAGG | 66 |
|  | QT234_08770(terminator hang) | R | GCGAAATTGAGCTCTTACTAAAGCACTTTTTCTACTTGTTTTTCTGCTTTAG | 65 |
| terminator | pTarget_terminator_hang_inverse | F | TAGTAAGAGCTCAATTTCGCGC | 58 |
|  | terminator (pTarget hang) | R | TCGGTGCACAGTAACAGCTGATACCTTAGGCTTCGCAAAAAAACG | 70 |
| **∆*rbs*(*A-R*)::(QSJ10_15455-60) CM.Gst42II (isoschizomer of M.BstEII)** | | | | |
| rhaR-P_rhaB_ | rhaR (rbsA up hang) | F | CTCTGCTGGTGCTGATCGTTAATCTTTCTGCGAATTGAGATGACGCCAC | 71 |
|  | rhaB (CM.BstEII_hang) | R | CTATATCACCCACTTACGACCAGTCTAAAAAGCGCCTGA | 67 |
| Insert | CM.BstEII (P_rhaB__hang) | F | GTCGTAAGTGGGTGATATAGATGAAATATGATTTAAATG | 59 |
|  | M.BstEII (terminator hang) | R | GAGCTCTTATTACAATCGATTGGCTTGAAAATTTATAATATCAGCCCATT | 65 |
| **∆*rbs*(*A-R*)::(P_rplsWT_-GK343-4)MS.Gka426II** | | | | |
| Insert | GK343_F (P_rplsWT_ hang) | F | gtctagataaggagtgattcgaatgctcacaggcgaattgcgca | 70 |
|  | GK344_R (term hang) | R | gcgaaattgagctcttactatcttacattacacaaaattcttgacggtaccttctc | 67 |
| **∆*atpI*::(P_QT235_04845_-QT235_04850-55) MS.Gst45II** | | | | |
| sgRNA | *atpI*-sgRNA inv. | F | CTTTCGCTGGTGTATGCGCCGTTTTAGAGCTAGAAATAGCAAGTT | 69 |
|  |  | R | GGCGCATACACCAGCGAAAGACTAGTATTATACCTAGGACTGAGC | 69 |
| Left arm | *atpI* down arm | F | AGAGTCGACCTGCAGAAGCTATCAGCTTGCTTTTGCCATGGTACATG | 71 |
|  |  | R | TACACCAGCGAAAGGCCGGGT | 66 |
| Insert | QT235_04855 (atpI hang) | F | CGGCCTTTCGCTGGTGTATCACCGCTGTCCGCTTGACG |  |
|  | P_QT235_04845_ hang | R | GGGATGGACAGATGAACAACCGAGAAATTGTCCAAAAGCTG | 68 |
| promoter | P_QT235_04845_ (QT235_04850 hang) | F | GTTGTTCATCTGTCCATCCCTCATCTTTGTCTCTTTTGCAA | 65 |
|  | P_QT235_04845_ (atpI_hang) | R | CTGGCGTCACCAGGCGCAGTGGATTATATTGCCTTTCCCCAAGAAGCG | 74 |
| Right arm | *atpI* up arm | F | TGCGCCTGGTGACGCCAG | 65 |
|  |  | R | TGCACATGAACTCGAGCATCATTTGCCAAGTAAATAAATATGCTGTGCG | 69 |
| **∆*atpI*::(QT234_17940-45)CM.Gst42III (isoschizomer of M.BamHI)** | | | | |
| Insert | SJEF4-2_M.BamHI (atpI_hang) | F | CGGCCTTTCGCTGGTGTAGAAGAATAGCACCCCTCTCCTTATG | 71 |
|  | SJEF4-2_M.BamHI (atpI hang) | R | CTGGCGTCACCAGGCGCACAATATGGATTAGCTTGTTAACAAACTAAGCCTC | 71 |
| **∆*atpI*::(GK_P8) M.Gka426III (isoschizomer of M.AlwI)** | | | | |
| Insert | P_GK_P8_-GK_P8 (atpI hang) | F | cggcctttcgctggtgtatcaacacatagacacgcgtatgtacg | 71 |
|  | P_GK_P8_-GK_P8 (atpI hang) | R | ctggcgtcaccaggcgcactaaaatccgacaaatagatactcttgaacttgg | 71 |
| **∆*atpI*::( GT3570_17460-65) CM.Gth3570I** | | | | |
| Insert | P_GT3570_17460_-GT3570_17460-65 (atpI hang) | F | cggcctttcgctggtgtatatttgctcacaacgcgaatcaac | 70 |
|  | P_GT3570_17460_-GT3570_17460-65 (atpI hang) | R | ctggcgtcaccaggcgcatccaaaccatagtcgtaacttccgc | 73 |
| **∆*aslB*::(QT235_01945-60) MS.Gst45I** | | | | |
| Left arm | aslB (pTarget hang) |  | AGTCGACCTGCAGAAGCTTTTACGCCTGTGATCACTATGTTTATCCGC | 71 |
|  | P_rplsWT_ (aslB up hang) | R | ATCGCTGAGATCTGCCTTTGCC | 61 |
| rhaR-P_rhaB_ | rhaR (aslB up hang) | F | AAGGCAGATCTCAGCGATTTAATCTTTCTGCGAATTGAGATGACGCCAC | 70 |
|  | P_rhaB_ (QT235_01945 hang) | R | CTCCAGTAATCATTCGAATCACTCCTTATCTAGATACGACCAGTCTAAAAAGCGCCTGAATTC | 69 |
| Insert 1 | QT235_01945 (P_rplsWT_ hang) | F | GATTCGAATGATTACTGGAGAATTAAAGAACAAAGTCGATAAAATATGGG | 64 |
|  | QT235_01950 (linker) | R | GAAATATATTCAGCAAATACCCCTCCCTCGACCAAACCA |  |
| Insert 2 | QT235_01950 (linker) | F | GGGTATTTGCTGAATATATTTCACACTTAACTATAAAAGGCATCTTAGAG | 63 |
|  | QT235_01960 (P_rplsWT__term hang) | R | GAGCTCTTACCTGTTTCTTTGTGAGTAAAGTTGATAAACTCATCGG | 66 |
| terminator | P_rplsWT__ter (QT235_01960 hang) | F | AGAAACAGGTAAGAGCTCAATTTCGCGCCCCGAAA | 68 |
|  | terminator (aslA hang) | R | TCTGACTAAGCCGGCGCTTGATACCTTAGGCTTCGCAAAAAAACG | 72 |
| Right arm | aslA | F | AGCGCCGGCTTAGTCAGATTTAAT | 62 |
|  | aslA (pTarget hang) | R | GCTGCACATGAACTCGATTTACTGGAAAGGGATGATCCAACCG | 69 |
| **∆*aslB*::(QT234_01885-90) MS.Gst42I** | | | | |
| Insert | QT234_01885 (P_rhaB_ hang) | F | CGCTTTTTAGACTGGTCGTAGTTATTTAGTTGCTAGGTTCTTTAACGATGTGAAAACC | 67 |
|  | QT234_01890 (term hang) | R | CGCGAAATTGAGCTCTTACAAACACCCCTCCCTCGACC | 70 |
| **∆*aslB*::(GK1380-1) MS.Gka426I (GK1380-1)** | | | | |
| Insert | GK1380 (P_rhaB_ hang) | F | cgctttttagactggtcgtaatgatggatgaccatgcaggtatgac | 68 |
|  | GK1381 (term hang) | R | cgcgaaattgagctcttagctgcttgcacctcctagtcg | 70 |
| **Plasmid construction for restriction enzyme sensitivity assay** | | | | |
| GGTACCN_4_CNC | pUC19_N_4_(MS.Gst45II) | F | TACGGGTACCGCCTCTCCCCGCGCG | 74 |
|  |  | R | AGAGGCGGTACCCGTATTGGGCGCTCTTCCGCTTCC | 75 |
| GGTACCN_5_CNC | pUC19_N_5_ (MS.Gst45I) | F | TACGGTACCCGCCTCTCCCCGCGCG | 74 |
|  |  | R | AGAGGCGGGTACCGTATTGGGCGCTCTTCCGCTTCC | 75 |
| AAAYN_5_RTCNC | pUC19_N_5__ SspI (MS.Gst42I) | F | gggacgaaatatttcccgactggaaagcggg | 68 |
|  |  | R | cgggaaatatttcgtcccagctgcattaatgaatcggcc | 69 |
| GGTACCN_3_GTAA | pUC19_N_3_ (MS.GK426I) | F | aTTACcgcGGTACCtgagctgataccgctcgccg | 73 |
|  |  | R | tcaGGTACCgcgGTAAtacggttatccacagaatcagggga | 70 |

*Underlined segments denote the hanging regions.

**Table S8. Primers used for the construction of replication-origin variant vectors**

| **Target** | **Primer** | **Sequence (5'-3')** | **T_m_ (℃)** |
| --- | --- | --- | --- |
| template | pG inv R | TTCCGTTCTCCTCCCGTTTCTCC | 62 |
|  | pG inv F | CACACTCGTTTTAAGGCGTCTAACAGGC | 62 |
| Rep2 | pG2_F | CGGGAGGAGAACGGAAAGGGCTTTTCGTGCGTCAGC | 72 |
|  | pG2_R | CGCCTTAAAACGAGTGTGCCCGTTTGTTGAACTACTCTTTAATAAAATAATTTTTCC | 66 |
| Rep3 | pG3_F | CGGGAGGAGAACGGAATTAACCAGCAGCAACCTCCCTTTTTG | 71 |
|  | pG3_R | CGCCTTAAAACGAGTGTGTTAGTTCCTAACCAATTTCTTGAGTTGCTGAACAC | 68 |
| Rep4 | pG4_F | CGGGAGGAGAACGGAAAGCATAGGGAGTCCAGAGGGC | 71 |
|  | pG4_R | CGCCTTAAAACGAGTGTGTGACGGCAGGTCTCCGCCT | 73 |
| Rep5 | pG5_F | CGGGAGGAGAACGGAAGGAGGGGCCCGCGTGGTG | 72 |
|  | pG5_R | CGCCTTAAAACGAGTGTGACAAGAGTGGCTCCTCCCG | 70 |
| Rep6 | pG6_F | CGGGAGGAGAACGGAACGATTTTTTTTATTATGTCTGACGAAAATATATTCGTATGACG | 67 |
|  | pG6_R | CGCCTTAAAACGAGTGTGCGAGGTTTCGCCTGCTGTCG | 72 |
| Rep7 | pG7_F | CGGGAGGAGAACGGAACTGTCACAACGCTGTCACAACG | 71 |
|  | pG7_R | CGCCTTAAAACGAGTGTGCTACTCATTTTCCCCTAATAGCGTATCTATTGCTTGG | 69 |
| Rep8 | pG8_F | AACGGGAGGAGAACGGAAATGTACATAATCACTAGCGAACAATTCACGC | 70 |
|  | pG8_R | GACGCCTTAAAACGAGTGTGTCACACCTCAACGCCCTTAATTCGTG | 71 |

*Underlined segments denote the hanging regions.

**Table S9. Primers used for plasmid copy number determination by qPCR**

| **Target** |  | **Sequence (5’-3’)** | **Length**  **(bp)** | **Tm**  **(°C)** | **G+C (%)** | **Amplicon size (bp)** |
| --- | --- | --- | --- | --- | --- | --- |
| *Rep1*  *(Rep8)* | F | AAGTGCAGAAGCGTCATCCT | 20 | 58.5 | 50 | 175 |
|  | R | GCCGTGAAGTCGTACCAAAT | 20 | 58.5 | 50 |  |
| *Rep2* | F | GCAATTGACGAAACTGCAAA | 20 | 55 | 40 | 217 |
|  | R | CGGCTTTTTCGTCATCATCT | 20 | 56.5 | 45 |  |
| *Rep3* | F | ACGACTAGGAGCGTCTTCCA | 20 | 60 | 55 | 200 |
|  | R | ATCTCTGCAGCGAGCTTCTC | 20 | 60 | 55 |  |
| *Rep4* | F | AAAGGGTATGGGGTCATGGT | 20 | 58.5 | 50 | 216 |
|  | R | TATTCGGACTTAGCGCAACC | 20 | 58.5 | 50 |  |
| *Rep5* | F | CAAGCTGGAAGGTGTTGTGA | 20 | 58.5 | 50 | 168 |
|  | R | TCAACGATTGAACCCCTAGC | 20 | 58.5 | 50 |  |
| *Rep6* | F | GGACAAGGGTGTCGTCTGTT | 20 | 60 | 55 | 169 |
|  | R | CGTCGTCCTTCAGCTTTTTC | 20 | 58.5 | 50 |  |
| *Rep7* | F | AAATTCAACATCCGCTGTCC | 20 | 56.5 | 45 | 182 |
|  | R | AATCCCAATTCATCCACCAA | 20 | 55 | 40 |  |
| *dnaA* | F | CGTACAACCCGCTTTTCATT | 20 | 56.5 | 45 | 214 |
|  | R | TCGATCAGCAGAACGTCAAC | 20 | 58.5 | 50 |  |
| *rpoB* | F | CAGCTGTCGCAGTTTATGGA | 20 | 58.5 | 50 | 194 |
|  | R | GAGTTGATGAGCCCGATGTT | 20 | 58.5 | 50 |  |
| *infB* | F | AAGCAAGGCGAAATGAAAGA | 20 | 55 | 40 | 165 |
|  | R | CGCCAGTGAAATATCCGATT | 20 | 56.5 | 45 |  |

**Table S10. Primers used for expression vector construction in *Geobacillus***

| **Target** | **Primer** | **Sequence (5'-3')** | **T_m_ (℃)** |
| --- | --- | --- | --- |
| template | P_rplsWT__up_R | CCTGCAGGCGTACTCGAGG | 62 |
|  | sfGFP_SD_intact_F | TCTAGATAAGGAGTGATTCGAATGCGTAAAGG | 61 |
| P_ald_ | P_ald__F (P_rplsWT__up_hang) | CGAGTACGCCTGCAGGCGCGAGAAGGGATGCGGAAACG | 76 |
|  | P_ald__R (sfGFP_SD_hang) | CGAATCACTCCTTATCTAGACACATTTCCTCCTTTTTCCTTCCCGCTTC | 69 |
| P_cggR_ | P_cggR__F (P_rplsWT_ _up_hang) | CGAGTACGCCTGCAGGGACGGGGAAGAAATCGGGTATGG | 74 |
|  | P_cgg_R_R (sfGFP_SD_hang) | CGAATCACTCCTTATCTAGACGATCCTTCCCCTTAGCAGAACATAG | 67 |
| P_gap_ | P_gap__F (P_rplsWT_ _up_hang) | CGAGTACGCCTGCAGGGAAAACATCATCGCCCGCCATGC | 75 |
|  | P_gap__R (sfGFP_SD_hang) | CGAATCACTCCTTATCTAGACGTGTTTTCCTCCTTAACTTTGCTGC | 68 |
| P_gdh_ | P_gdh__F (P_rplsWT_ _up_hang) | CGAGTACGCCTGCAGGACTGGATGGTATTTTACTGGCTTACTCACTTG | 71 |
|  | P_gdh__R (sfGFP_SD_hang) | CGAATCACTCCTTATCTAGAATTAAGCCTCCCCATATTTTTTAGAATTTCGTG | 66 |
| P_igG_ | P_igG__F (P_rplsWT_ _up_hang) | CGAGTACGCCTGCAGGTGTTTTTGCACAAAATGTTTGCCAACCA | 72 |
|  | P_igG__R (sfGFP_SD_hang) | CGAATCACTCCTTATCTAGAAAAGCCTAAAATCCCCCTTCATTTTTATGAAACT | 67 |
| P_ldh_ | P_ldh__F (P_rplsWT_ _up_hang) | CGAGTACGCCTGCAGGTTCGTGTTTATTTTGCTTGTGATTGGCGG | 72 |
|  | P_ldh__R (sfGFP_SD_hang) | CGAATCACTCCTTATCTAGATGCATTCATCCCTTTCAATATAATGTGAATACTTTCAC | 66 |
| P_pdhA_ | P_pdh__F (P_rplsWT_ _up_hang) | CGAGTACGCCTGCAGGTCGCCTTTTCACTCCTGTCTCTCGCATC | 74 |
|  | P_pdh__R (sfGFP_SD_hang) | CGAATCACTCCTTATCTAGACTTGTTCACCTCTGCCTTTCATCAAGAAATTTGG | 68 |
| P_pdhC_ | P_pdhC__F (P_rplsWT_ _up_hang) | CGAGTACGCCTGCAGGTGTCGTTCAAGAAGCGCAACGG | 74 |
|  | P_pdh_C_R (sfGFP_SD_hang) | CGAATCACTCCTTATCTAGATGTCTGTCTACCTCCTATCGTTTGC | 67 |
| P_rplK_ | P_rplK2__F (P_rplsWT_ _up_hang) | CGAGTACGCCTGCAGGTGGATAAAGAACTTGCAATCGTTATTGAAAAGTG | 70 |
|  | P_rplK2__R (sfGFP_SD_hang) | CGAATCACTCCTTATCTAGAGAGACACACCTCCTTAAGTCCGTG | 68 |
| P_rplKA_ | P_rplKA__F (P_rplsWT_ _up_hang) | CGAGTACGCCTGCAGGGGCTGGTGTGTTGTCCGGG | 76 |
|  | P_rplKA__R (sfGFP_SD_hang) | CGAATCACTCCTTATCTAGAGTAAAATCCTCCTCAATTGTGGTTTTAGCGG | 67 |

*Underlined segments denote the hanging regions.

**Table S11. Primers used for construction of inducible expression vectors in *Geobacillus***

| **Target**  **element** | **Primer name** | **Sequence (5'-3')** | **Tm**  **(℃)** |
| --- | --- | --- | --- |
| **GKaraR-P_araD_ (derived from *G. kaustophilus* HTA 426)** | | | |
| template | sfGFP inverse F (GKaraD hang) | TGGGAGCCAATGCGTAAAGGCGAAGAGCTG | 70 |
|  | sfGFP inverse R (GKaraC hang) | TGCGCCATCCCTGCAGGCGTACTCG | 70 |
| GKaraC | GKaraC_R  (pG1AK hang) | CTGCAGGGATGGCGCAGCCATGTG | 68 |
|  | GKaraC_F  (GKaraD hang) | CATTTTATTGAGACAACTCCTTTTTCAAATTCCG | 59 |
| GK_P_araD_ | araD up_F  (GKaraC hang) | GGAGTTGTCTCAATAAAATGTATGAGAGATCAAAAAAAGTTATAG | 60 |
|  | araD up_R  (sfGFP hang) | TTTACGCATTGGCTCCCACTCCCTATG | 64 |
| **GKaraC-P_araD_ (derived from *G. kaustophilus* HTA 426)** | | | |
| template | sfGFP inverse F (GKaraD hang) | TGGGAGCCAATGCGTAAAGGCGAAGAGCTG | 70 |
|  | sfGFP inverse R (GKaraR hang) | GAAATTTAACCTGCAGGCGTACTCGAGG | 63 |
| GKaraR | GKaraR_R  (pG1AK hang) | CCTGCAGGTTAAATTTCCTTTGTTGAGTTACGGAC | 64 |
|  | GKaraR_F  (araD up hang) | CTCTCATACATTTTGCGTGTACAAGTTCCCTCATG | 64 |
| GK_P_araD_ | araD up_F  (GKaraR hang) | ACGCAAAATGTATGAGAGATCAAAAAAAGTTATAG | 58 |
|  | araD up_R  (sfGFP hang) | TTTACGCATTGGCTCCCACTCCCTATG | 64 |
| **GSfruR-P_pfkB_ (derived from *G. stearothermophilus* EF60045)** | | | |
| template | sfGFP_intact_F | ATGCGTAAAGGCGAAGAGCTG | 61 |
|  | sfGFP inverse R | CCTCGAGTACGCCTGCAGG | 61 |
| GSFruR-P_pfkB_ | PfruR_F  (pG1AK hang) | CGAGTACGCCTGCAGGGGGAAAAAGGCACTTGCCGATGG | 75 |
|  | PfruR_R (sfGFP_hang) | GCTCTTCGCCTTTACGCATCATGGAGCCACAACCTCTACTGTTG | 71 |
| **GSgalR-P_galK_ (derived from *G. stearothermophilus* EF60045)** | | | |
| template | sfGFP inverse F (GS_P_galK_ hang) | GGCTCTTTCATGCGTAAAGGCGAAGAGCTG | 66 |
|  | sfGFP inverse R (GSgalR hang) | TAGTTCCATCCTGCAGGCGTACTCGAGG | 67 |
| GSgalR | GSgalR_R  (sfGFP hang) | GCCTGCAGGATGGAACTAAAGTATGTGAAAGAGACCACC | 67 |
|  | GSgalR_F  (P_galK_ hang) | CGTTAAAAGGAACATGCCGGGGTGTTTAAGC | 65 |
| GS_P_galK_ | GS_P_galK__F  (galR hang) | CCGGCATGTTCCTTTTAACGAATTATACGGACAAGGAGGGCAT | 69 |
|  | GS_P_galK__R  (sfGFP hang) | CCTTTACGCATGAAAGAGCCTCCTTAAGATCATC | 63 |
| **GSlutR-P_lutP_ (derived from *G. stearothermophilus* EF60045)** | | | |
| template | sfGFP inverse F (GSlutP hang) | GAAAACCGCATGCGTAAAGGCGAAGAGCTG | 67 |
|  | sfGFP inverse R (GSlutR hang) | GGGAAACGACCTGCAGGCGTACTCGAGG | 69 |
| GSlutR-P_lutP_ | GSlutR_R  (sfGFP hang) | GCCTGCAGGTCGTTTCCCTCCTGTTTTTTGAGGATTCAG | 69 |
|  | P_lutP__R (sfGFP hang) | CCTTTACGCATGCGGTTTTCTCCTCCCCTTTATGTCC | 68 |
| **GS_P_acot_ (derived from *G. stearothermophilus* EF60045)** | | | |
| template | sfGFP_SD_intact_F | TCTAGATAAGGAGTGATTCGAATGCGTAAAGG | 61 |
|  | sfGFP inverse R | CCTCGAGTACGCCTGCAGG | 61 |
| GS_P_acot_ | Pacot_F  (pG1AK hang) | CGAGTACGCCTGCAGGTCCATATTGAGTCAACAATAAGAAGGCTTCCTC | 71 |
|  | Pacot_R (sfGFP_SD_hang) | CGAATCACTCCTTATCTAGATATGATCACCTTATATATAACAATACTTTTGAAGCGTACGC | 66 |
| **PTrhaR-P_rhaB_ (derived from *P. thermoglucosidasius* C56-YS93)** | | | |
| template | sfGFP inverse F  (rhaB hang) | AGATAAATCGATGCGTAAAGGCGAAGAGCTG | 64 |
|  | sfGFP inverse R (rhaB hang) | ATTTATTGACCTGCAGGCGTACTCGAGGT | 65 |
| PTrhaR | PTrhaR_F  (pG1AK hang) | GCCTGCAGGTCAATAAATCACCGTAACATTTTGCTCATGA | 67 |
|  | PTrhaR_R  (rhaB up hang) | AAATCCACTTACTACAAATGGCGCAAAAGCC | 63 |
| PT_P_rhaB_ | P_PTrhaB__F  (PTrhaR hang) | CCATTTGTAGTAAGTGGATTTGGAGAAATTAGCAAATAATGC | 62 |
|  | P_PTrhaB__R  (sfGFP hang) | TTTACGCATCGATTTATCTCCTCCTTCAGCTCAT | 64 |
| **PTxylR-P_xylA_ (derived from *P. thermoglucosidasius* C56-YS93)** | | | |
| template | sfGFP_intact_F | ATGCGTAAAGGCGAAGAGCTG | 61 |
|  | sfGFP inverse R (PTxylR hang) | TTTTTGTCACCTGCAGGCGTACTCGAGG | 66 |
| PTxylR | PTxylR_F  (pG1AK hang) | GCCTGCAGGTGACAAAAAGAAAGATGGAAGCCATCC | 68 |
|  | PTxylR_R (PxylA_hang) | CCAGAGCCATATGAAACAGAAACAGTGATTCATTTTTATGTTTGC | 64 |
| P_PTxylA_ | P_PTxylA__F  (xylR hang) | CTGTTTCATATGGCTCTGGGCAAAATAACTAAGCG | 64 |
|  | P_PTxylA__R  (sfGFP hang) | CAGCTCTTCGCCTTTACGCATATCTAACTCCTCCTTAACTTTTAGTAGATTGTCA | 67 |

**Table S12. Primers used for construction of the pG1t-GeoCas9EF genome editing vector**

| **Target**  **element** | **Primer**  **name** | **Sequence (5'-3')** | **Tm (℃)** |
| --- | --- | --- | --- |
| ***oriT* insertion module** | | | |
| *oriT* | oriT_F (pBR322ori_hang) | GCCAACGCGCCCGCCTTTTCCTCAATCGCTC | 73 |
|  | oriT_R (pBR322ori_hang) | CCTCTCCCCAGCTCTTTGGCATCGTCTCTC | 67 |
| template | pBR322ori_down_inv_F (oriT hang) | CAAAGAGCTGGGGAGAGGCGGTTTGC | 66 |
|  | pBR322ori_down_inv_R (oriT hang) | AGGCGGGCGCGTTGGCCGATTCAT | 73 |
| **Cas9 promoter screening** | | | |
| Cas9 promoter | GeoCas9_F  (SD hang) | TCTAGATAAGGAGTGATTCGAATGAGATACAAAATCGGCCTTGATATCGG | 67 |
|  | ColE1_ori_intact_R | TCATGACCAAAATCCCTTAACGTGAG | 59 |
| P_cas_ | P_cas__F (colE1 hang) | CGTTAAGGGATTTTGGTCATGAGGATGCAGCGCTAGGGCAGAC | 71 |
|  | P_cas__R (GeoCas9EF hang) | GCCGATTTTGTATCTCATTTCGATCCCCTCCCATTAATAGAGC | 67 |
| P_gap_ | P_gap__F (colE1 hang) | CGTTAAGGGATTTTGGTCATGAGAAAACATCATCGCCCGCCATGC | 70 |
|  | P_gap__R (SD_hang) | CGAATCACTCCTTATCTAGACGTGTTTTCCTCCTTAACTTTGCTGC | 68 |
| P_gdh_ | P_gdh__F (colE1 hang) | CGTTAAGGGATTTTGGTCATGAACTGGATGGTATTTTACTGGCTTACTCACTTG | 68 |
|  | P_gdh__R (SD_hang) | CGAATCACTCCTTATCTAGAATTAAGCCTCCCCATATTTTTTAGAATTTCGTG | 66 |
| P_ldh_ | p_ldh__F (colE1 hang) | CGTTAAGGGATTTTGGTCATGATTCGTGTTTATTTTGCTTGTGATTGGCGG | 69 |
|  | P_ldh__R (SD_hang) | CGAATCACTCCTTATCTAGATGCATTCATCCCTTTCAATATAATGTGAATACTTTCAC | 66 |
| P_rplK2_ | P_rplK2__F (ColE1 hang) | AAGGGATTTTGGTCATGATGGATAAAGAACTTGCAATCGTTATTGAAAAG | 66 |
|  | P_rplK2__R (SD_hang) | CGAATCACTCCTTATCTAGAGAGACACACCTCCTTAAGTCCGTG | 68 |
| P_rplsWT_ | P_rplSWT__F (colE1 hang) | CGTTAAGGGATTTTGGTCATGAAACAATCGTTAAAGCGGACGTTTTTGCG | 69 |
|  | P_rplSWT__R (SD_hang) | CGAATCACTCCTTATCTAGACAATGCTTTTTCATCATTGCAGCGGAACA | 69 |
| **Cas9 change** | | | |
| template | SD_inv R  (GeoCas9_ hang) | GCCGATTTTGTATCTCATTCGAATCACTCCTTATCTAGA | 62 |
|  | rrnBT1_  (GeoCas_hang)_F | TACAATCAACTCGTGATTGACTCGAGTAAGGATCTCCAGGCATC | 68 |
| GeoCas9 | GeoCas9_F (SD hang) | TCTAGATAAGGAGTGATTCGAATGAGATACAAAATCGGCCTTGATATCGG | 67 |
|  | GeoCas9_intact_R | TCAATCACGAGTTGATTGTAACGGACGG | 63 |
| **Cas9 terminator** | | | |
| template | pG1_colE1_down_F | GATTATCAAAAAGGATCTTCACCTAGATCC | 57 |
|  | GeoCas9_intact_R | TCAATCACGAGTTGATTGTAACGGACGG | 63 |
| rrnBT1-T7Te  terminator | rrnBT1_  (GeoCas9_hang)_F | TACAATCAACTCGTGATTGACTCGAGTAAGGATCTCCAGGCATC | 68 |
|  | rrnBT1_T7Te_  (colE1_down_hang)_R | AGGTGAAGATCCTTTTTGATAATCTTGGTAACGAATCAGACAATTGACGGC | 68 |
| **Recombinase** | | | |
| template | pG-PvuII_inv_R | CTGGCACGACAGGTTTCCCG | 62 |
|  | pG1-term_intact_F | TAAGAGCTCAATTTCGCGCCC | 60 |
| *GKaraR* | GKaraR_F  (pG hang) | GGAAACCTGTCGTGCCAGTTAAATTTCCTTTGTTGAGTTACGGACAATTAGCTCC | 70 |
|  | GKaraR_R  (araD up hang) | CTCTCATACATTTTGCGTGTACAAGTTCCCTCATG | 64 |
| P_araD_ | araD up_F  (GKaraR hang) | ACGCAAAATGTATGAGAGATCAAAAAAAGTTATAG | 58 |
|  | GK P_araD__intact_R | TGGCTCCCACTCCCTATG | 57 |
| *AC* (*bet-exo*) | ACbet_F  (GKParaD hang) | CATAGGGAGTGGGAGCCAATGAACGCGGTCGTCCAATCG | 73 |
|  | ACexo_R  (pG1-term_intact hang) | CGCGAAATTGAGCTCTTATTAGGCGGCATCCGCGAC | 70 |

*Underlined segments denote the hanging regions.

**Table S13. Primers used for the construction of CRISPR editing vectors targeting defense genes**

| **Target**  **element** | **Primer**  **name** | **Sequence**  **(5'-3')** | **Tm (℃)** |
| --- | --- | --- | --- |
| ***oriT* insertion module** | | | |
| *oriT* | oriT_F (pBR322ori_hang) | GCCAACGCGCCCGCCTTTTCCTCAATCGCTC | 73 |
|  | oriT_R (pBR322ori_hang) | CCTCTCCCCAGCTCTTTGGCATCGTCTCTC | 67 |
| template | pBR322ori_down_inv_F (oriT hang) | CAAAGAGCTGGGGAGAGGCGGTTTGC | 66 |
|  | pBR322ori_down_inv_R (oriT hang) | AGGCGGGCGCGTTGGCCGATTCAT | 73 |
| **Homologous arm template amplification** | | | |
| template | pGeoCas9_inverse_F | CTAGGCGCATAGGAACAACTCCTAAATGC | 62 |
|  | pGeoCas9_inverse_R | AGGGCTTTTCGTGCGTCAGC | 63 |
| ∆R.Gst45I (QT235_04845) | Gst45I_UF (pGeoCas9 hang) | TGACGCACGAAAAGCCCTGTGGATTATATTGCCTTTCCCCAAGAAGCG | 72 |
|  | Gst45I_UR (linker) | GTTGTTCATCTGTCCATCCCTCATCTTTGTCTCTTTTGCAA | 65 |
|  | Gst45I_DF (linker) | GGGATGGACAGATGAACAACCGAGAAATTGTCCAAAAGCTG | 68 |
|  | Gst45I_DR (pGeoCas9 hang) | TTGTTCCTATGCGCCTAGTCACCGCTGTCCGCTTGACG | 73 |
| ∆R.Gst45II (QT235_01955) | P_RplsWT__F  (pGtCas9-hang) | TGACGCACGAAAAGCCCTAACAATCGTTAAAGCGGACGTTTTTGC | 71 |
|  | P_rplsWT__R (Gst45II hang) | CTCCAGTAATCATTCGAATCACTCCTTATCTAGACAATGCTTTTTCATC | 65 |
|  | Gst45II_UF  (P_rplsWT_ hang) | GATTCGAATGATTACTGGAGAATTAAAGAACAAAGTCGATAAAATATGGG | 64 |
|  | Gst45II_UR (linker) | GAAATATATTCAGCAAATACCCCTCCCTCGACCAAACCA | 66 |
|  | Gst45II_DF (linker) | GGGTATTTGCTGAATATATTTCACACTTAACTATAAAAGGCATCTTAGAG | 63 |
|  | Gst45II_DR (P_rplsWT__term hang) | GAGCTCTTACCTGTTTCTTTGTGAGTAAAGTTGATAAACTCATCGG | 66 |
|  | P_rplsWT_ _term_F (Gst45II hang) | AGAAACAGGTAAGAGCTCAATTTCGCGCCCCGAAA | 68 |
|  | P_rplsWT_ -term_R  (pGeoCas9- hang) | TTGTTCCTATGCGCCTAGTGATACCTTAGGCTTCGCAAAAAAACG | 69 |
| ∆R.Gst45III (QT235_00960) | Gst45III_UF (pGeoCas9 hang) | TGACGCACGAAAAGCCCTGAAGCGGATGAAACTTGGTTTTATGATGACAC | 71 |
|  | Gst45III_UR (linker) | TTCAGGGGTGGGATTGTCCCCCTCCCTCTTTTGTTTG | 70 |
|  | Gst45III_DF (linker) | GACAATCCCACCCCTGAAAGAAGGGGTGTTTTG | 67 |
|  | Gst45III_DR (pGeoCas9 hang) | TTGTTCCTATGCGCCTAGCTCAACTTGTCGACTTTGCCGACAATC | 71 |
| ∆R.Gst45IV (QT235_10955) | Gst45IV_UF (pGeoCas9 hang) | TGACGCACGAAAAGCCCTCCTTGATACACCCGGGTTCATGAGC | 73 |
|  | Gst45IV _UR (linker) | GGATATATGTGACTCACTTGAATCCGTGATCGTCCATTG | 65 |
|  | Gst45IV _DF (linker) | CAAGTGAGTCACATATATCCCTCCCTCTAGAAATGTTGGTATTGTC | 66 |
|  | Gst45IV _DR (pGeoCas9 hang) | TTGTTCCTATGCGCCTAGGTGTTGCGCCAGCCGGTAAA | 73 |
| ∆R.Gst45V  (QT235_07640) | Gst45V _UF (pGeoCas9 hang) | TGACGCACGAAAAGCCCTCTTCAAGGAGAGCTTGCTGGATCCG | 73 |
|  | Gst45V _UR (linker) | CAACGCCTGCGCTTTCCTATATACACCATTCGTG | 67 |
|  | Gst45V _DF (linker) | AGGAAAGCGCAGGCGTTGCGTCTATGTTCCGA | 70 |
|  | Gst45V _DR (pGeoCas9 hang) | TTGTTCCTATGCGCCTAGCTTTTTGCAGGAAGCTGCTATTTTCCCT | 71 |
| ∆Gst45_Wadjet_III (QT235 07660-75 | Gst45_JetIII_UF (pGeoCas9 hang) | GCTGACGCACGAAAAGCCCTAGCTGCGGTGAACCGTATCA | 74 |
|  | Gst45_JetIII_UR (linker) | CTGGAATCATAAATGCTAAAGACGGGGTTCTTGT | 64 |
|  | Gst45_JetIII_DF (linker) | TTTAGCATTTATGATTCCAGAAAGCCGCC | 62 |
|  | Gst45_JetIII_DR (pGeoCas9 hang) | AGTTGTTCCTATGCGCCTAGGTTCATTGAAGGATATAACACAATG | 66 |
| ∆Gst45_pAgo_Long (QT235_07775) | Gst45_pAgo_L_UF (pGeoCas9 hang) | GCTGACGCACGAAAAGCCCTTATTTGCGCGCCGATTATTCAG | 72 |
|  | Gst45_pAgo_L_UR (linker) | CCCTTTCATACCCTTCTTTGAGTAACGATGGGTC | 64 |
|  | Gst45_pAgo_L_DF (linker) | CAAAGAAGGGTATGAAAGGGTTGCTTGCCC | 65 |
|  | Gst45_pAgo_L_DR (pGeoCas9 hang) | AGTTGTTCCTATGCGCCTAGCATGCGCAAAAACTGTTTTTTC | 69 |
| ∆Gst45_SspBCDE (QT235_11435-60) | Gst45_SspBCDE_UF (pGeoCas9 hang) | GCTGACGCACGAAAAGCCCTGCGTCTCATGCTTGGACA | 74 |
|  | Gst45_SspBCDE_UR (linker) | TAGATTAGAGACATTCGTATAATGGTTGAGTATAG | 57 |
|  | Gst45_SspBCDE_DF (linker) | ATACGAATGTCTCTAATCTAAGTAAACAGAATTC | 57 |
|  | Gst45_SspBCDE_DR (pGeoCas9 hang) | AGTTGTTCCTATGCGCCTAGCCAACTAGGAGAAATAGACATG | 68 |
| ∆Gst45_Gabija (QT235_01955-70) | Gst45_Gabija_UF (pGeoCas9 hang) | GCTGACGCACGAAAAGCCCTcgtaatttaataaataatataggat | 65 |
|  | Gst45_Gabija_UR (linker) | TTGCATTACTCTGTATCCCTCCTCGAACTTTAAC | 63 |
|  | Gst45_Gabija_DF (linker) | AGGGATACAGAGTAATGCAATCCAATTAAAACCTC | 62 |
|  | Gst45_Gabija_DR (pGeoCas9 hang) | AGTTGTTCCTATGCGCCTAGATGAAGAACAAAATATGGTTTTACT | 66 |
| ∆Gst45_AbiD (QT235_03025) | Gst45_AbiD_UF (pGeoCas9 hang) | GCTGACGCACGAAAAGCCCTagcatttggttccctgtacatttccg | 73 |
|  | Gst45_AbiD_UR (linker) | cgccccacaatcactccaaaaaaactcgtagttagctg | 67 |
|  | Gst45_AbiD_DF (linker) | tttggagtgattgtggggcggggtaaaaatttttgactg | 67 |
|  | Gst45_AbiD_DR (pGeoCas9 hang) | AGTTGTTCCTATGCGCCTAGttgttttttcttcgtgagtagaggatgagg | 69 |
| ∆Gst45_Wadjet_II (QT235-01700-20) | Gst45_JetII_UF (pGeoCas9 hang) | GACGCACGAAAAGCCCTgcgagggcaggcagcac | 76 |
|  | Gst45_JetII_UR (linker) | gtgaatgcactccgtccacctttgcttcgg | 68 |
|  | Gst45_JetII_DF (linker) | tggacggagtgcattcacgagctagaggaaggac | 69 |
|  | Gst45_JetII_DR (pGeoCas9 hang) | AGTTGTTCCTATGCGCCTAGcgtggtgttcatcatagctgaagatgg | 71 |
| ∆Gst45_AbiE (QT235_17105-10) | Gst45_AbiE_UF (pGeoCas9 hang) | GCTGACGCACGAAAAGCCCTTAAAAACGTTAAAAGCCGGCGG | 73 |
|  | Gst45_AbiE_UR (linker) | GTGATCATGACGAAGAAGACAACAACGGACAAC | 64 |
|  | Gst45_AbiE_DF (linker) | GTCTTCTTCGTCATGATCACCACTCTTTTCACC | 64 |
|  | Gst45_AbiE_DR (pGeoCas9 hang) | AGTTGTTCCTATGCGCCTAGGCCGTCCCATGACGGAGG | 74 |
| ∆Gst45_CBASS_I (QT235_00950-60) | Gst45_CBASS_I_UF (pGeoCas9 hang) | GCTGACGCACGAAAAGCCCTcatctcgatgcgcggcct | 77 |
|  | Gst45_CBASS_I_UR (linker) | actgtagagaaaattgaaacggaatttcgtctattttttagtaaatttg | 62 |
|  | Gst45_CBASS_I_DF (linker) | ccgtttcaattttctctacagtttgtagaaaaattcataggagatgc | 64 |
|  | Gst45_CBASS_I_DR (pGeoCas9 hang) | AGTTGTTCCTATGCGCCTAGactcttctaatcgcaattctctttcttttccc | 69 |
| ∆R.Gst42I (QT234_01895) | R.Gst42I_UF (pG hang) | TGGCCGGCCCTGACAGCGGCAAGAGGTACAATCGGAAACATC | 75 |
|  | R.Gst42I_UR (linker) | TGTTCTCGTTGGTTGCTTGCACCTCCTAGTCAAAAATAAGTC | 67 |
|  | R.Gst42I_DF (linker) | GCAAGCAACCAACGAGAACAGGATAGCACTTTCTTTTCACG | 69 |
|  | R.Gst42I_DR (pG hang) | ACTTTTCGGGGAAATGTGCTCACCTAGATATAGGCGATTATGAAGACCC | 69 |
| ∆R.Gst42II (QT234_08775) | R.Gst42II_UF (pG hang) | AATGGCCGGCCCTGACAGCTTCAAAGACCCTACTTACGTCTCGCTAG | 75 |
|  | R.Gst42II_UR (linker) | CGCCCATCGCGAATTTCCTTCACATTTATTAATCGTTGAACAACTCTCCC | 69 |
|  | R.Gst42II_DF (linker) | GGAAATTCGCGATGGGCGGCATCAACC | 67 |
|  | R.Gst42II_DR (pG hang) | CGCCCATCGCGAATTTCCTTCACATTTATTAATCGTTGAACAACTCTCCC | 69 |
| ∆Gst42 Wadjet_II  (QT234_01685-700) | Gst42_JetII_UF (pGeoCas9 hang) | GACGCACGAAAAGCCCTgcgagggcaggcagcac | 76 |
|  | Gst42_JetII_UR (linker) | tcgtgaatgcattccgtccacctttgcttcga | 68 |
|  | Gst42_JetII_DF (linker) | tggacggaatgcattcacgagctagaggaaggac | 68 |
|  | Gst42_JetII_DR (pGeoCas9 hang) | AGTTGTTCCTATGCGCCTAGcgtggtgttcatcatagctgaagatgg | 71 |
| ∆Gst42 BREX_III  (QT234_06800-15) | Gst42_BREX_III_UF (pGeoCas9 hang) | GACGCACGAAAAGCCCTcatctttcgctcttctatgcacacg | 71 |
|  | Gst42_BREX_III_UR (linker) | ccatgtatgatcctcaaggatatcctcttttcccctaaattagtg | 64 |
|  | Gst42_BREX_III_DF (linker) | tatccttgaggatcatacatggtaaaaccagtagg | 62 |
|  | Gst42_BREX_III_DR (pGeoCas9 hang) | AGTTGTTCCTATGCGCCTAGcgtatacacccggctttttaaccac | 71 |
| ∆Gst42_CBASS_II (QT234_08520-80) | Gst42_CBASS_II_UF (pGeoCas9 hang) | GACGCACGAAAAGCCCTacgaaacgtttctttctgccaattatcagc | 71 |
|  | Gst42_CBASS_II_UR (linker) | ctaaacatcacaccacgggtaatatgtatgctctttt | 64 |
|  | Gst42_CBASS_II_DF (linker) | ttacccgtggtgtgatgtttagatttgtcagagggttactaatacg | 67 |
|  | Gst42_CBASS_II_DR (pGeoCas9 hang) | AGTTGTTCCTATGCGCCTAGaccttttcaagaaccctgatagatgca | 70 |
| ∆Gst42_SoFic (QT234_03560) | Gst42_SoFic_UF (pGeoCas9 hang) | GACGCACGAAAAGCCCTgctctcattatgatgctgtttatcgttgt | 71 |
|  | Gst42_SoFic_UR (linker) | aggcgcctgtcctcattttcattcccctttacgaacattttag | 68 |
|  | Gst42_SoFic_DF (linker) | gaaaatgaggacaggcgcctgtgttgtgc | 68 |
|  | Gst42_SoFic_DR (pGeoCas9 hang) | AGTTGTTCCTATGCGCCTAGacaccgtacaacttgctttgtgaaacgatcgag | 72 |
| ∆GstT Wadjet_II (QSJ10_01575-90) | GstT_JetII_UF (pGeoCas9 hang) | GACGCACGAAAAGCCCTcgatcgccttcgtttcattcgtgc | 73 |
|  | GstT_JetII_UR (linker) | gcacatctctactccgtccaccttcgcttcg | 67 |
|  | GstT_JetII_DF (linker) | tggacggagtagagatgtgctgtcgcttgtccg | 70 |
|  | GstT_JetII_DR (pGeoCas9 hang) | AGTTGTTCCTATGCGCCTAGaaggagacccgatcgctccg | 73 |
| ∆Gka426_Wadjet_II (GK0317-0320) | Gka426_JetII_UF (pGeoCas9 hang) | GCTGACGCACGAAAAGCCCTacatttctctgtggggaacgtg | 73 |
|  | Gka426_JetII_UR (linker) | cgtggatggactccgtccacctttgcttcgg | 71 |
|  | Gka426_JetII_DF (linker) | ggacggagtccatccacgagccagagaaggag | 70 |
|  | Gka426_JetII_DR (pGeoCas9 hang) | AGTTGTTCCTATGCGCCTAGgcacgattgggggtcgacga | 73 |
| ∆Gth3570 Wadjet II (GT3570_01570-85) | Gth3570_JetII_UF (pGeoCas9 hang) | GCTGACGCACGAAAAGCCCTacatttctctgtggggaacgtg | 73 |
|  | Gth3570_JetII_UR (linker) | cgtggatggactccgtccacctttgcttcgg | 71 |
|  | Gth3570_JetII_DF (linker) | ggacggagtccatccacgagccagagaaggag | 70 |
|  | Gth3570_JetII_DR (pGeoCas9 hang) | AGTTGTTCCTATGCGCCTAGgtcaaccaagcccatccatgatggc | 73 |
| **P*_pta_*-N21 protospacer change** | | | |
| R.Gst45I | sgRNA-Gst45I _inverse_F | CGTCCGTACGTGAAACAGCTGTCATAGTTCCCCTGAGATTATCGC | 70 |
|  | sgRNA-Gst45I _inverse_R | AGCTGTTTCACGTACGGACGGTATAACGGTATCCATTTTAAGAATAATCC | 67 |
| R.Gst45II | sgRNA-Gst45II_inverse_F | TGACGATACGATTGATACGGGTCATAGTTCCCCTGAGATTATCGCTG | 69 |
|  | sgRNA-Gst45II_inverse_R | CCGTATCAATCGTATCGTCAGTATAACGGTATCCATTTTAAGAATAATCC | 64 |
| R.Gst45III | sgRNA-Gst45III_inverse_F | TGGATGAGTGTCATCGTGGGGTCATAGTTCCCCTGAGATTATCGCTG | 71 |
|  | sgRNA-Gst45III_inverse_R | CCCACGATGACACTCATCCAGTATAACGGTATCCATTTTAAGAATAATCC | 66 |
| R.Gst45IV | sgRNA-Gst45IV_inverse_F | TGAAAGGCAGGCAGACATTAGTCATAGTTCCCCTGAGATTATCGC | 69 |
|  | sgRNA-Gst45IV_inverse_R | TAATGTCTGCCTGCCTTTCAGTATAACGGTATCCATTTTAAGAATAATCC | 65 |
| R.Gst45V | sgRNA-Gst45V_inverse_F | CAAGAAACGAGACACGTCCTGTCATAGTTCCCCTGAGATTATCGCTG | 70 |
|  | sgRNA-Gst45V_inverse_R | GGACGTGTCTCGTTTCTTGAGTATAACGGTATCCATTTTAAGAATAATCC | 65 |
| Gst45_Wadjet_III | sgRNA-Gst45_Wadjet_III_inverse_F | cgattaaggaatacaaggcgtgtcatagttcccctgagattatcgc | 68 |
|  | sgRNA-Gst45_Wadjet_III_inverse_R | acgccttgtattccttaatcggtataacggtatccattttaagaataatcc | 65 |
| Gst45_pAgo_Long | sgRNA-Gst45_pAgo_L_inverse_F | cctcgccaactatacggtcaagtcatagttcccctgagattatcgc | 69 |
|  | sgRNA-Gst45_pAgo_L_inverse_R | ttgaccgtatagttggcgagggtataacggtatccattttaagaataatcc | 67 |
| Gst45_SspBCDE | sgRNA-Gst45_ SspBCDE_inverse_F | gtgcagccaactcatcatcctgtcatagttcccctgagattatcgc | 70 |
|  | sgRNA-Gst45_SspBCDE_inverse_R | aggatgatgagttggctgcacgtataacggtatccattttaagaataatcc | 67 |
| Gst45_Gabija | sgRNA-Gst45_Gabija_inverse_F | tgcttcgtaatgactcaagatgtcatagttcccctgagattatcgc | 68 |
|  | sgRNA-Gst45_ Gabija_inverse_R | atcttgagtcattacgaagcagtataacggtatccattttaagaataatcc | 64 |
| Gst45_AbiD | sgRNA-Gst45_AbiD_inverse_F | agcataagcacaaagaacagcgtcatagttcccctgagattatc | 67 |
|  | sgRNA-AbiD_ Gabija_inverse_R | ctgttctttgtgcttatgctgtataacggtatccattttaagaataatcc | 64 |
| Gst45_Wadjet_II (GstT, Gst42, Gka426 andGth3570 compatible) | sgRNA-Gst45_Wadjet_II_inverse_F | aacggcacgtcatttacctcggtcatagttcccctgagattatcgc | 71 |
|  | sgRNA-Gst45_Wadjet_II_inverse_R | gaggtaaatgacgtgccgttgtataacggtatccattttaagaataatcc | 66 |
| Gst45_AbiE | sgRNA-Gst45_AbiE_inverse_F | aataaaaagagaccgccttttgtcatagttcccctgagattatcgc | 67 |
|  | sgRNA-Gst45_AbiE_inverse_R | aaaaggcggtctctttttattgtataacggtatccattttaagaataatcc | 64 |
| Gst45_CBASS_I | sgRNA-Gst45_CBASS_I_inverse_F | tgcgaatttattatgctctccgtcatagttcccctgagattatcgc | 68 |
|  | sgRNA-Gst45_CBASS_I_inverse_R | gagagcataataaattcgcagtataacggtatccattttaagaataatcc | 63 |
| Gst42I (QT234_08775) | sgRNA_Gst42I_inverse_F | GTCAACTGGGGGATCTAAGGGTCATAGTTCCCCTGAAAAGTCAGGG | 70 |
|  | sgRNA_Gst42I_inverse_R | CCTTAGATCCCCCAGTTGACGGAAAATTCCTCCCGCGCCATTTTC | 72 |
| Gst42II (QT234_01895) | sgRNA_Gst42II_inverse_F | CGTCGAAGGAGGAGACAAACGTCATAGTTCCCCTGAAAAGTCAGGG | 71 |
|  | sgRNA_Gst42II_inverse_R | TTTGTCTCCTCCTTCGACGCAAAATTCCTCCCGCGCCATTTTC | 72 |
| ∆pSJEF4-2-1 | sgRNA-rep8_inverse_F | TGATCGTTCAGATCAGCGAAGTCATAGTTCCCCTGAGATTATCGC | 69 |
|  | sgRNA-rep8_inverse_R | TTCGCTGATCTGAACGATCACGTATAACGGTATCCATTTTAAGAATAATCC | 66 |
| ∆pSJEF4-2-2 | sgRNA-rep7_inverse_F | TGGCCGCTCCTAGGGGCTATGGTCATAGTTCCCCTGAGATTATCGC | 73 |
|  | sgRNA-rep7_inverse_R | CCCCTAGGAGCGGCCAGTATAACGGTATCCATTTTAAGAATAATCC | 68 |
| ∆pSJEF4-2-3 | sgRNA-rep5_inverse_F | TGTGTCATAGCGGTGTCCAGGTCATAGTTCCCCTGAGATTATCGC | 71 |
|  | sgRNA-rep5_inverse_R | CTGGACACCGCTATGACACACGTATAACGGTATCCATTTTAAGAATAATCC | 67 |
| ∆Gst42 BREX_III | sgRNA-Gst42 BREX_III_inverse_F | gcgccagcagttattaaagaagtcatagttcccctgagattatcgc | 68 |
|  | sgRNA-Gst42 BREX_III_inverse_R | ttctttaataactgctggcgcgtataacggtatccattttaagaataatcc | 66 |
| ∆Gst42 CBASS_II | sgRNA-Gst42 CBASS_II_inverse_F | gccttacggttcaatggatgcgtcatagttcccctgagattatcgc | 70 |
|  | sgRNA-Gst42 CBASS_II_inverse_R | catccattgaaccgtaaggcgtataacggtatccattttaagaataatcc | 66 |
| ∆Gst42 SoFic | sgRNA-Gst42 SoFic_inverse_F | ggatgaagtgatggagactgagtcatagttcccctgagattatcgc | 68 |
|  | sgRNA-Gst42 SoFic_inverse_R | tcagtctccatcacttcatccgtataacggtatccattttaagaataatcc | 66 |
| ∆pBSO1 (ATCC12980, KCTC3570 compatible) | sgRNA-repT_inverse_F | TCTATGACAATATCTTGCCTGTCATAGTTCCCCTGAGATTATCGC | 67 |
|  | sgRNA-repT_inverse_R | AGGCAAGATATTGTCATAGAGTATAACGGTATCCATTTTAAGAATAATCC | 63 |
| ∆pHTA426 | sgRNA-GK426rep_inverse_R | acgcaaagttgcggcgatccggtcatagttcccctgagattatcgc | 74 |
|  | sgRNA-GK426rep_inverse_R | gatcgccgcaactttgcgttgtataacggtatccattttaagaataatc | 67 |

**Table S14. Primers used for the construction of genome-editing vectors for metabolic engineering in *G. stearothermophilus* SJEF4-2**

| **Target** | **Primer name** | **Sequence (5'-3')** | **Tm (℃)** |
| --- | --- | --- | --- |
| **Homologous arm** | | | |
| template | pG_arm_inv_F | CACATTTCCCCGAAAAGTGCCACC | 63 |
|  | pG_arm_inv_R | CTGTCAGGGCCGGCCATT | 62 |
| ∆*araA* | araA_UF (pG hang) | AATGGCCGGCCCTGACAGCGTTGCTAACGAAACCAGAAGAGATTTACCG | 74 |
|  | araA_UR (linker) | AATTTCGGCATCATTGCTCCTCCCCGTTCAC | 67 |
|  | araA_DF (linker) | GGAGCAATGATGCCGAAATTGCCGGTCGCCC | 70 |
|  | araA_DR (pG hang) | ACTTTTCGGGGAAATGTGCCTCTATTGATTTCCAACTTTTATATAAAACTACTTGATGG | 67 |
| ∆*galKETR* | gal_UF (pG hang) | AATGGCCGGCCCTGACAGGCATCGATCAACTCGTCGCAGACGC | 77 |
|  | gal_UR (linker) | AGAAGCAGAAAAAGCCTCCTTAAGGTCATCTAGTC | 64 |
|  | gal_DF (linker) | AGGAGGCTTTTTCTGCTTCTATCTGAGGCGCCAT | 69 |
|  | gal_DR (pG hang) | ACTTTTCGGGGAAATGTGCGCCATCACGATGTCAAAGAAGAAC | 70 |
| ∆*dgoDKA* | dgo_DR (pG1 hang) | AATGGCCGGCCCTGACAGCAAAGCTGGAACGAGACCTTTGCAAGAG | 75 |
|  | dgo_DF (linker) | CAAGACTGTGATGAGCCCTCCAAGCGTTTGC | 67 |
|  | dgo_UR (linker) | GGGCTCATCACAGTCTTGGGGAGTCCGCG | 70 |
|  | dgo_UF (pG hang) | ACTTTTCGGGGAAATGTGATCAATACTTCCCCTATCGGCATGGAAC | 69 |
| ∆(*fbaB-fsa*) | fsa_UF | TGGCCGGCCCTGACAGGACGACCAGTACGGCATCCG | 77 |
|  | fsa_UR | CGCCTTATTTGGTTACATCCTCCTTAGTTGATCGTCGTC | 66 |
|  | fbaB_DF | GGATGTAACCAAATAAGGCGTTTGCCTCTAAAGGCAAG | 66 |
|  | fbaB_DR | ACTTTTCGGGGAAATGTGCGACGACCGCTTTTTTGAAGCGTC | 72 |
| **P_cggR_-N21 protospacer change** | | | |
| *araA*  (SJEF4-2) | sgRNA_araA_F | GGTTGATGAAAGTCATGGCCGGTCATAGTTCCCCTGAAAAGTCAGGG | 72 |
|  | sgRNA_araA_R | GGCCATGACTTTCATCAACCAAAATTCCTCCCGCGCCATTTTC | 70 |
| *galK*  (SJEF4-2) | sgRNA_galK_F | ATTGTTCGATTGTCATCGCGGTCATAGTTCCCCTGAAAAGTCAGGG | 71 |
|  | sgRNA_galK_R | CGCGATGACAATCGAACAATCGAAAATTCCTCCCGCGCCATTTTC | 71 |
| *dgoA*  (SJEF4-2) | sgRNA_*dgoA*_F | CCGACGGAAATCGTTAAGGCGTCATAGTTCCCCTGAAAAGTCAGGG | 71 |
|  | sgRNA_*dgoA*_R | GCCTTAACGATTTCCGTCGGGGAAAATTCCTCCCGCGCCATTTTC | 72 |
| *fbaB*  (SJEF4-2) | sgRNA_*fbaB*_F | GTGAGGTAATGCGGACGTTTGTCATAGTTCCCCTGAAAAGTCAGGG | 71 |
|  | sgRNA_*fbaB*_R | AAACGTCCGCATTACCTCACAAAAAATTCCTCCCGCGCCATTTTC | 71 |

*Underlined segments denote the hanging regions.

**Table S15. Primers used for the construction of expression vectors for l-AI evolution experiments in *G. stearothermophilus***

| **Target** | **Primer name** | **Sequence (5'-3')** | **Tm (℃)** |
| --- | --- | --- | --- |
| **Promoter insertion (P_igG_)** | | | |
| pG1K-P_igG_ | P_igG__F (PrplS_Up_hang) | cgagtacgcctgcaggtgtttttgcacaaaatgtttgccaacca | 72 |
|  | P_igG__R (SD_hang) | cgaatcactccttatctagaaaagcctaaaatcccccttcatttttatgaaact | 67 |
| **Empty vector control** | | | |
| pG1K empty vector | P_rplS__Up_R | Cctgcaggcgtactcgagg | 61 |
|  | P_igG_-Null_F | tctagataaggagtgattcgatgataagagctcaatttcgcgcccc | 69 |
| **pG1K-P_igG_-TMaraA** | | | |
| Insert (TM_araA) | TM_AraA_F (SD_hang) | tctagataaggagtgattcgaATGATAGATCTCAAGCAGTACGAGTTCTG | 66 |
|  | TM_AraA_R (term_hang) | ggcgcgaaattgagctcttatcaTCTTTTCAAAAGCCCCCAGTAGAGTTCG | 71 |
| pG1K-P_igG_ constructs | P_igG__R (SD_hang) | cgaatcactccttatctagaaaagcctaaaatcccccttcatttttatgaaact | 67 |
|  | Term_F | taagagctcaatttcgcgcccc | 62 |
| **pG1K-P_igG_-GSgatY** | | | |
| Insert (GS_gatY) | GS_gatY_F (SD hang) | tctagataaggagtgattcgaATGTTCAATTGTTTAGTCAACACAAAAGAC | 65 |
|  | GS_gatY_R (term hang) | ggcgcgaaattgagctcttaTCAATAACGATTGACACTCATGCAC | 68 |
| pG1K-P_igG_ constructs | P_igG__R (SD_hang) | cgaatcactccttatctagaaaagcctaaaatcccccttcatttttatgaaact | 67 |
|  | Term_F | taagagctcaatttcgcgcccc | 62 |
| **pG1K-P_igG_-GSgatY-fruK** | | | |
| Insert (GS_fruK) | GS_fruK_F (GS_gatY hang) | tgaaaaaggtgagatggatgattttgacgatcacattaaatccg | 65 |
|  | GS_fruK_R (term hang) | cgcgaaattgagctcttattaccttattgattttccacttaagg | 64 |
| pG1K-P_igG_-GSgatY constructs | GS_gatY_R (GS_fruK hang) | tccatctcacctttttcaataacgattgacactcatgcac | 66 |
|  | Term_F | taagagctcaatttcgcgcccc | 62 |
| **pG1K-P_igG_-TMaraA-P_pdhA_-GSgatY** | | | |
| pG1K-P_igG_-TMaraA  constructs | TM_araA_R (P_pdhA_ hang) | agacaggagtgaaaaggcgatcaTCTTTTCAAAAGCCCCCAGTAGAGTTC | 70 |
|  | Term_F | taagagctcaatttcgcgcccc | 62 |
| P_pdhA_ | P_pdhA__intact_F | tcgccttttcactcctgtctctcg | 63 |
|  | P_pdh__R (SD_hang) | cgaatcactccttatctagacttgttcacctctgcctttcatcaagaaatttgg | 68 |
| Insert (GS_gatY) | GS_gatY_F (SD hang) | tctagataaggagtgattcgaATGTTCAATTGTTTAGTCAACACAAAAGAC | 65 |
|  | GS_gatY_R (term hang) | ggcgcgaaattgagctcttaTCAATAACGATTGACACTCATGCAC | 68 |
| **pG1K-P_igG_-TMaraA-P_pdhA_-GSgatY-fruK** | | | |
| pG1K-P_igG_-TMaraA-P_pdhA_ constructs | GS_gatY_R (pfkB hang) | tccatctcacctttttcaataacgattgacactcatgcac | 66 |
|  | Term_F | taagagctcaatttcgcgcccc | 62 |
| Insert (GS_fruK) | GS_fruK_F (GS_gatY hang) | tgaaaaaggtgagatggatgattttgacgatcacattaaatccg | 65 |
|  | GS_fruK_R (term hang) | cgcgaaattgagctcttattaccttattgattttccacttaagg | 64 |
| **pG1K-P_gdh_-TMaraA-P_pdhA_-GSgatY-fruK** | | | |
| pG1K constructs -TMaraA-P_pdhA_-GSgatY-fruK | P_rplS__UP_R | cctgcaggcgtactcgagg | 63 |
|  | TM_AraA_F (SD_hang) | tctagataaggagtgattcgaATGATAGATCTCAAGCAGTACGAGTTCTG | 66 |
| P_gdh_ | P_gdh__F (P_rplS__UP_hang) | cgagtacgcctgcaggactggatggtattttactggcttactcacttg | 71 |
|  | P_gdh__R (SD_hang) | cgaatcactccttatctagaattaagcctccccatattttttagaatttcgtg | 66 |

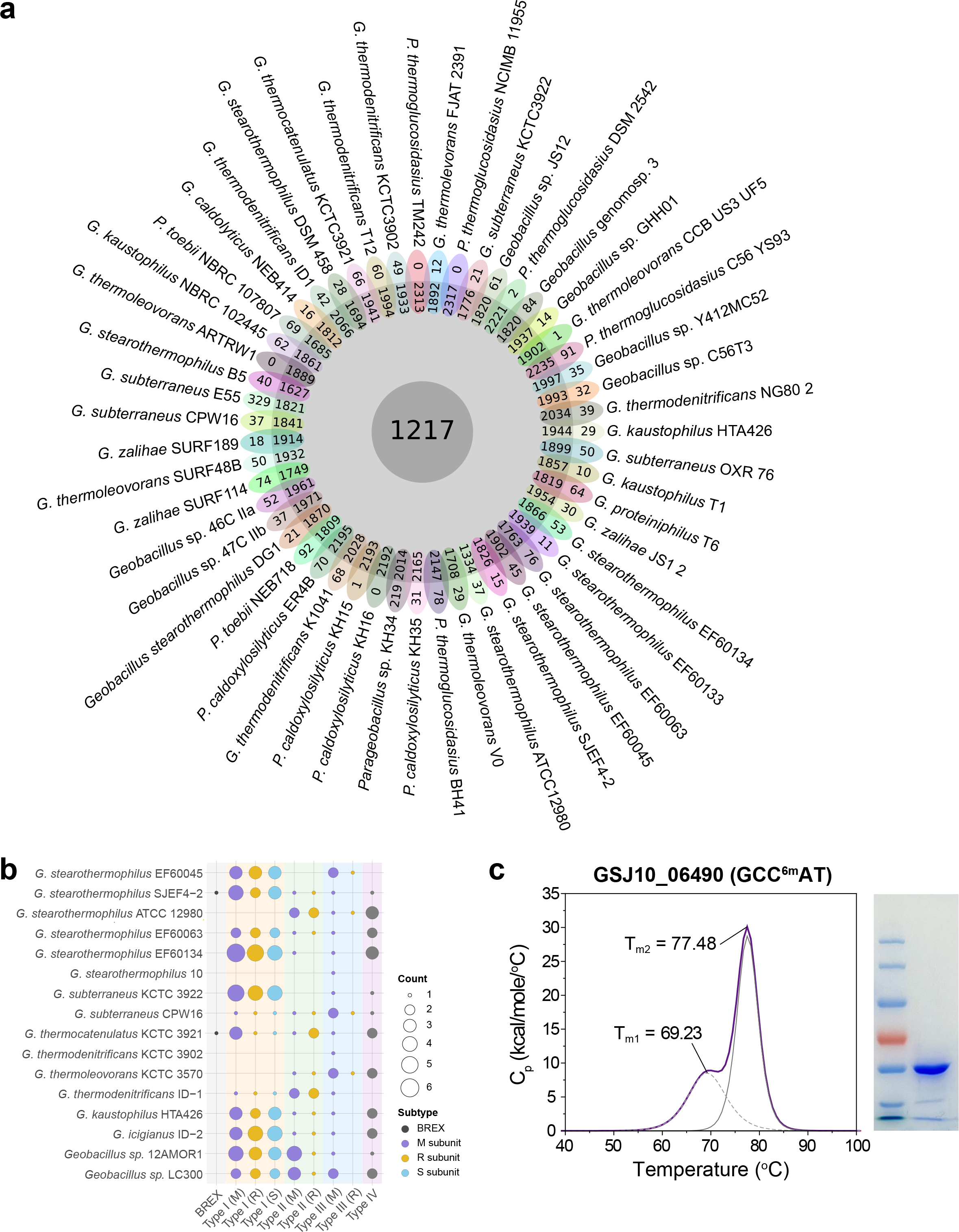

**Figure S1. Comparative analysis of pan-genome structure, R–M systems, and a conserved methyltransferase in (*Para*)*Geobacillus*.** (**a**) Pan-genome structure of 51 *Geobacillus* and *Parageobacillus* strains visualized as a flower plot. The central gray circle represents the 1,217 core genes shared across all strains (≥40% sequence identity). Inner rings denote accessory genes present in multiple strains, whereas outer petals represent strain-specific genes. Each color corresponds to a different strain. Detailed gene lists are provided in **DataSet 3** . (**b**) Distribution of R–M systems across representative strains. Systems are categorized by type (Type I–IV and BREX), with methylation activity inferred from PacBio SMRT sequencing ^13, 14^. Dot size indicates the number of subunits detected, and color denotes system class according to REBASE. (**c**) Thermal stability of the conserved Type III adenine methyltransferase GSJ10_06490 (recognizing GCC^6m^AT), measured by differential scanning calorimetry (DSC). Two melting transitions (T_m1_ and T_m2_) indicate distinct thermally stable structural domains consistent with adaptation to high temperatures. The purified enzyme used for analysis is shown by SDS-PAGE.

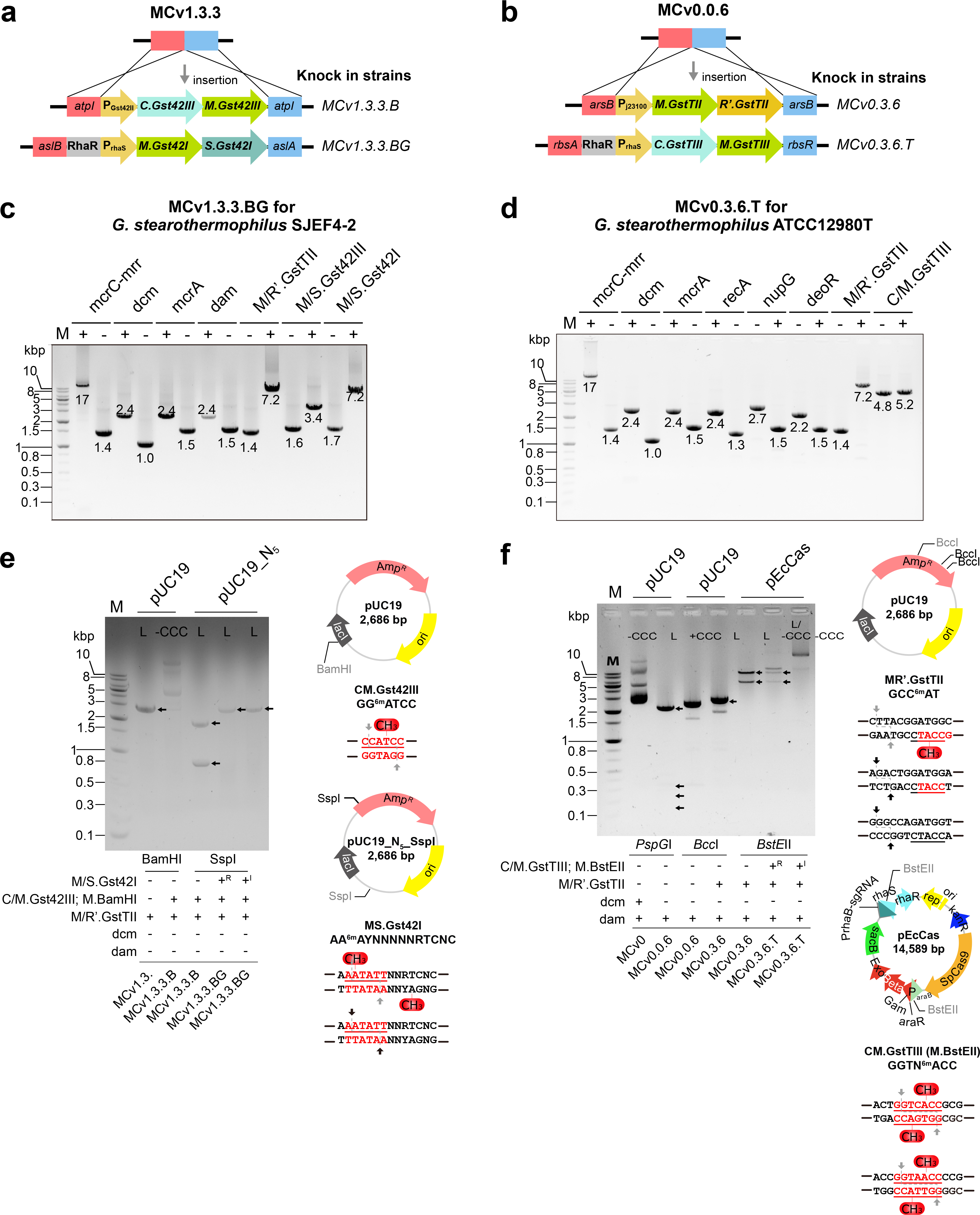

**Figure S2. Construction and methylation validation of strain-specific plasmid artificial modification hosts for *G. stearothermophilus* SJEF4-2 and ATCC12980^T^.** (**a**) Design of the MCv1.3.3.BG plasmid artificial modification host for SJEF4-2. The type II methyltransferase CM.Gst42II, with its native regulatory region, was integrated at the *atpI* locus, and the type I methyltransferase MS.Gst42I, driven by the RhaR-P_rhaS_ system, was inserted between *aslB* and *aslA*. (**b**) Design of the MCv0.6.3T plasmid artificial modification host for ATCC12980^T^. The type III methyltransferase M.GstTII, together with a catalytically inactive restriction subunit (MR’.GstTII) under the synthetic promoter P_J23100,_ was integrated at the *arsB* locus, while the type II methylase CM.GstTIII, controlled by RhaR-P_rhaS_, was inserted between *rbsA* and *rbsR*. (**c**) PCR validation of stepwise plasmid artificial modification host engineering in SJEF4-2. Sequential deletions of Type IV restriction systems (*mcrA*, *mcrC-mrr*) and endogenous methylases were followed by integration of SJEF4-2-derived methylation modules. Expected fragment sizes for deletion (∆) and insertion (ins) events are indicated. (**d**) PCR validation of stepwise plasmid artificial modification host engineering in ATCC12980^T^. Type IV restriction genes (*mcrA*, *mcrC–mrr*), the endogenous dcm methyltransferase, and additional genes affecting DNA stability (*recA*, *nupG*, and *deoR*) were removed prior to insertion of strain-specific methylation cassettes. (**e**) Restriction enzyme protection assay demonstrating functional methylation in MCv1.3.3BG derivatives. Plasmid DNA isolated from engineered hosts showed resistance to enzymes targeting motifs methylated by CM.Gst42III (GG^6m^ATCC) and MS.Gst42I (AA^6m^AYNNNNNRTCNC), resulting in distinct DNA topologies (linear versus covalently closed circular). Plasmid maps of pUC19 and the modified pUC19_N_5__SspI vectors indicate methylation sites and predicted cleavage patterns. (**f**) Restriction protection assay for MCv0.3.6.T derivatives. Methylation by MR’.GstTII (GCC^6m^AT) and CM.GstTIII (GGTN^6m^ACC) altered sensitivity to motif-specific restriction enzymes, confirming in vivo activity of the introduced methylation systems. Plasmid maps of pUC19 and pEcCas highlight recognition motifs and cleavage outcomes.

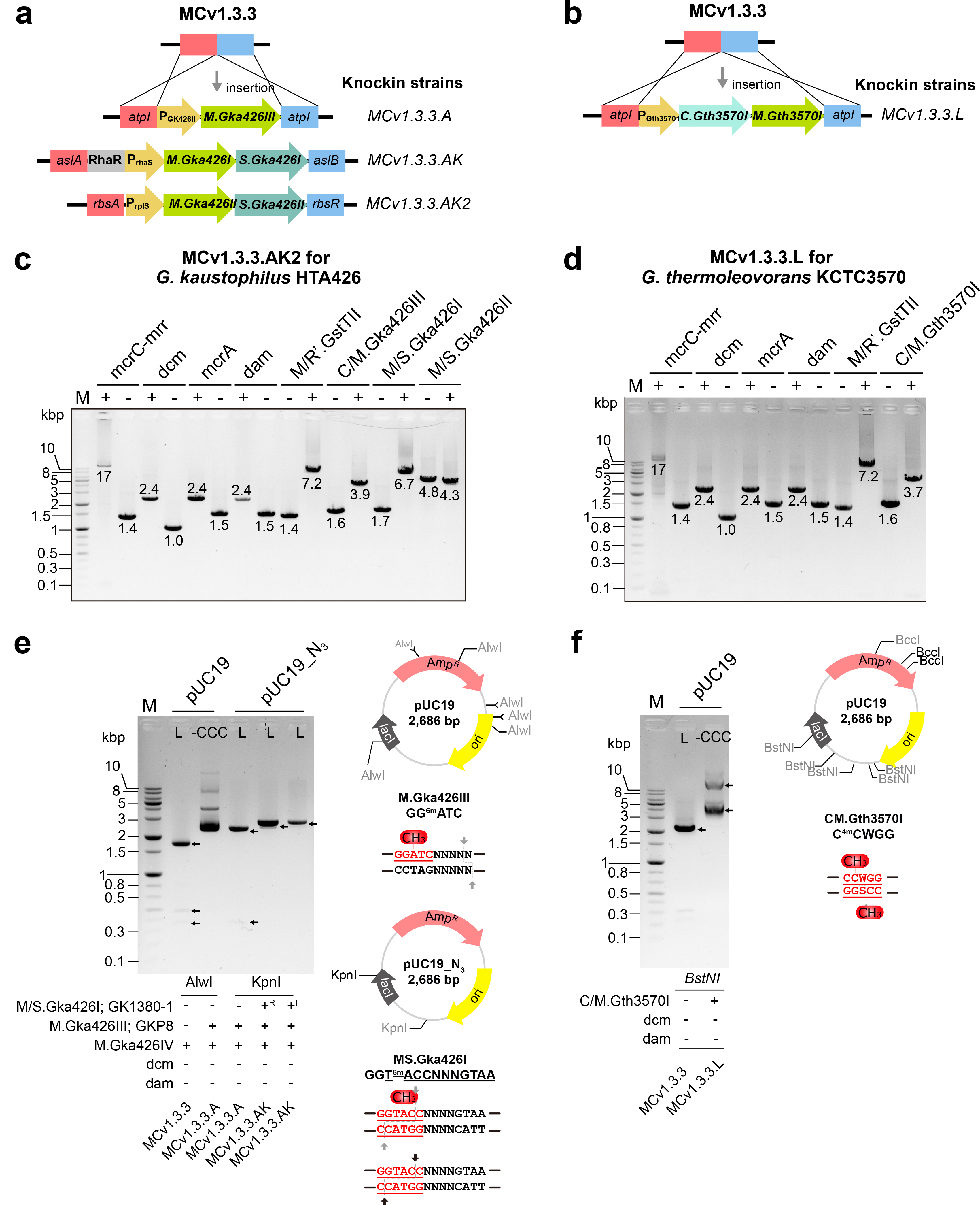

**Figure S3. Construction and methylation validation of strain-specific plasmid artificial modification hosts for *G. kaustophilus* HTA426 and *G. thermoleovorans* KCTC3570^T^.** (**a**) Design of the MCv1.3.3AK2 plasmid artificial modification host for HTA426. The Type II methyltransferase M.Gka426III with its native regulatory region was integrated at the *atpI* locus, while two Type I methyltransferases (MS.Gka426I and MS.Gka426II) under control of the RhaR-P_rhaS_ system were inserted at the *aslA* and *aslB* And *rbsA* –*rbsR* loci, respectively. (**b**) Design of the MCv1.3.3L plasmid artificial modification host for KCTC3570^T^. The Type II methyltransferase CM.Gth3570I with its native upstream region was integrated at the *atpI* locus. (**c**) PCR validation of stepwise plasmid artificial modification host engineering in HTA426. Sequential removal of Type IV restriction systems (*mcrA*, *mcrC-mrr*) and endogenous *dcm* and *dam* methylation modules. Expected fragment sizes for deletion and insertion events are indicated. (**d**) PCR validation of plasmid artificial modification host engineering in KCTC3570^T^, showing analogous removal of endogenous restriction systems and integration of the strain-specific methylation cassette. (**e**) Restriction protection assay demonstrating functional methylation in MCv1.3.3AK2 derivatives. Plasmid DNA isolated from engineered hosts displayed altered sensitivity to enzymes targeting motifs methylated by M.Gka426III (GG^6m^ATC) and MS.Gka426I (Y^6m^ACNNNNGTAA), resulting in protection from cleavage. Plasmid maps of pUC19 and the modified pUC19_N3_KpnI vectors indicate methylation sites. (**f**) Restriction protection assay for MCv1.3.3L derivatives, confirming activity of CM.Gth3570I (C^4m^CWGG) through motif-specific resistance to restriction enzymes. Plasmid maps of pUC19 indicate recognition sequences.

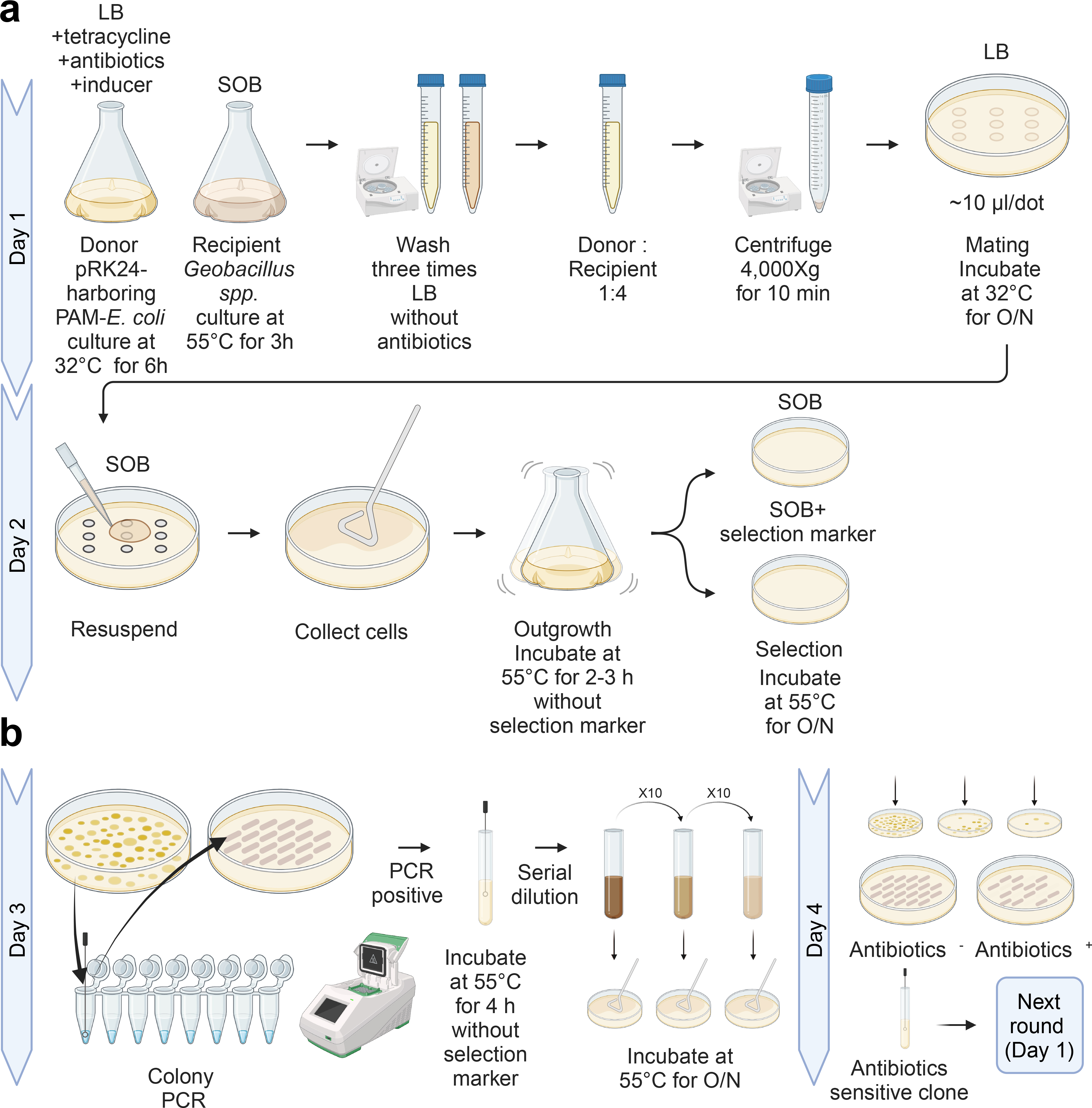

**Figure S4. GeoCas9EF-mediated genome editing workflow in *Geobacillus* via conjugation.** This schematic illustrates the conjugation-based genome editing process using the thermophilic CRISPR nuclease GeoCas9EF. **(a)** Standard conjugation procedure for plasmid transfer from a plasmid artificial modification donor strain to *Geobacillus*. The donor strain is an F-plasmid–containing *E. coli* (pRK24) engineered with an *in vivo* methylation-mimicry system compatible with *Geobacillus*. Donor cells harboring the editing plasmid are cultured at 32°C, while recipient *Geobacillus* cells are grown at 55°C. Cells are washed to remove antibiotics, mixed at a donor-to-recipient ratio of 1:4, concentrated, and spotted onto LB agar for overnight mating at 32°C. The following day, cells are recovered, allowed to outgrow at 55°C without selection, and plated on selective medium to isolate transconjugants. **(b)** GeoCas9EF-mediated genome editing and plasmid curing workflow. Individual colonies are screened by colony PCR to identify successful edits. PCR-positive clones are propagated without antibiotic selection to promote loss of the editing plasmid. Serial dilution and replica plating on media with and without antibiotics enable identification of antibiotic-sensitive clones that have cured the plasmid. These clones can serve as recipients for subsequent editing cycles.

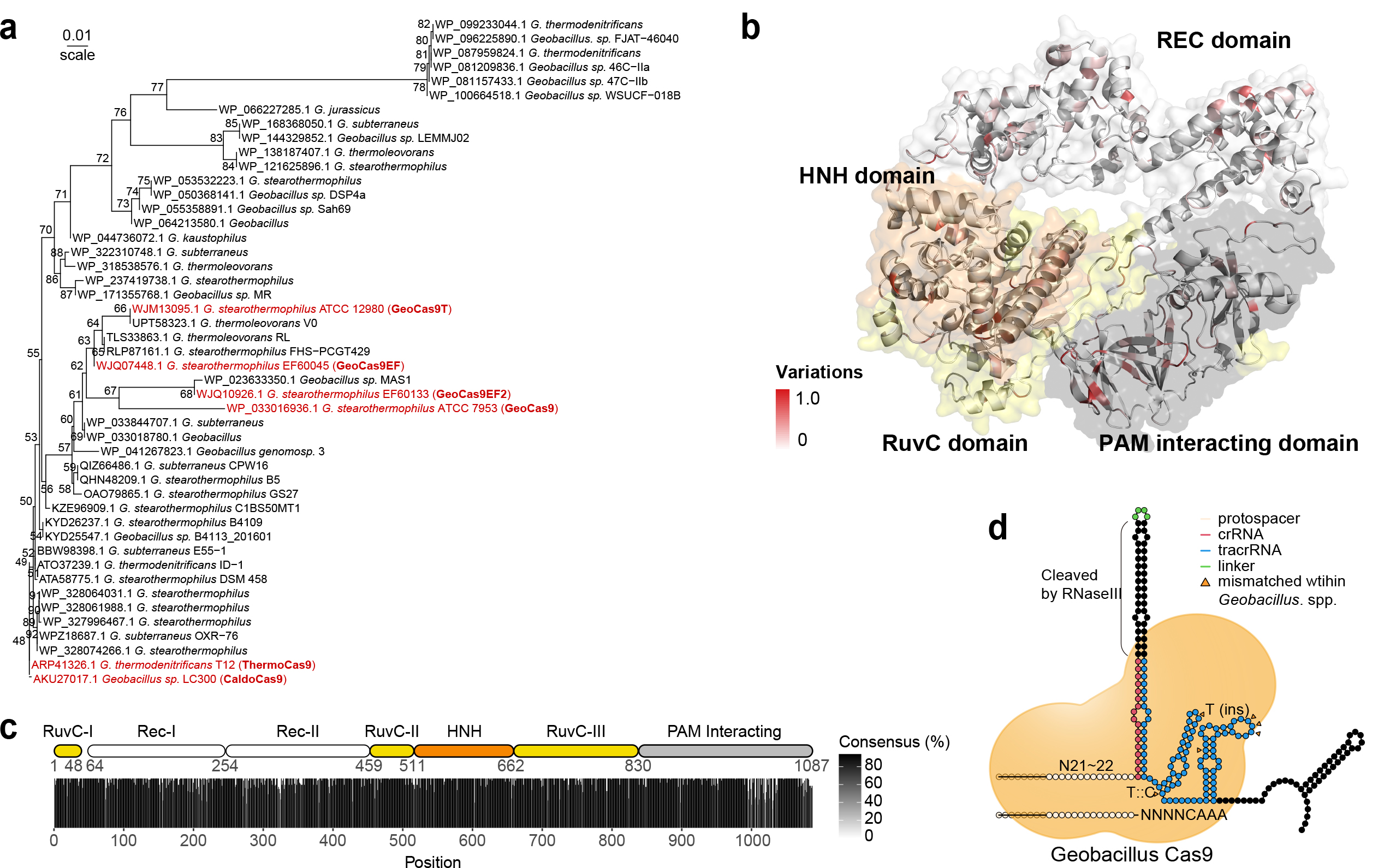

**Figure S5. Phylogenetic tree of thermostable Cas9s derived from *Geobacillus* spp.** (**a**) This phylogenetic tree represents the evolutionary relationships of Cas9/Csn1 proteins derived from *Geobacillus* spp., based on an analysis of 47 non-redundant Cas9 sequences available in the NCBI protein database. The phylogenetic analysis was conducted using IQ-TREE2 version 2.2.3, examining all available models to identify the best fit for my data. The Q.bird+I+G4 model was determined to be the most suitable. To assess the reliability of the inferred phylogenetic tree, a bootstrap analysis was performed 1,000 times, ensuring the robustness of the tree topology presented. (**b**) The 3D structural model of GeoCas9 (WP_033016936.1) was constructed using AlphaFold2. Domains within the model are named according to existing information on GeoCas9, with distinct surface colors utilized for differentiation. Additionally, the color coding of residues indicates the degree of variation based on an alignment of 47 Cas9 sequences. (**c**) The proportion of the most frequently occurring amino acids at each position is based on an analysis of the aligned Cas9 sequences. (**d**) The schematic illustrates *Geobacillus* Cas9's interactions with crRNA, tracrRNA, and a protospacer adjacent motif (PAM), and depicts species-specific sequence mismatches within the scaffold between the GeoCas variants and GtCas9.

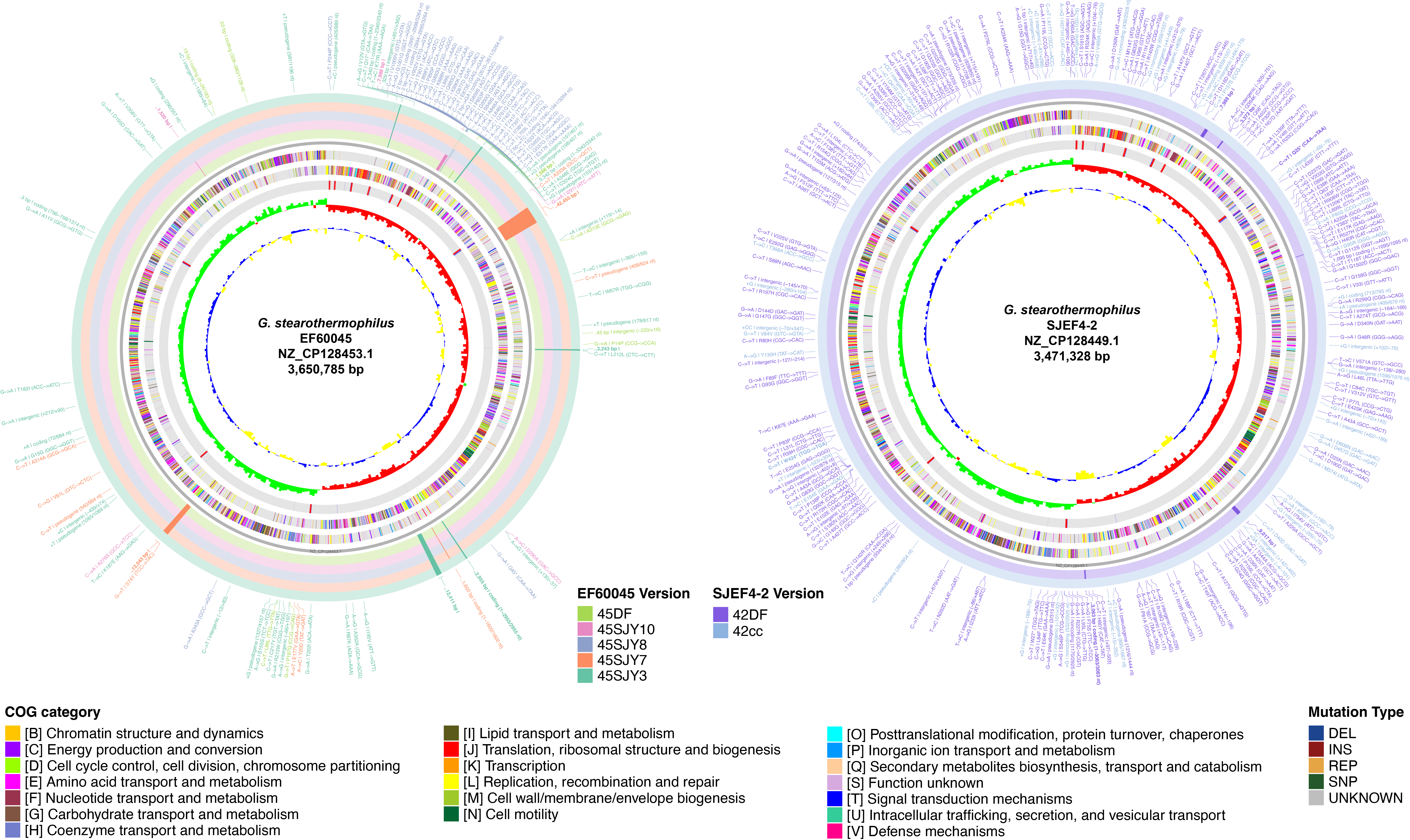

**Figure S6. Genome-wide mutation mapping in *G. stearothermophilus* strains during stepwise elimination of defense systems.** Circular genome maps of *G. stearothermophilus* EF60045 (left) and SJEF4-2 (right) depict mutations accumulated during sequential engineering toward defense-free strains. Mutations were identified relative to the reference genomes EF60045 (NZ_CP128453.1) and SJEF4-2 (NZ_CP128449.1) using breseq ^15^. From inside: The inner track indicates the GC skew wherein sharp blue peaks and dark yellow peaks represent regions with positive and negative values, respectively. The red- and green-colored histograms in the second track represent regions of AT and GC richness, respectively. The multicolored histogram in the third track represents ORFs coded according to functional classification: blue rods, tRNAs; red, rRNAs. The fourth outer circle indicates protease loci in the genome to scale. Outer Each ring represents a different ALE lineage, ordered from the outermost (early) to innermost (late) stage of evolution. Mutation types are categorized as deletions (DEL), insertions (INS), replacements (REP), single nucleotide polymorphisms (SNP), and unknown (UNKNOWN). The visualization was generated using the R packages circlize ^16^ and breseqConverter.

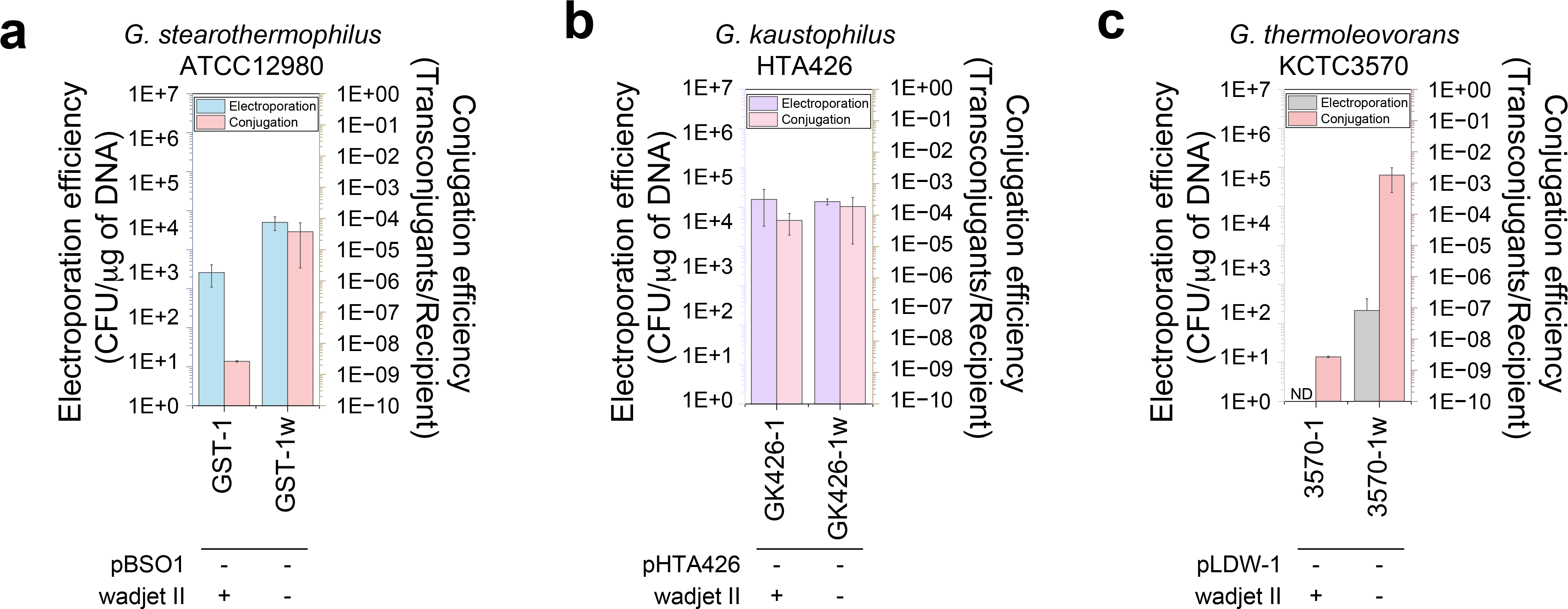

**Figure S7. Effects of defense system removal on DNA transfer efficiencies in diverse *Geobacillus* strains.** (**a-c**) Electroporation and conjugation efficiencies of representative strains—*G. stearothermophilus* ATCC 12980ᵀ (**a**), *G. kaustophilus* HTA426 (**b**), and *G. thermoleovorans* KCTC 3570ᵀ (**c**)—following sequential deletion of endogenous defense genes and/or native plasmids. Transformation efficiencies were quantified as colony-forming units per microgram of DNA (CFU µg⁻¹) for electroporation (left y-axis), and transconjugants per recipient cell for conjugation (right y-axis). Bars represent mean values with standard deviation from independent experiments. Removal of the native plasmid and the Wadjet II defense system resulted in stepwise increases in DNA uptake efficiency, although the magnitude of improvement varied among species and transfer methods. ND indicates not detected.

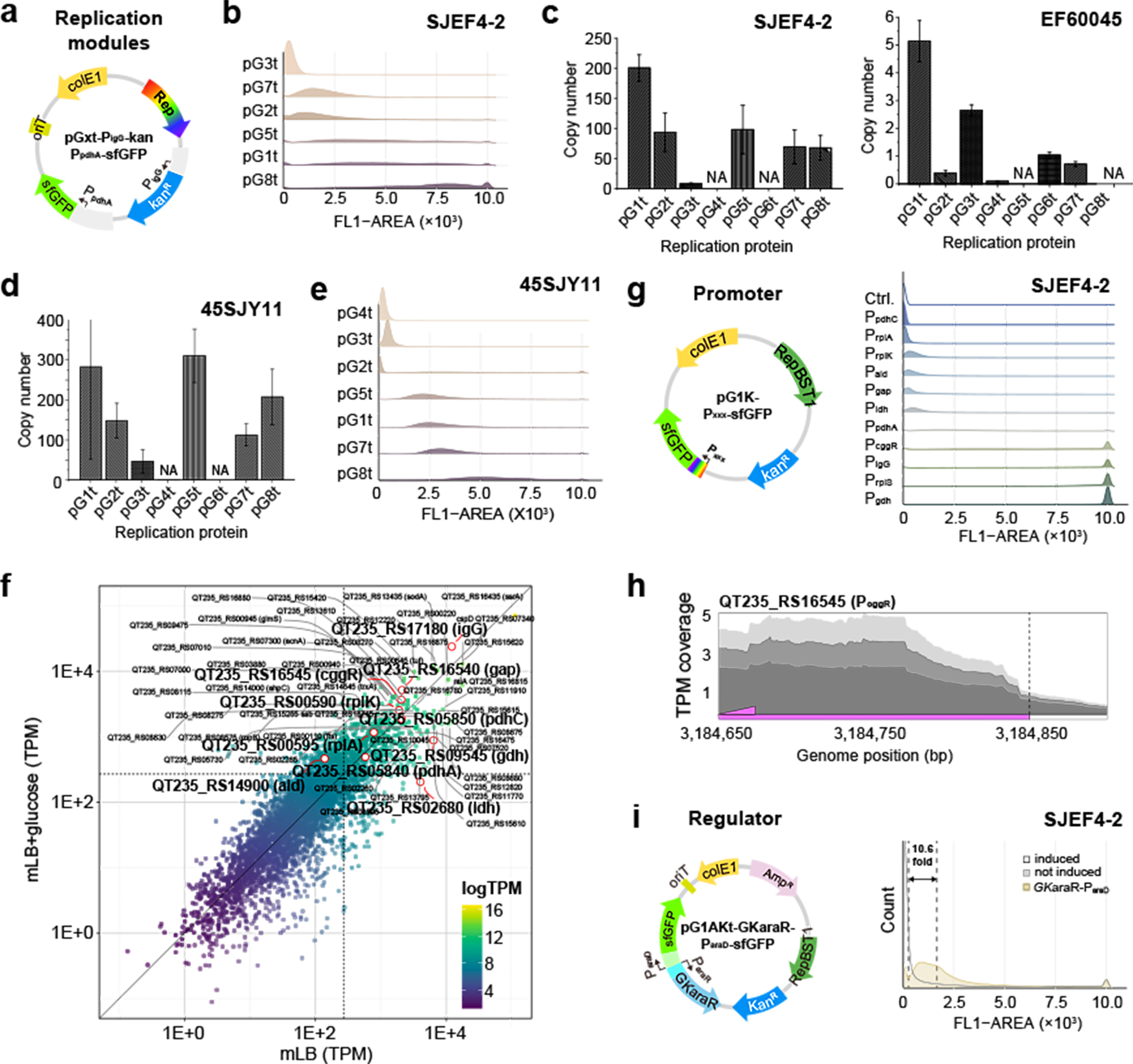

**Figure S8. Modular thermophilic genetic toolkit for programmable replication, expression, and regulation in *Geobacillus.*** (**a**) Replication-module architecture. Shuttle vector (pGxt-P_igG_-kan-P_pdhA_-sfGFP plasmid) carrying a swappable replication protein (Rep) and sfGFP reporter used to evaluate plasmid copy number and expression. (**b**) Flow cytometry histograms (FL1-AREA) of sfGFP fluorescence in *G. stearothermophilus* SJEF4-2 harboring vectors with eight candidate replication proteins (pG1t–pG8t). (**c**) qPCR-based plasmid copy number determination for the same constructs in SJEF4-2 (left) and EF60045 (right). NA indicates replication modules that failed to transform or be stably maintained. Error bars represent the standard deviation (s.d.) of biological replicates. (**d**) Plasmid copy numbers measured in a defense-free strain (45SJY11), showing increased copy number relative to parental backgrounds. (**e**) Corresponding flow cytometry distributions of sfGFP expression in 45SJY11 carrying replication-module variants, demonstrating a positive relationship between copy number and reporter output. (**f**) Transcriptome comparison of EF60045 grown in mLB medium with and without glucose supplementation. Each point represents a gene plotted by expression level (TPM), highlighting candidates with high and stable transcription selected for promoter library construction. (**g**) Promoter library evaluation in SJEF4-2 using the pG1K-P_xxx_-sfGFP reporters. Flow cytometry histograms quantify relative promoter strengths across candidate promoters. (**h**) RNA-seq read coverage across the P_cggR_ locus, illustrating transcriptional architecture and promoter activity.(**i**) Performance of an l-arabinose-inducible regulatory system (GK*araR*-P_araD_) in SJEF4-2, showing sfGFP expression with and without induction.

.
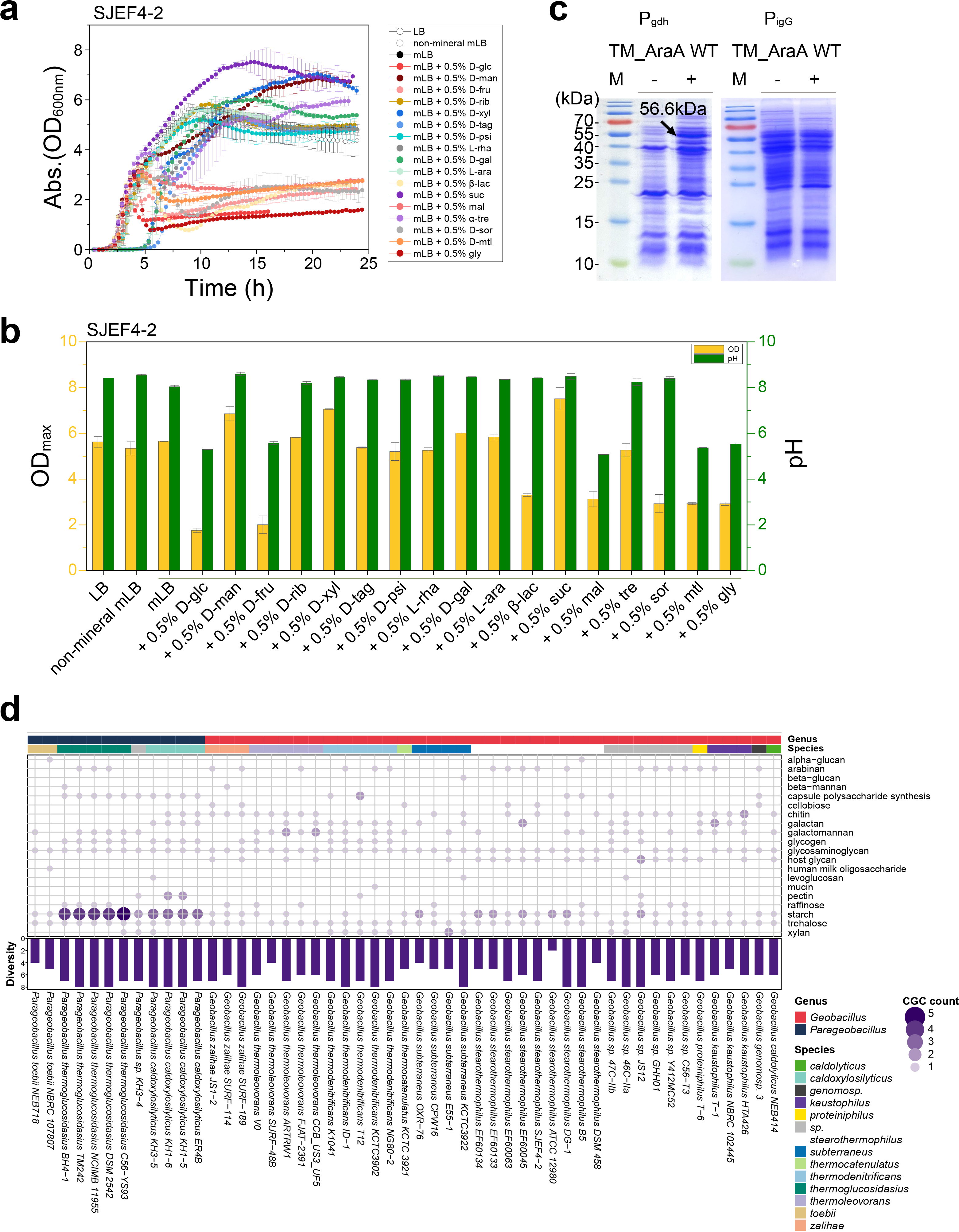

**Figure S9. Broad sugar utilization profile of *G. stearothermophilus*.** (**a**) SDS-PAGE analysis showing expression of l-AI from *T. maritima* MSB8 driven by the P*_igG_* promoter in *G. stearothermophilus* SJEF4-2. A 12% SDS-PAGE gel confirmed a protein band corresponding to the predicted molecular weight of 56.6 kDa. (**b**) Growth curves of SJEF4-2 in mLB medium supplemented with 0.5% (w/v) of individual sugars: d-glc; d-glucose, d-man; d-mannose, d-fru; d-fructose, d-rib; d-ribose, d-xyl; d-xylose, d-tag; d-tagatose, d-psi; d-psicose, d-rha; d-rhamnose, d-gal; d-galactose, l-ara; l-arabinose, b-lac; beta-lactose, suc; sucrose, mal; maltose, a-tre; alpha-trehalose, d-sor; d-sorbitol, d-mtl; d-mannitol, gly; glycerol). Non-mineral mLB is defined as mLB medium without the addition of four minerals: NTA, FeSO_4_, MgSO_4_, and CaCl_2_. Growth was measured at 55°C and 2,000 rpm using a Biosan RTS-8 device. (**c**) Final optical density (OD₆₀₀) and pH values after 24 h of cultivation. (**d**) Distribution of carbohydrate utilization gene clusters (CGCs) across (*Para*)*Geobacillus* genomes. Each row represents a distinct CGC type, and each column represents an individual strain. Dots indicate the presence of a given CGC in a strain. Purple bars (bottom) denote the total number of CGCs identified per strain. All growth measurements and pH determinations were performed in biological triplicate.
